# Mapping Selective Oxidations of Unspecific Peroxygenases

**DOI:** 10.1101/2024.09.10.612301

**Authors:** Dominik Homann, Pascal Püllmann, Martin Weissenborn

## Abstract

Several unspecific peroxygenases (UPOs) have been identified that perform a broad range of selective oxyfunctionalizations and hence represent a pivotal addition to the biocatalysis ‘toolbox’. To make these ‘oxidation tools’ broadly applicable it is crucial to provide a detailed ‘user manual’ for their substrate preference, chemo- and regioselectivity. We therefore selected 16 different substrates with a panel of 15 diverse UPOs and mapped their preferences. Various UPOs proved to be highly selective — discriminating based on either position or chemical properties of the substrate — with up to 99 % chemo- and regioselectivity while achieving turnover numbers (TONs) of a few hundred up to multiple thousands. This map of UPO selectivity shall serve as a starting point for new chemoenzymatic routes and starting points for protein engineering endeavors..

## Introduction

Unspecific peroxygenases (UPOs) perform versatile oxyfunctionalizations similar to P450 enzymes, while only consuming hydrogen peroxide as sole oxygen and electron source.^1-3^ A broad substrate scope and excellent turnover numbers (TONs) render UPOs a highly promising and sought-after enzyme class for future applications.^4^ Major bottlenecks could be overcome in recent years that hindered their broader implementation in academic and industrial research. Amongst them was the UPO production in standard hosts, limiting their widespread use and thus complicating engineering campaigns. Various research groups have contributed to addressing this limitation by various approaches like promoter and signal peptide shuffling.^5-9^ In a recent study, the combination of AlphaFold2 and the PROSS algorithm together with a signal peptide shuffling approach enabled the production of novel UPOs otherwise inaccessible UPOs.^10^ The development of new production systems led to the discovery of a variety of highly active UPOs. Understanding the reactivity patterns of these enzymes is crucial for future applications. To address this need, a set of well-characterized UPO enzymes is required. This understanding would enable researchers from academia and industry to make informed decisions about which UPO can catalyze specific reactions of interest. For protein engineering efforts this information set would be of high value as it substantially reduces the engineering effort by using suitable starting points, i.e. UPOs that already carry the desired property inherently. A map detailing the reactivity and selectivity for a set of UPOs on a broad substrate scope is thus of utmost importance.

To achieve this map, we applied a GC-MS-based fingerprinting method to a diverse set of UPOs (Figure 1).^11^ Three pools with each of four substrates were tested with 15 different enzymes (enzymes were not pooled) resulting in 36 different products. The different substrate pools were created with insights into aromatic, aliphatic, or olefinic oxidations and their chemo- and regioselectivities. The highlights from each pooling experiment in terms of selectivity or activity were analyzed in single substrate experiments. We found regioselective conversions for sp^3^-hybridized carbons in alkane chains, enabling the selective oxidation of different positions. The oxidation of a non-activated sp^3^-hybridized carbon adjacent to an activated benzylic carbon was observed, showcasing a very uncommon reaction for UPOs. Selective epoxidation instead of allylic oxidation and vice versa was found, allowing the targeted oxidation based on the chemical properties.

**Figure 1.**
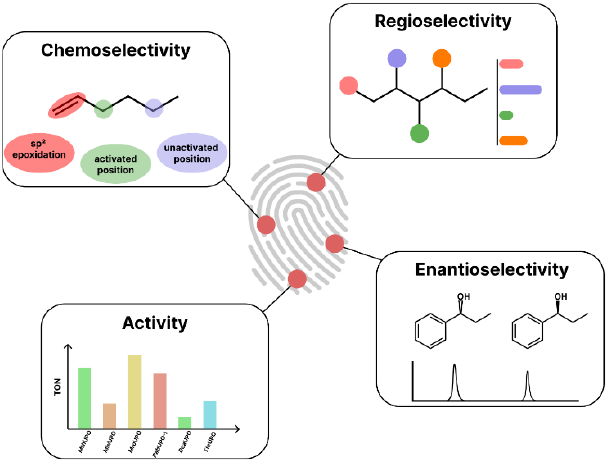
Identifying the substrate spectra of 15 UPOs using the fingerprinting method.

## Results and Discussion

### Initial testing and optimization

15 different UPOs were studied in this work, eight short-type and seven long-type UPOs, respectively. Short-type UPOs typically have a mean weight of 30 kDa, while long-type UPOs exhibit weights around 44.4 kDa.^8^ *Chimera*UPOs have been constructed by Golden Gate shuffling from three different parental UPOs, containing different parts of the *Aae*UPO*, *Gma*UPO, and *Cci*UPO (Figure S2).^5^ Two ancestral UPOs (*Anc*UPOs) were constructed based on *Mth*UPO as a template utilizing the FIREPROT ASR algorithm.^6, 12^

To study and verify the pooling approach, an alkane pool — consisting of cyclohexane, hexane, octane, and decane — was studied with *Chimera*UPO-II. This UPO revealed in the pool a regioselective oxidation at the 2-position of hexane. To verify that the measured selectivities can be reproduced in single analytical runs, this reaction was repeated with only hexane. To our delight, the results from the initial pooling experiment could be reproduced (Figure S1).

In the next step, we investigated the optimal reaction conditions to enable the comparison between different substrates and enzymes. Enzyme concentrations were set to 300 nM for all enzymes and every product was assigned to two *m/z* values (see SI for more information). These *m/z* values help identify the products and enable direct comparison between the enzymes. Every product was analyzed with GC-MS at a fixed concentration (500 µM) and with a specific *m/z* value. The ionization of every product (at the same concentration) is different, which leads to different peak areas. This problem was circumvented by normalizing the obtained peak areas and thus enabling the comparison of all products inside one substrate pool (Figure S44-S46).

### Fingerprinting of 15 UPOs with an alkane substrate pool

The alkane pool was tested with all 15 UPOs individually. The substrates were selected to offer numerous potential positions for the oxidation of non-activated sp^3^-hybridized C-H bonds (Figure 2). These reactions are of particular interest, as they are energetically and stereochemically challenging for classical organic synthetic chemists.^13-15^

**Figure 2.**
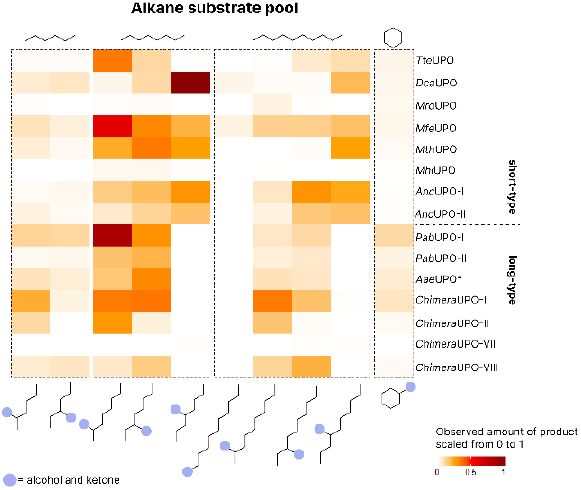
Heatmap of the conversion of the alkane substrate pool, displaying the activities of 15 different UPOs. Reaction conditions: Final volume 400 µl, potassium phosphate buffer (100 mM, pH = 7), 5 v/v % acetone, substrate — (4 mM), hydrogen peroxide — (1 mM), 25 °C, 400 rpm, 1 hour. Concentration was determined by GC-MS.

Each of the 15 UPOs was incubated with the four different substrates leading to a broad distribution of 20 different products and their relative quantities. To focus attention, particularly on the different regioselectivities, overoxidation products to the carbonyl functions were combined with the corresponding alcohol product in the heatmap illustration in Figure 2. Furthermore, overoxidation values are highly setup-specific as they are dependent on factors such as hydrogen peroxide concentration and dosing as well as the overall reaction time. The highest detected product formation was set to the relative activity of 1 and no product formation to 0. This representation gives a “fingerprint” of all UPOs with all the different substrate preferences and regioselectivities in one glance. This fingerprinting further enables the identification of the most selective enzyme for a given product and shows the preference of each UPO with each substrate/product.

### Detailed analysis of most promising UPO/substrate pairs from the alkane substrate pool

The regioselective ratios obtained from the primary fingerprint screening were all reproducible in the single substrate setup. *Dca*UPO showed a high regioselective ratio of 89:11 towards the 4-position of octane in the screening, which could be verified in the single substrate setup, achieving a regioselective ratio of 93:7 (Figure 2, Entry 2d). *Dca*UPO was thus able to outperform the previously most selective UPO *Aae*UPO with a reported regioselective ratio towards hexane of 55:45.^16^ Enantiomeric excess was determined to be 25 %, slightly favoring the (R)*-*enantiomer of the alcohol. 3-octanol was produced by *Aae*UPO* with a low regioselective ratio of 54:46 and a TON of 75 (Entry 2c). The oxidation of octane to 2-octanol was observed for the enzyme *Chimera*UPO-II, achieving an excellent regioselective ratio of 99:1 (Entry 2b). A TON of 724 and an enantiomeric excess of 6 %, favoring the (R)*-*enantiomer, was observed. While *Tte*UPO produced 2-octanol in lower overall amounts, achieving a TON of 512 and a regioselectivity ratio of 81:19, respectively, an enantiomeric excess of 54 of the (R)*-*enantiomer, was measured (Entry 2a).

No terminal oxidation could be observed when using octane as substrate. The oxidation of decane could be catalyzed by all UPOs tested, albeit with overall lower activities compared to octane. The oxidation of decane to 2-decanol by *Chimera*UPO-II resulted in a regioselective ratio of 99:1 (Entry 3b), performing even slightly better than *Mro*UPO, reaching a ratio of 91:9 (Entry 3a). TONs, however were comparably low, accumulating to 32 for *Mro*UPO and 85 for *Chimera*UPO-II. The enantiomeric excess was found to be 23 % and 27 % for *Mro*UPO and *Chimera*UPO-II. *Chimera*UPO-VIII converted decane to 3-decanol with a moderate selectivity of 65:35, whilst achieving a very low TON of 4 (Entry 3c). Finally, the oxidation of decane to 4-decanol was performed by *Mth*UPO with a regioselective ratio of 78:22 and a TON value of 284 (Entry 3e). *Dca*UPO preferred once again the oxidation of the 4-position, as seen before with octane, but reaches a lower regioselective ratio of 85:15 accompanied by a TON of 284 (Entry 3d). While the oxidation of octane and decane could be performed with regioselective ratios of above 70:30 (except 3-octanol), the oxidation of hexane appeared to be more challenging at the 3-position among the tested UPOs. Here, no selective conversion was observed, though the highly selective conversion of hexane to 2-hexanol by *Chimera*UPO-II was observed (Entry 1b). For this reaction a regioselective ratio of 96:4 was achieved, accompanied by a high TON of 3787 and a promising enantiomeric excess of 32 %, favoring the (R)*-*enantiomer. *Mro*UPO was able to reach an even higher regioselective ratio of 99:1, albeit with low activity (44 TON) (Entry 1a). Remarkably, *Mro*UPO slightly favors the (S)-enantiomer, with an enantiomeric excess of 7 %. *Mro*UPO has been reported to oxidize terminal carbons towards the carboxylic acid.^17^ These products could have been formed, but the utilized screening method would not allow to detect them, as the applied extraction was not suitable for isolating carboxylic acids. Screening 15 UPOs towards the alkane substrate pool revealed a UPO that exhibits high selectivity for nearly every tested conversion, accounting for more than 70 percent of product at only one specific position, in some cases even reaching regioselectivities above 90 percent. This was accompanied by already decent non-optimized catalytic activities, ranging from 44 to approximately 4000 TONs. The enantiomeric excess remained low with approximately 20 % to 30 %, rarely exceeding these values. Whilst the primary screening hinted towards *Dca*UPO as the most active enzyme, this could not be verified. Within the secondary screening reactions *Dca*UPO produced 4-octanol with a TON of 163, which is substantially lower than e.g., the formation of 2-hexanol by *Chimera*UPO-II, with a TON of 3787. Possible reasons for this shift are the enzyme concentration, substrate competition, and generally the reaction conditions.

### Fingerprinting of 15 UPOs with an aromatic substrate pool

This pool contained four different substrates with different degrees of aromatic stability — benzene is more difficult to oxidize than naphthalene — and offering benzylic carbons with varying binding energies — toluene’s benzylic group is more stable than phenylpropane’s. Also, the different preference for aromatic oxidation, i.e. via an initial epoxide formation mechanism, versus a benzylic or even aliphatic oxidation is of interest in this pool.^18^ The benzylic position is readily oxidized by most UPOs, so particular attention was paid to oxidations that diverge from this position. The screening was performed as described above and the four aromatic substrates could be converted to 12 different products by the 15 enzymes.

### Detailed analysis of most promising UPO/substrate pairs from the aromatic substrate pool

Table 2 shows the results of single substrate reactions, which were used to give detailed numbers and selectivities to the fingerprinting results.

**Table 1:**
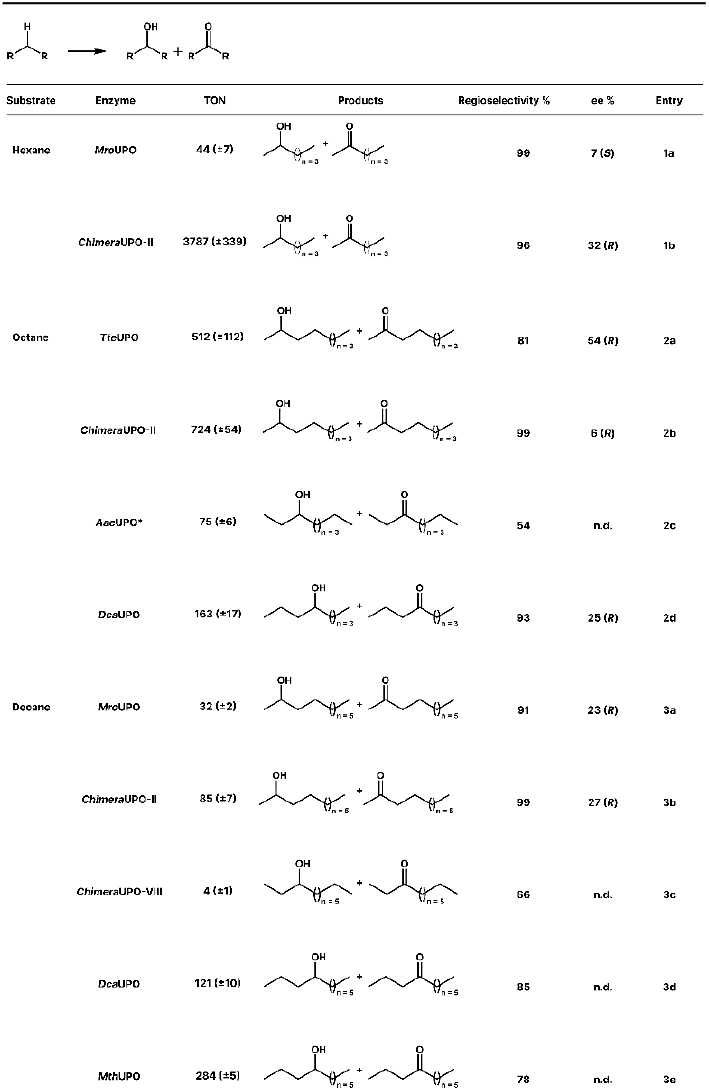
The most selective conversions from the screening of the alkane substrate pool. Reaction conditions: Final volume 500 µl, potassium phosphate buffer (100 mM, pH = 7), 5 v/v % acetone, substrate — (4 mM), hydrogen peroxide — (1 mM), 25 °C, 400 rpm, 1 hour. Concentration was determined by GC-MS. Concentration was determined by GC-MS. Chirality measurements were performed with a chiral column on a GC-MS.

**Table 2:**
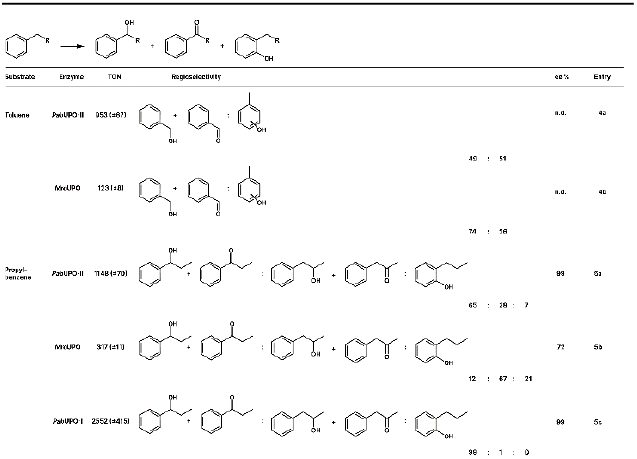
Summary of the most selective conversions from the screening of the aromatics substrate pool. Reaction conditions: Final volume 500 µl, potassium phosphate buffer (100 mM, pH = 7), 5 v/v % acetone, substrate — (4 mM), hydrogen peroxide — (1 mM), 25 °C, 400 rpm, 1 h. Concentration was determined by GC-MS. Chirality measurements were performed with a chiral column on a GC-MS.

The formation of 1-phenyl-1-propanol was detected for every UPO tested (Figure 3). *Pab*UPO-I exhibited a very high regioselectivity ratio of 99:1 towards the production of 1-phenyl-1-propanol (Entry 5c) as previously reported in the literature.^19^ The reaction proceeded with a TON of 2552 and a high enantiomeric excess of 99 %. The oxidation of non-activated positions, adjacent to the benzylic position was performed by *Pab*UPO-II, albeit the regioselective ratio of 65:28 still favors the benzylic position over all other positions (Entry 5a). Interestingly within this setup, aromatic ring oxidation at the ortho position was observed, accounting for an amount of 7 % of the total products. The oxidation of the benzylic position was performed with a TON of 1148 and an enantiomeric excess of 99 %, favoring the (R)-enantiomer. *Mro*UPO showed a switch in selectivity, as the oxidation towards 1-phenyl-2-propanol was the major product, accounting for 67 % of the total product formation with a TON of 317 (Entry 5b). 21 % of the overall products were formed by the oxidation at the aromatic moiety. A similar result was observed regarding the oxidation of toluene to benzyl alcohol and cresol products, respectively, by *Pab*UPO-II (Entry 4a). A regioselective ratio of 49:51 was observed, showing near parity between the benzylic position and the oxidation of the aromatic moiety (a mixture of all cresol products was obtained). The cresols were obtained with a TON of 953. *Mro*UPO provided the most selective conversion of toluene towards benzyl alcohol with a regioselective ratio of 74:26 and a TON value of 123 (Entry 4b). The conversion of phenylpropane to 1-phenyl-2-propanol by *Mro*UPO breaks the hegemony of UPOs oxidizing the benzylic position, opening up new synthetic routes, and broadening the synthetic usefulness of UPOs.

**Figure 3.**
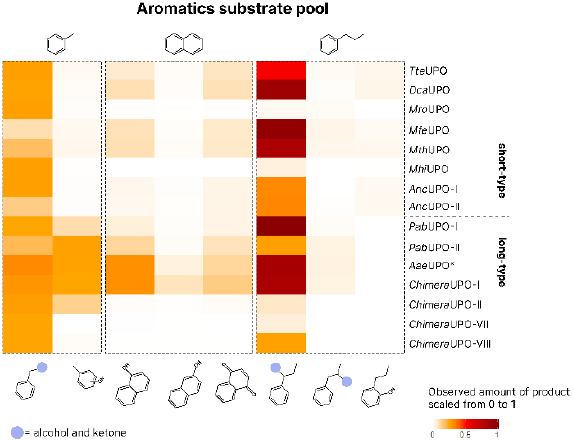
Heatmap of the aromatic substrate pool, displaying the activities of 15 different UPOs. Reaction conditions: Final volume 400 µl, potassium phosphate buffer (100 mM, pH = 7), 5 v/v % acetone, substrate — (4 mM), hydrogen peroxide — (1 mM), 25 °C, 400 rpm, 1 hour. Concentration was determined by GC-MS. Concentration was determined by GC-MS. Chirality measurements were performed with a chiral column on a GC-MS.

### Fingerprinting of 15 UPOs with a substrate pool of alkenes

The epoxidation substrate pool consists of multiple substrates possessing C=C bonds. The utilized substrates cyclohexene, hexene, styrene, and beta-methyl-styrene led to the occurrence of nine different products. The results of the screening of the epoxide substrate pool are displayed as a heatmap in Figure 4.

**Figure 4.**
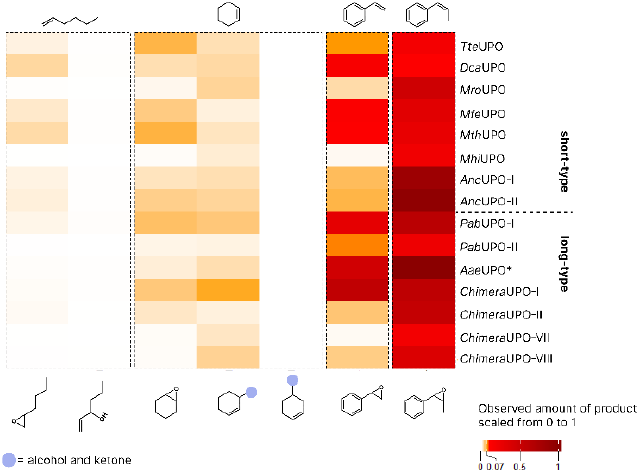
Heatmap of the epoxide substrate pool, displaying the activities of 15 different UPOs. Reaction conditions: Final volume 400 µl, potassium phosphate buffer (100 mM, pH = 7), 5 v/v % acetone, substrate — (4 mM), hydrogen peroxide — (1 mM), 25 °C, 400 rpm, 1 hour. Concentration was determined by GC-MS. Chirality measurements were performed with a chiral column on a GC-MS.

### Detailed analysis of most promising UPO/substrate pairs from the substrate pool of alkenes

Table 3 shows the results of single substrate reactions of the most promising variants from the initial fingerprinting.

**Table 3:**
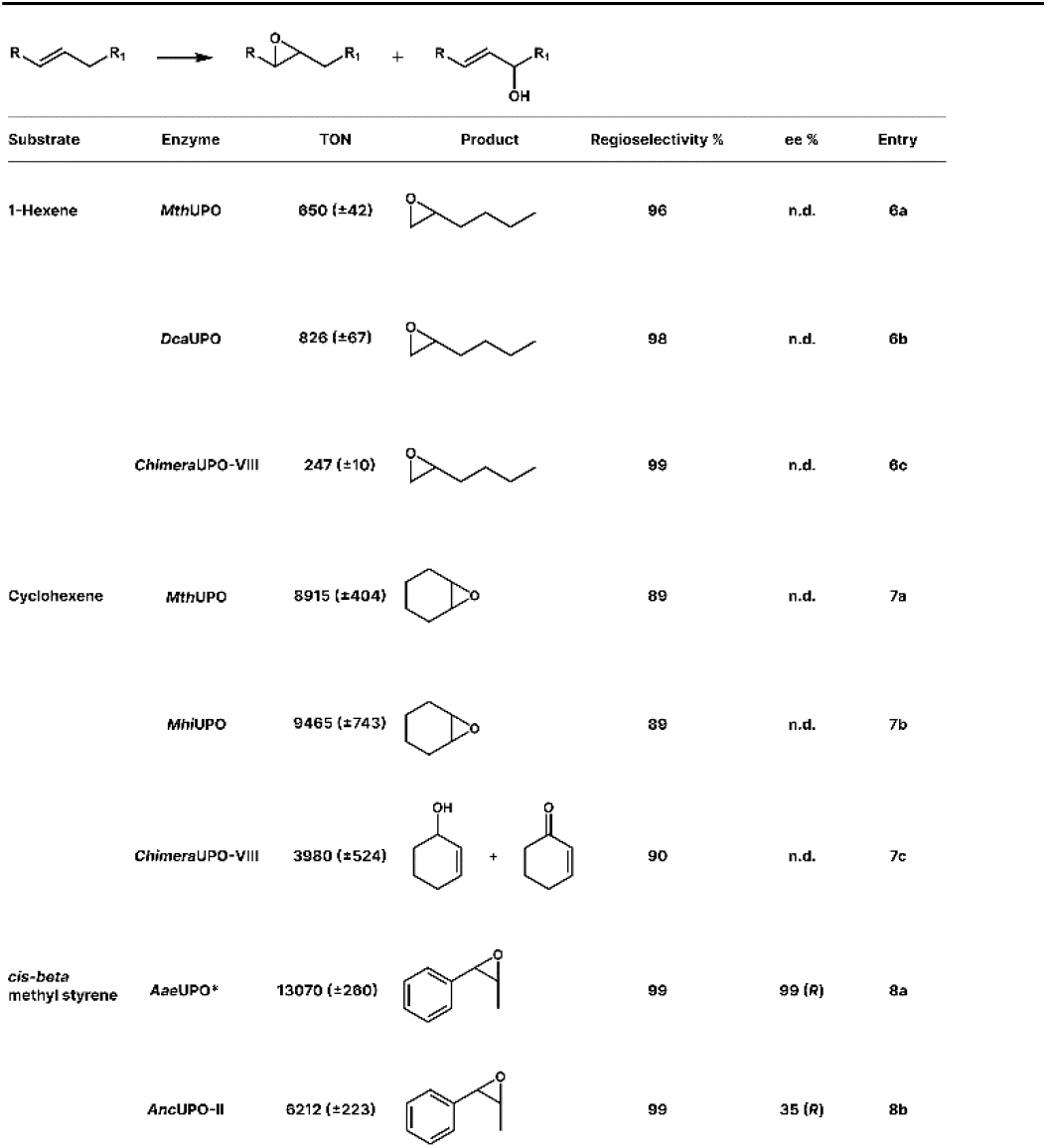
Summary of the most selective conversions from the screening of the epoxide substrate pool. Reaction conditions: Final volume 500 µl, potassium phosphate buffer (100 mM, pH = 7), 5 v/v % acetone, substrate — (4 mM), hydrogen peroxide — (1 mM), 25 °C, 400 rpm, 1 hour. Concentration was determined by GC-MS. Concentration was determined by GC-MS. Chirality measurements were performed with a chiral column on a GC-MS.

| Substrate | Enzyme | TON | Product | Regioselectivity % | ee % | Entry |
| --- | --- | --- | --- | --- | --- | --- |
| 1-Hexene | *Mth*UPO | 650 (±42) | | 96 | n.d. | 6a |
| | *Dca*UPO | 826 (±67) | | 98 | n.d. | 6b |
| | *Chimera*UPO-VIII | 247 (±10) | | 99 | n.d. | 6c |
| Cyclohexene | *Mth*UPO | 8915 (±404) | | 89 | n.d. | 7a |
| | *Mhi*UPO | 9465 (±743) | | 89 | n.d. | 7b |
| | *Chimera*UPO-VIII | 3980 (±524) | | 90 | n.d. | 7c |
| cis-beta methyl styrene | *Aae*UPO* | 13070 (±280) | | 99 | 99 (R) | 8a |
| | *Anc*UPO-II | 6212 (±223) | | 99 | 35 (R) | 8b |

The epoxidation of 1-hexene was observed for all tested UPOs. *Mth*UPO converted 1-hexene with a TON value of 650 and a regioselective ratio of 96:4 (Entry 6a). *Dca*UPO reached a higher TON value of 826 and a regioselective ratio of 98:2 (Entry 6b), thus slightly outperforming *Mth*UPO in both categories. *Chimera*UPO-VIII proved less active, as a TON value of 247 was observed, but showed near complete selective conversion, with a regioselectivity ratio of 99:1 (Entry 6c). The epoxidation of cyclohexene proved to be less challenging, as indicated by the high TONs observed. Reactions with *Mth*UPO and *Mhi*UPO afforded TONs of 8915 and 9465, respectively (Entry 7a and 7b). Both enzymes showed a regioselective ratio of 89:11. A complete switch in selectivity was observed with *Chimera*UPO-VIII (Entry 7c). The oxidation of the allylic position was achieved with a regioselective ratio of 90:10 relative to the epoxidation while providing a high TON value of 3980. This selectivity shift presents a strong increase in regioselectivity compared to the best-studied enzyme *Aae*UPO*, with a regioselective ratio towards the allylic oxidation of 66:34. Beta-methyl-styrene was readily converted by a multitude of enzymes, showing high activity as well as excellent stereoselectivities. The ancestral reconstructed enzyme *Anc*UPO-II reached a TON of 6212 with a regioselective ratio of 99:1 (Entry 8b). This excellent regioselective ratio was also observed with *Aae*UPO*, although an even higher TON of 13070 was achieved (Entry 8a). The ability of *Chimera*UPO-VIII to oxidize the allylic position of cyclohexene presents a highlight, as these oxidations are underdeveloped in enzyme catalysis, and only a few examples are known.^20-22^

### Upscaling of four reactions to a preparative scale

To further confirm the feasibility of our identification approach, four selected reactions were realized on a preparative scale. Namely, the benzylic hydroxylation of propylbenzene (*Pab*UPO-I), the hydroxylation of propylbenzene at the non-activated position (*Mro*UPO), the epoxidation of cyclohexene (*Mhi*UPO) and the allylic hydroxylation of cyclohexene (*Chimera*UPO-VIII) were chosen. A syringe pump setup was used to enable controlled hydrogen peroxide dosing and the reaction was measured over time, to avoid overoxidation of the hydroxylated products. The time progression curves are shown in Figure 5.

**Figure 5.**
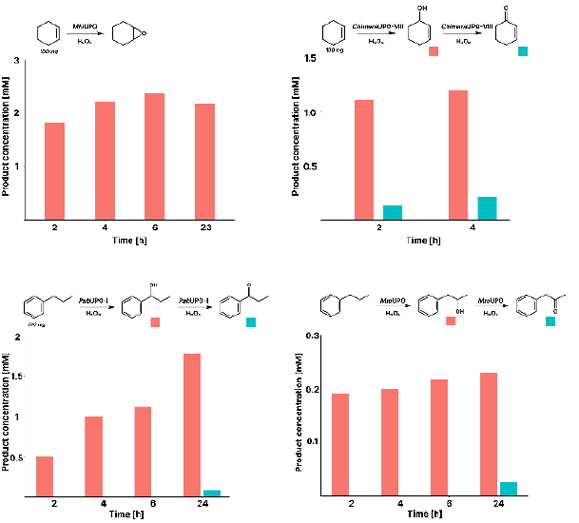
Upscaling reactions performed in a round bottom flask with a syringe pump setup to supply hydrogen peroxide. Samples were taken after the indicated time and analyzed with GC-MS. Reaction conditions: 100 nM enzyme, potassium phosphate buffer (100 mM, pH 7), 5 v/v% acetone, substrate — (4 mM), hydrogen peroxide — (1 mM). Hydrogen peroxide was added with a syringe pump at a rate of 5 mL/h. After the initial amount of 20 mL (10 mM) hydrogen peroxide was fed, the reaction was monitored with GC-MS. Additional hydrogen peroxide was then added if needed, until no further conversion was observed.

*Mhi*UPO produced epoxy cyclohexane from cyclohexene up to a concentration of 2.37 mM, which relates to a TON of 23700. The reaction was performed overnight, which seemingly led to a minor decrease in product concentration. The epoxide is seemingly not stable over longer reaction times, which was further confirmed by occurring problems in the isolation of the product (SI). The production of 2-cyclohexene-1-ol is performed by *Chimera*UPO-VIII with a TON of 11800, although minor overoxidation could already be observed after 2 hours. The alcohol product seems to be a suitable substrate for the enzyme, as the concentration of the overoxidation product (ketone) was rising linearly with the first product concentration (alcohol). *Pab*UPO-I performs the benzylic oxidation of propylbenzene with a TON of 15200. Here, longer reaction times were applied, as the overoxidation occurred only after 24 hours. The oxidation of phenylpropane towards 1-phenyl-2-propanol was performed by *Mro*UPO with low activity, as the concentration peaked at 0.23 mM, relating to a TON of 2300. This observation aligns with prior substrate screenings, in which *Mro*UPO repeatedly exhibited lower catalytic efficiencies in comparison to other recombinant UPOs.^6, 23^

### Short-vs long-type UPOs

Starting with the alkane substrate pool, no generally applicable scheme for the activity of the UPOs could be found. However, short-type UPOs proved to be superior at the oxidation of the 4-position of linear alkanes, for both octane and decane (Figure S47 and S48). While six different short-type UPOs (75 % of the respective class) enabled the oxidation of the 4-position, *Chimera*UPO-VII was found to be the only long-type UPO to catalyze this reaction. Examining the aromatic substrate pool, three enzymes (*Pab*UPO-I, *Pab*UPO-II, and *Mro*UPO) proved to be of particular interest, of which two are long-type UPOs (*Pab*UPO-I and *Pab*UPO-II) and one a short-type UPO (*Mro*UPO). Here, no type-specific pattern could be observed. The formation of cresols was found to be more pronounced for long-type UPOs (Figure 3). Otherwise, no trends could be observed. The same conclusion can be drawn from the epoxidation substrate pool, as the reaction behavior of short and long-type UPOs differed randomly.

One UPO of particular interest is *Chimera*UPO-II, as it performed the selective oxidation of linear alkanes at the 2-position. *Chimera*UPO-II was genetically constructed out of *Aae*UPO* and *Gma*UPO (Figure S2), whereby only one-fifth of the enzyme was based on *Gma*UPO. Comparing the amino acid sequences of *Aae*UPO* and *Chimera*UPO-II, both enzymes only differ at 11 positions, located in the sequence interval between positions 69 and 110. This slight change as performed before highlights enzyme chimera construction as a simple method to alter reaction specificity based on a working secretion scaffold (*Aae*UPO*).^24^

## Conclusions

We have analyzed 15 different UPOs regarding their regio and chemoselectivity towards 12 different substrates, divided into three distinct substrate pools. In this work, we describe the overall initial occurrence and testing of ancestral UPOs, which we reconstructed from the commonly used enzyme *Mth*UPO. Our approach was based on a GC-MS fingerprinting method and allowed a drastic reduction of experiments while gaining a maximal information output. Several regioselective UPOs could be identified, enabling the production of a wide range of different aliphatic and benzylic products containing a hydroxy moiety. Especially of interest were the oxidation of a sp^3^-hybridized carbon adjacent to a benzylic carbon, as well as the chemoselective oxidation of an allylic position instead of the epoxidation of the double bond. Regioselectivity ratios of up to 99:1 were obtained while already achieving non-optimized robust TON values ranging between the lower hundreds to several thousands. Additionally, highly stereoselective conversions were observed and four reactions were easily transferred to a 100 mg reaction scale. This work deepens the knowledge about the substrate profiles and selectivity of diverse UPOs and provides valuable information for future targeted applications of respective enzymes.

## Author contributions

D.H. performed all experiments. P.P. identified and designed the UPO enzymes and performed the strain construction. D.H., P.P., and M.J.W. developed and designed the experiments and wrote the manuscript.

## Acknowledgements

D.H. thanks the Friedrich-Ebert-Stiftung for a PhD scholarship.

M.J.W. thanks the Bundesministerium für Bildung und Forschung (Maßgeschneiderte Inhaltsstoffe 2, 031B0834A) and M.J.W. and D.H. thank the German Research Foundation (DFG, project ID 43649874, TP A05, RTG 2670).

