## Supplemental Information for "Mapping Selective Oxidations of Unspecific Peroxygenases"

Pascal Püllmann – Molecular Design and Engineering, Bayer AG, 42133 Wuppertal, Germany

Martin J. Weissenborn – Department of Chemistry, Martin Luther-University Halle-Wittenberg, 06120 Halle(Saale), Germany

RESTRICTED

### I. General Information

**Chemicals and Materials.** All chemicals were purchased and utilized in the highest available purity and purchased either from abcr (Karlsruhe, DE), Carl Roth (Karlsruhe, DE), TCI Chemicals (Tokyo, JP), Carbolution (Sankt Ingbert, DE) or Sigma-Aldrich (St. Louis, USA). All solvents were used in GC-Grade purity and purchased from Carl Roth (Karlsruhe, DE).

**Shake flask cultivation *P. pastoris*.** Performed as described before.[1-2] After cultivation, the cells were separated from the enzyme containing supernatant by centrifugation (5300 rpm; 35 min; 4 °C).

**Supernatant ultrafiltration and preparation.** The previously prepared *P. pastoris* supernatant was concentrated approx. 20-fold by means of ultrafiltration as previously described.[2]

**Heme CO complex measurements.** For all measurements, an acryl cuvette with a path length of 10 mm was used. Spectra were recorded on a Biospectrometer Basic device (Eppendorf, Hamburg, DE) at 444 nm (absorption) and 490 nm (reference wavelength) subtracting the utilized storage buffer (100 mM potassium phosphate pH 7.0). Heme carbon dioxide spectra (CO assay) were recorded to determine enzyme concentration in the concentrated supernatant. First, the heme iron was reduced to its ferrous form ($Fe^{2+}$). For this, a spatula tip of sodium dithionite as the reducing agent was added to 2 ml of the concentration enzyme supernatant sample and mixed thoroughly until complete dissolution. The sample was divided into three parts. Following, fresh saturated carbon monoxide buffer was prepared by flushing the utilized buffer with a constant carbon monoxide flow for at least 5 minutes. 0.25 ml of fresh CO-buffer was added to two 0.75 ml reduced supernatant samples (duplicates) in cuvettes to obtain the heme-CO complex. The absorption was immediately measured as described above. The third sample was treated with non-gassed 0.25 ml buffer and used as supernatant blank reference. The extinction coefficient was determined with a calibration curve of *Mth*UPO wildtype in storage buffer (10 nM – 1000 nM) leading to ε (444 nM – 490 nM) = 70,700 $M^{-1}$ $cm^{-1}$. The enzyme concentration in the supernatant was then calculated using the formula:

$$c[\mu M] = df x \frac{4}{3} x \frac{Sample(A_{444nm} - A_{490\,nm}) - SB(A_{444\,nm} - A_{490\,nm})}{0{,}707\ \mu M^{-1} cm^{-1}}$$

With df = dilution factor and SB = supernatant blank reference.

**Multiple substrate reaction setup for initial screening.** Reactions were performed in 2 ml reaction vials from Carl Roth (Karlsruhe, DE). A stock solution containing all substrates (e.g. hexane, octane, decane, and cyclohexane) in acetone and a stock solution containing hydrogen peroxide in potassium

RESTRICTED

phosphate buffer (100 mM, pH = 7) were prepared freshly before the experiment. The reaction volume was set to 400 µl with the following conditions: 300 nM enzyme, 5 v/v% acetone, 4 mM substrate, 1 mM $H_2O_2$, potassium phosphate buffer (100 mM, pH = 7). The reaction was performed in a shaker at 400 rpm and 25 °C for 1 hour. The reaction was quenched by addition of 400 µl ethyl acetate containing an internal standard (250 µM) and the extraction was performed at 800 rpm for 30 minutes. 150 µl of the organic layer was transferred for GC-MS analysis.

**Single substrate reaction setup.**

Reactions were performed in 2 ml reaction vials from Carl Roth (Karlsruhe, DE). A stock solution containing the substrate in acetone and a stock solution containing hydrogen peroxide in potassium phosphate buffer (100 mM, pH = 7) were prepared freshly before the experiment. The reaction volume was set to 500 µl with the following conditions: 100 nM enzyme, 5 v/v% acetone, 4 mM substrate, 1 mM $H_2O_2$, potassium phosphate buffer (100 mM, pH = 7). The reaction was performed in a shaker at 400 rpm and 25 °C for 1 hour. The reaction was quenched by addition of 400 µl ethyl acetate containing an internal standard (250 µM) and the extraction was performed at 800 rpm for 30 minutes. 150 µl of the organic layer was transferred for GC-MS analysis.

**GC-MS measurements.** Measurements were performed on a Shimadzu GCMS-QP2010 Ultra instrument (Shimadzu, Kyoto, JP) with an SH-Rxi-5Sil MS column (30 m x 0.25 mm, Shimadzu, Kyoto, JP) for achiral measurements, utilizing helium as carrier gas. The samples were injected splitless or with split (1 µl) with a liner temperature of 280 °C. The interface temperature was set to 290 °C. Ionisation was obtained by electron impact with a voltage of 70 V, and the temperature of the ion source was 250 °C. The oven temperature profile for each compound is shown in Table 2. The detector voltage of the secondary electron multiplier was adjusted to the tuning results with perfluorotributylamine. The GC-MS parameters were controlled with GCMS Real Time Analysis, and for data evaluation, GCMS Postrun Analysis (GCMSsolution Version 4.45, Shimadzu, Kyoto, JP) was used.

Chrial measurements were performed with a Lipodex E column (30 m x 0.25 mm, Macherey Nagel,Düren,DE) utilizing helium as carrier gas.

**Derivatization for chiral measurements.** Reactions were performed as described above. The reaction was quenched by addition of 400 µl DCM and the extraction was performed at 800 rpm for 30 minutes. 150 µl of the organic layer were transferred and 2 v/v % trifluoro acetic anhydride were added. The mixture was vortexed for 1 minute at room temperature and analysed with GC-MS.

RESTRICTED

**Data processing after screening.** Each product was assigned with a specific combination of two *m/z* values, which were used for identifying the product (Table 1). All products were identified by three independent samples. Following the identification, the measured peak area of the product was divided by the area of the internal standard. The obtained values were used to calculate the mean value, which was subtracted with background reactions. Following this, the values were divided by the intensity factor of the products. This factor was determined by measuring all required products with the same concentration at the GC-MS and comparing the peak area of the different *m/z* used. By this, the differences in ionization could be neglected and made all products for all enzymes comparable. Depending on the products, ketones and alcohols were added up, to represent the oxidation at a certain position (e.g. 2-octanol and 2-octanone for the oxyfunctionalization at the 2-position of octane).

**Preparative Scale conversion – General remarks**

Preparative Scale conversions proved to be challenging, as the extraction of the products was either bad or insufficient. Only 1-phenyl-1-propanol and 2-cyclohexene-1-ol were extracted sufficiently and therefore only from these two NMR-spectra were recorded. Nonetheless, the other products were identified in the reaction mixture by GC-MS and the amount was determined by calibration curves.

**Preparative Scale conversion of propylbenzene towards 1-phenyl-1-propanol.**

Propylbenzene (103 µl, 0.74 mmol) was dissolved in acetone (13 ml) and poured into a solution of potassium phosphate buffer (200 ml, 100 mM, pH 7.0) and 100 nM *Pab*UPO-I (22 ml, stock solution 1160 nM). 1 mM hydrogen peroxide (20 ml, stock solution 10 mM) was added via a syringe pump with a rate of 5 ml/h. The reaction was monitored over time via GC-MS and additional hydrogen peroxide was added if needed. The reaction was stirred over night at 30 °C. The mixture was extracted with ethyl acetate (3x 100 ml), the organic phase was washed with brine and dried with sodium sulfate. The solvent was removed *in vacuo* and the residue was purified by column chromatography on silica gel using n-hexane/ethyl acetate with a gradient (5:5 → 9:1).

**Preparative Scale conversion of propylbenzene towards 1-phenyl-2-propanol.**

Propylbenzene (103 µl, 0.74 mmol) was dissolved in acetone (3.5 ml) and poured into a solution of potassium phosphate buffer (22.5 ml, 100 mM, pH 7.0) and 100 nM *Mro*UPO (24.2 ml, stock solution 340 nM). 1 mM hydrogen peroxide (20 ml, stock solution 10 mM) was added via a syringe pump with a rate of 5 ml/h. The reaction was monitored over time via GC-MS and additional hydrogen peroxide was added if needed. The reaction was stirred over night at 30 °C. The mixture was extracted with ethyl

RESTRICTED

acetate (3x 100 ml), the organic phase was washed with brine and dried with sodium sulfate. The solvent was removed *in vacuo*. Extraction was not sufficient, as the product could not be isolated.

**Preparative Scale conversion of cyclohexene towards 2-cyclohexen-1-ol.**

Cyclohexene (145 µl, 0.74 mmol) was dissolved in acetone (5 ml) and poured into a solution of potassium phosphate buffer (11 ml, 100 mM, pH 7.0) and 100 nM *Mhi*UPO (27.2 ml, stock solution 184 nM). 1 mM hydrogen peroxide (5 ml, stock solution 10 mM) was added via a syringe pump with a rate of 5 ml/h. The reaction was monitored over time via GC-MS and additional hydrogen peroxide was added if needed. The reaction was stirred for 4 hours at 30 °C. The mixture was extracted with ethyl acetate (3x 100 ml), the organic phase was washed with brine and dried with sodium sulfate. The solvent was removed *in vacuo* and the residue was purified by column chromatography on silica gel using n-hexane/ethyl acetate (3:7).

**Preparative Scale conversion of cyclohexene towards epoxycyclohexane.**

Cyclohexene (104 µl, 0.74 mmol) was dissolved in acetone (13 ml) and poured into a solution of potassium phosphate buffer (227 ml, 100 mM, pH 7.0) and 100 nM *Mhi*UPO (15.5 ml, stock solution 1655 nM). 1 mM hydrogen peroxide (20 ml, stock solution 10 mM) was added via a syringe pump with a rate of 5 ml/h. The reaction was monitored over time via GC-MS and additional hydrogen peroxide was added if needed. The reaction was stirred over night at 30 °C. The mixture was extracted with ethyl acetate (3x 100 ml), the organic phase was washed with brine and dried with sodium sulfate. The solvent was removed *in vacuo*. The product was not isolated, as purification by column chromatography led to the diminishing of the epoxide.

**UPO sequences**

Signal peptide

Mature protein

C-terminal TwinStrep-GFP11 tag

*Tte*UPO

MRFPSIFTAVLFAASSALAAPVNTTTEDETAQIPAEAVIGYLDLEGDFDVAVLPFSNSTNNGLLFINTTIA
SIAAKEEGVSLDKREAGFDSWHPPAPGDRRGPCPMLNTLANHGFLPHNGRNITKEITVNALNSALNVNK
TLGELLFNFAVTTNPQPNATFFDLDHLSRHNILEHDASLSRADYYFGHDDHTFNQTVFDQTKSYWKTPII
DVQQAANARLARVLTSNATNPTFVLSQIGEAFSFGETAAYILALGDRVSGTVPRQWVEYLFENERLPLE
LGWRRAKEVISNSDLDQLTNRVINATGALANITRKIKVRDFHAGRFPGEGGGSAWSHPQFEKGGGSGG
GSGGSAWSHPQFEKDGGSGGGSTSRDHMVLHEYVNAAGIT

*Dca*UPO

RESTRICTED

MFAFYFLTACISLKGVFGAGFAPWKAPGPDDVRGPCPMLNTLANHGFLPHDGKNIDVNTTVNALSSAL
NLDDELSRDLHTFAVTTNPQPNATWFSLNHLSRHNVLEHDASLSRQDAYFGPPDVFNAAVFNETKAY
WTGDIINFQMAANALTARLMTSNLTNPEFSMSQLGRGFGLGETVAYVTILGSKETRTVPKAFVEYLFEN
ERLPYELGFKKMKSALTEDELTTMMGEIYSLQHLPESFTKPFAKRSEAPFEKRAEKRCPFHSGGSAWSH
PQFEKGGGSGGGSGGSAWSHPQFEKDGGSGGGSTSRDHMVLHEYVNAAGIT

*Mro*UPO

MKLAISSSLIALVSVTTALANSQDVVDFGASAHPWKAPGPNDSRGPCPGLNTLANHGFLPRNGRNISVP
MIVKAGFEGYNVQSDILILAGKIGMLTSREADTISLEDLKLHGTIEHDASLSREDVAIGDNLHFNEAIFTT
LANSNPGADVYNISSAAQVQHDRLADSLARNPNVTNTDLTATIRSSESAFFLTVMSAGDPLRGEAPKKF
VNVFFREERMPIKEGWKRSTTPITIPLLGPIIERITELSDWKPTGDNCGAIVLSPELGGSAWSHPQFEKGG
GSGGGSGGSAWSHPQFEKDGGSGGGSTSRDHMVLHEYVNAAGIT

*Mfe*UPO

MLGKNDPMCLVLVLLGLTALLGICQGAGFDTWSPPGPYDVRAPCPMLNTLANHGFLPHDGKDITRENT
ENALFEALHINKTLGSFLFDFALTTNPRNTSTFSLNDLGNHNILEHDASLSRADAYFGNVLQFNQTVFDE
TKTYWDGDVIDLRMAARARLGRIKTSQATNPTYSMSELGDAFTYGESAAYVVVLGDKESRTAKRSWV
EWFFEHEQLPQHLGWKRPASSLEEEDLFTIMDEIRQYTSELEGSTSSSDAQTSRRQLPRRRTHFGFGGSA
WSHPQFEKGGGSGGGSGGSAWSHPQFEKDGGSGGGSTSRDHMVLHEYVNAAGIT

*Mth*UPO

MLGKNDPMCLVLVLLGLTALLGICQGAGFDTWSPPGPYDVRAPCPMLNTLANHGFLPHDGKDITREQT
ENALFEALHINKTLASFLFDFALTTNPKNTSTFSLNDLGNHNILEHDASLSRADAYFGNVLQFNQTVFDE
TKTYWEGDTIDLRMAAKARLGRIKTSQATNPTYSMSELGDAFTYGESAAYVVVLGDKESRTVKRSWV
EWFFEHEQLPQHLGWKRPAASFEEEDLNSSMEEIEKYTKELEGSNSTSGSQKHRRRLPRRRAHFGFSGG
SAWSHPQFEKGGGSGGGSGGSAWSHPQFEKDGGSGGGSTSRDHMVLHEYVNAAGIT

*Mhi*UPO

MFAFYFLTACISLKGVFGAGFDTWSPPGPTDVRAPCPMLNTLANHGFLPHDGKDITREQTENALFDALN
INKTLASFLFDFALTTNPKNTSTFSLNDLGNHNILEHDASLSRADAYFGNVLQFNQTVFDETKTYWEGD
TIDLRMAAKARLGRIKTSQATNPTYSMSELGDAFTYGESAAYVVVLGDLESRTVNRSWVEWFFEHEQL
PQHLGWKRPAVSFEEEDLNRFMEEIEKYTKGLEGSNSTSGSQKHRRRLPRRRTHFGFGGSAWSHPQFEK
GGGSGGGSGGSAWSHPQFEKDGGSGGGSTSRDHMVLHEYVNAAGIT

*Anc*UPO-I

MRFPSIFTAVLFAASSALAAPVNTTTEDETAQIPAEAVIGYLDLEGDFDVAVLPFSNSTNNGLLFINTTIA
SIAAKEEGVSLDKREAGFDPWQPPGPGDVRSPCPALNTLANHGFLPHNGKNITMDVLIKAFKEGLNIGP
DVATVVGGAALTTNPTAPNATTFDLDDLNKHNVIEHDGSLSRQDFYFGDNHSFNQEIWDQTLSYFTGN
ETISIETAAKARAARVATAKATNPEFNLTASQQAASLGETALYLSVFGDPVTGNAPKDWVKVFFENERL
PYELGWTPPETEITLTTLGAMVGKIAAASGGSAWSHPQFEKGGGSGGGSGGSAWSHPQFEKDGGSGGG
STSRDHMVLHEYVNAAGIT

*Anc*UPO-II

MFSKVLPFVGAVAALPHSVRAGFSEWQPPGPGDVRGPCPMLNTLANHGFLPHDGKNITQEQTINALGE
ALNIDKELAQFLFFDFAITTNPEPNATTFSLDDLSRHNILEHDASLSRADAYFGNAHTFNQTVFDETRSY
WTGPIIDVQMAANARLARLQTSNATNPTYTLSELGEAFSFGETAAYIIVLGDKTGTVPRSWVEYFFENE
RLPTELGWKRPEEAFTQDDLNDMMERIINATGATQGSAKTKRRRIHRKRRGFSGGSAWSHPQFEKGGG
SGGGSGGSAWSHPQFEKDGGSGGGSTSRDHMVLHEYVNAAGIT

*Pab*UPO-I

MRGTPIFASLIALFAHAAIAFPAYGSLAGLTREQLDEILPTLEIRADPPTPPGPLAYNGTKLVHDDAHPFK
APEQGDIRGPCPGLNTLANHGYLPHNGVATPSQIIEAVQEGFNMEHATALFVTYAAHLVDGNLVTDLLS
IGEKTGLTGLDPPAPAIVGGLNTHAVFEGDASMTRADFFFGDNHNFNQTLFDQFVDFSNRFGGGFYNYT
VAAELRFQRIQESIATNPQFSFISPRFFTAYAESTFPVNFFVDGRSTEKKLDMEAATSFIRDGKYPQDFHR

RESTRICTED

AAQPSSTEGIDIVASAHPVAPGENRDGKINNYVPDPTSADFSTFCLLYTNFVNQTIGGLYPNPTGVLRRN
LIKNLRFFYSGIADAGCEELFPYGQLGSGGSAWSHPQFEKGGGSGGGSGGSAWSHPQFEKDGGSGGGS
TSRDHMVLHEYVNAAGIT

*Pab*UPO-II

MRGTPIFASLIALFAHAAIAFPAYGSLAGLTREQLDEILPTLEIRADPPPPPGPPSFTGTKLVNDADHPWQP
LREGDIRGPCPGLNTLASHGYLPRDGVATPAQIITATQEGFNFENNAAIVATYLGHLLNGNLVTDLLSIG
GATPKTGPPPPPPAHAGGLNVHGTFEGDAGMTRADEFFGDNHSFNQTLFDKFVDFSNRYGGGFYNLTV
AGELRYSRIQDSIATNPEFQFKNVRFITAYGETVFPINLFVDGRVTTDRKLSMEDAASIFRDMRFPDDFH
RSAVPASNEGADQVLAAHPWVPGGNADNQVNNYVEDPDSADFTHLCRLYEFVVGSVQELYPNPTGIL
RRNLIKNLHYWWTGVNVAFGGCDELFPYGQLGSGGSAWSHPQFEKGGGSGGGSGGSAWSHPQFEKD
GGSGGGSTSRDHMVLHEYVNAAGIT

*Aae*UPO*

MRGTPIFASLIALFAHAAIAFPAYGSLAGLTREQLDEILPTLEIRAEPGLPPGPLENSSAKLVNDEAHPWK
PLRPGDIRGPCPGLNTLASHGYLPRNGVATPAQIINAVQEGFNFDNQAAIFATYAAHLVDGNLITDLLSI
GRKTRLTGPDPPPPASVGGLNEHGTFEGDASMTRGDAFFGNNHDFNETLFEQLVDYSNRFGGGKYNLT
VAGELRFKRIQDSIATNPNFSFVDFRFFTAYGETTFPANLFVDGRRDDGQLDMDAARSFFQFSRMPDDF
FRAPSPRSGTGVEVVVQAHPMQPGRNVGKINSYTVDPTSSDFSTPCLMYEKFVNITVKSLYPNPTVQLR
KALNTNLDFLFQGVAAGCTQVFPYGRDSGGSAWSHPQFEKGGGSGGGSGGSAWSHPQFEKDGGSGGG
STSRDHMVLHEYVNAAGIT

*Chimera*UPO-I

MRGTPIFASLIALFAHAAIAFPAYGSLAGLTREQLDEILPTLEIRAEPGLPPGPLENSSAKLVNDEAHPWK
PLRPGDIRGPCPGLNTLASHGYLPRNGVATPAQIINAVQEGFNFDNQAAIFATYAAHLVDGNLITDLLSI
GRKTRLTGPDPPPPASVGGLNEHGTFEGDASMTRGDAFFGNNHDFNETLFEQLVDYSNRFGGGKYNLT
VAGELRFKRIQDSIATNPNFSFVDFRFFTAYGETTFPANLFVDGRRDDGQLDMDAARSFFQFSRMPDDF
FRAPSPRSGTGVEVVVQAHPMQPGRNVGKINSYTVDPTSADFNTFCLLYENFVNKTIKGLYPNPTGALR
KALNTNLGFFFSGISDTGCTQVFPYGKGSGGSAWSHPQFEKGGGSGGGSGGSAWSHPQFEKDGGSGGG
STSRDHMVLHEYVNAAGIT

*Chimera*UPO-II

MRGTPIFASLIALFAHAAIAFPAYGSLAGLTREQLDEILPTLEIRAEPGLPPGPLENSSAKLVNDEAHPWK
PLRPGDIRGPCPGLNTLASHGYLPRNGVATPAQIINAVQEGFNMDNSLAIFVTYAAHLVDGNILTDKLSI
GGKTALTGPNPPAPASVGGLNEHGTFEGDASMTRGDAFFGNNHDFNETLFEQLVDYSNRFGGGKYNLT
VAGELRFKRIQDSIATNPNFSFVDFRFFTAYGETTFPANLFVDGRRDDGQLDMDAARSFFQFSRMPDDF
FRAPSPRSGTGVEVVVQAHPMQPGRNVGKINSYTVDPTSSDFSTPCLMYEKFVNITVKSLYPNPTVQLR
KALNTNLDFLFQGVAAGCTQVFPYGRDSGGSAWSHPQFEKGGGSGGGSGGSAWSHPQFEKDGGSGGG
STSRDHMVLHEYVNAAGIT

*Chimera*UPO-VII

MRGTPIFASLIALFAHAAIAFPAYGSLAGLTREQLDEILPTLEIRAEPGLPPGPLENSSAKLVNDEAHP
WKPLRPGDIRGPCPGLNTLASHGYLPRNGVATPAQIINAVQEGFNMDNSLAIFVTYAAHLVD
GNILTDKLSIGGKTALTGPNPPAPAIVGGLNTHAVFEGDTSMTRGDFFFGNNHDFNETLFDEF
VDFSNRFGGGKYNLTVAGEFRWQRIQDSIATNGQFDFTSPRYFTAYAESVFPINFFTDGRLFTS
NTTAPGPDMDSALSFFRDHRYPKDFHRAPVPSGARGLDVVAAAYPIQPGYNADGKVNNYVL
DPTSSDFSTPCLMYEKFVNITVKSLYPNPTVQLRKALNTNLDFLFQGVAAGCTQVFPYGRDGG
SAWSHPQFEKGGGSGGGSGGSAWSHPQFEKDGGSGGGSTSRDHMVLHEYVNAAGIT

*Chimera*UPO-VIII

MRGTPIFASLIALFAHAAIAFPAYGSLAGLTREQLDEILPTLEIRAEPGLPPGPLENSSAKLVNDEAHPWK
PLRPGDIRGPCPGLNTLASHGYLPRNGVATPAQIINAVQEGFNMDNSLAIFVTYAAHLVDGNILTDKLSI

RESTRICTED

GGKTALTGPNPPAPAIVGGLNTHAVFEGDASMTRGDFHLGDNFNFNQTLWEQFKDYSNRYGGGRYNL
TAAAELRWARIQQSMATNPNFSFVSPRYFTAYAESTFPINFFIDGRQNDGQLNLTVARGFFQNSRMPDG
FHRANGTRGTEGIDVIAEAHPIEPGSNVGGVNNYVVDPTSSDFSTPCLMYEKFVNITVKSLYPNPTVQLR
KALNTNLDFLFQGVAAGCTQVFPYGRDGGSAWSHPQFEKGGGSGGGSGGSAWSHPQFEKDGGSGGGS
TSRDHMVLHEYVNAAGIT

### II. Supplementary Figures

#### S1. Initial testing of screening and single substrate reaction

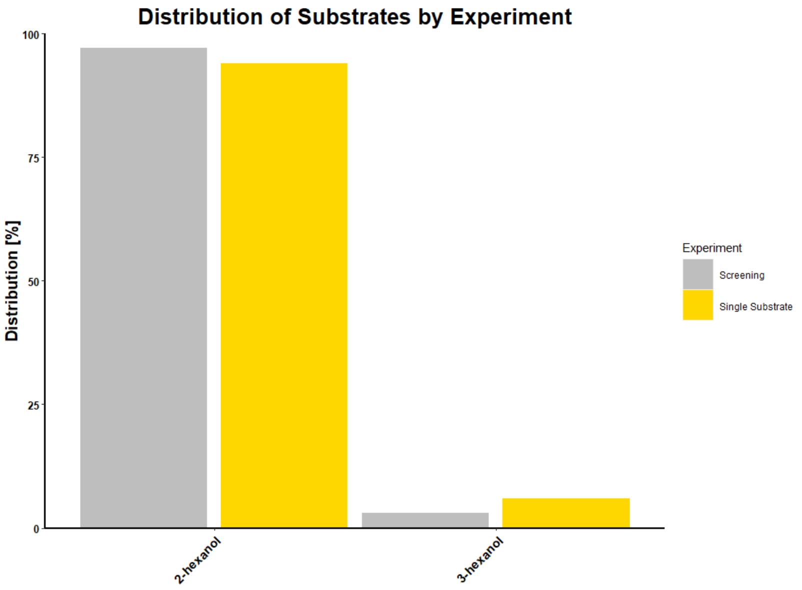


*Figure 1: Initial screening results compared to single substrate reaction.*

RESTRICTED

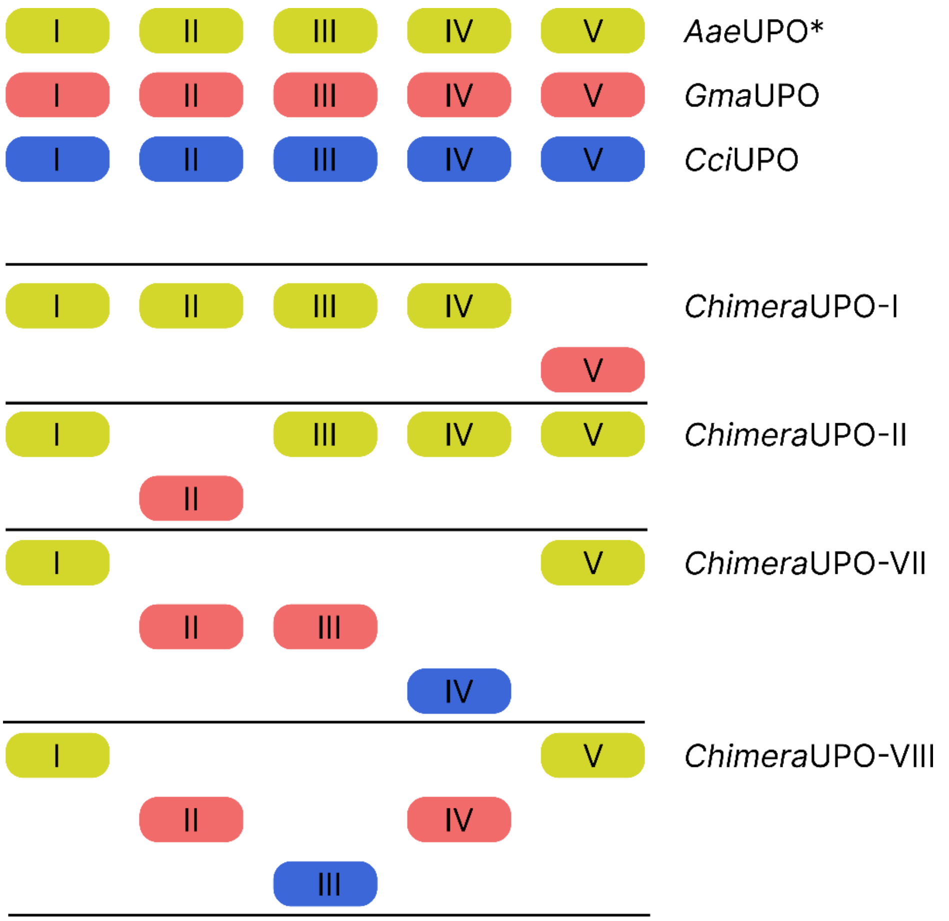


*Figure 2: Subunits of the ChimeraUPOs. The ChimeraUPOs were constructed from AaeUPO*, GmaUPO and CciUPO.*

**Calibration curves of products**

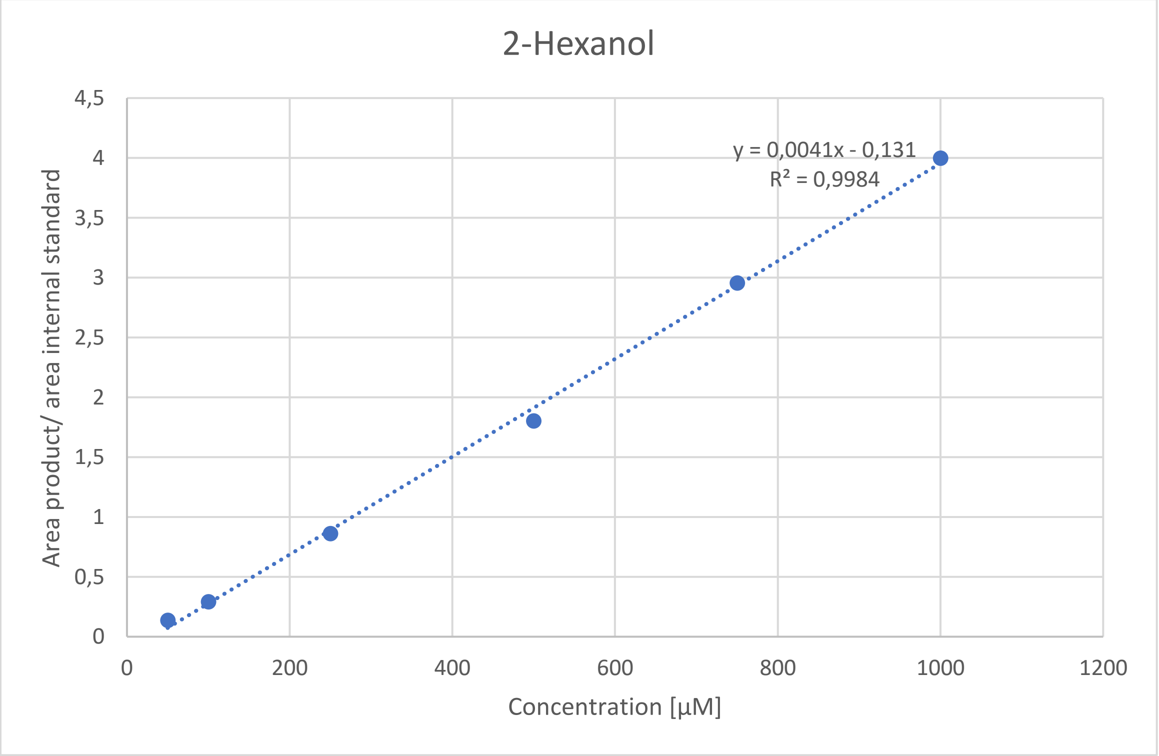


*Figure 3: Calibration curve of 2-Hexanol.*

RESTRICTED

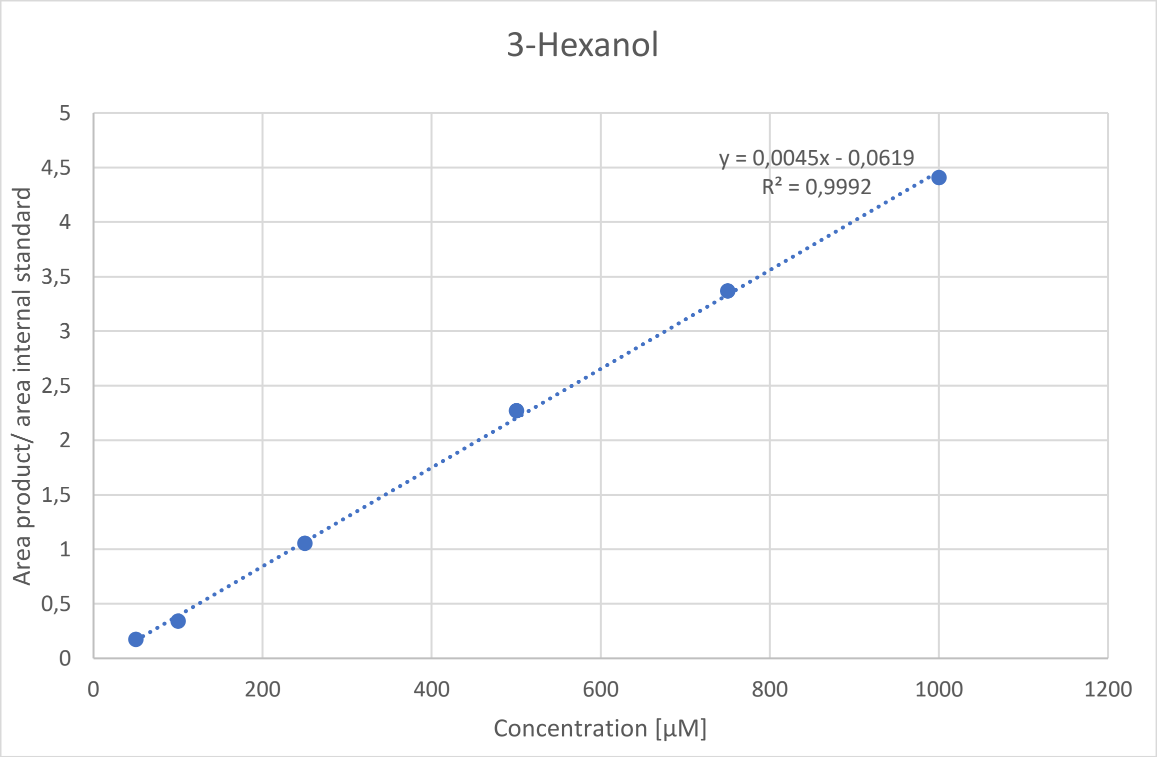


*Figure 4: Calibration curve of 3-Hexanol.*

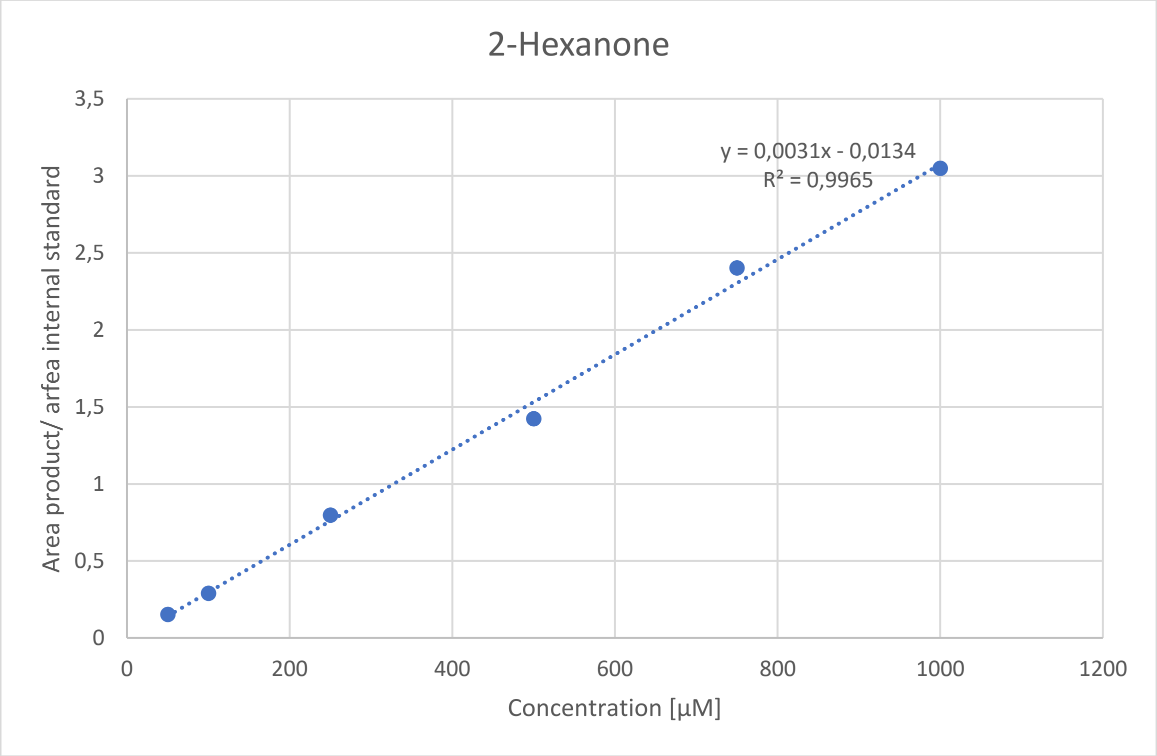


*Figure 5: Calibration curve of 2-Hexanone.*

RESTRICTED

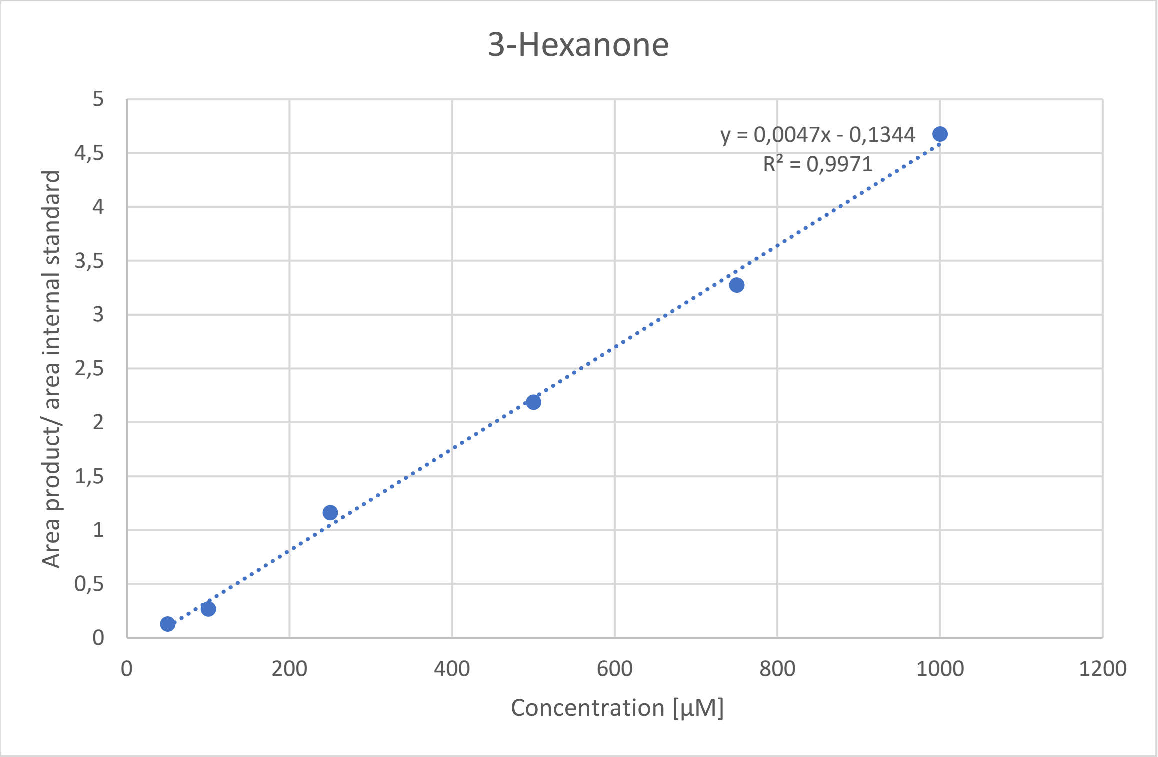


*Figure 6: Calibration curve of 3-Hexanone.*

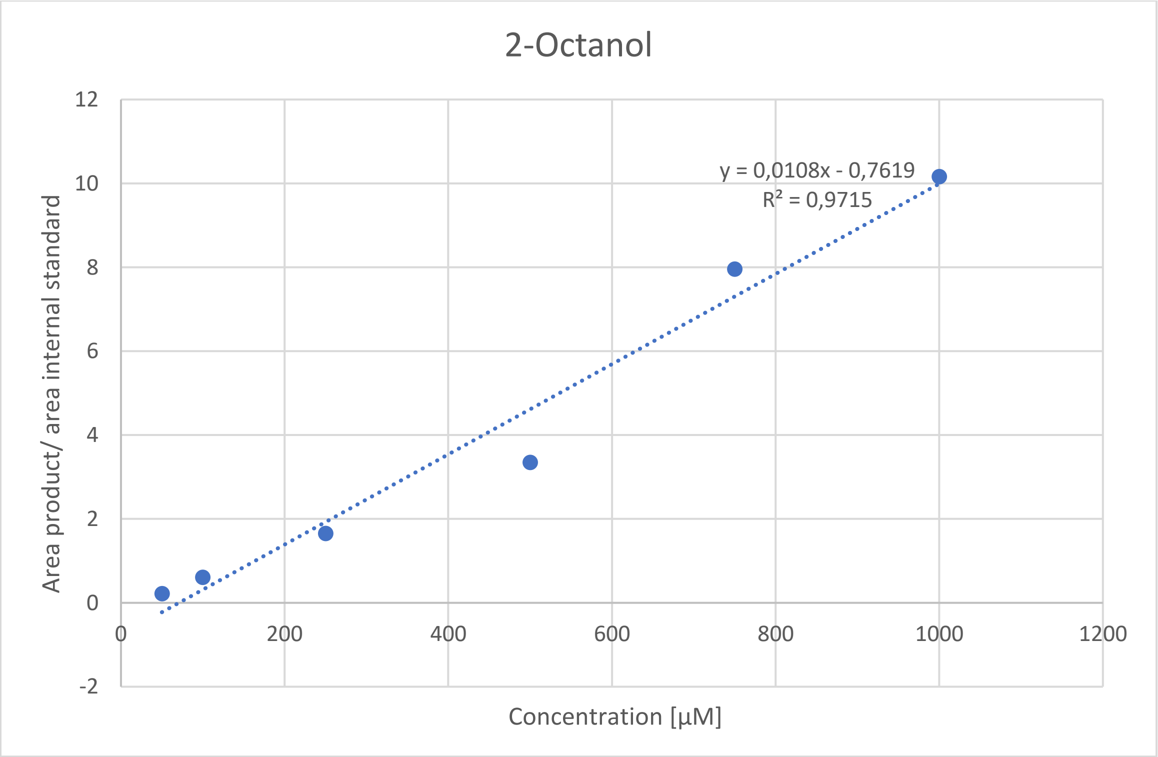


*Figure 7: Calibration curve of 2-Octanol.*

RESTRICTED

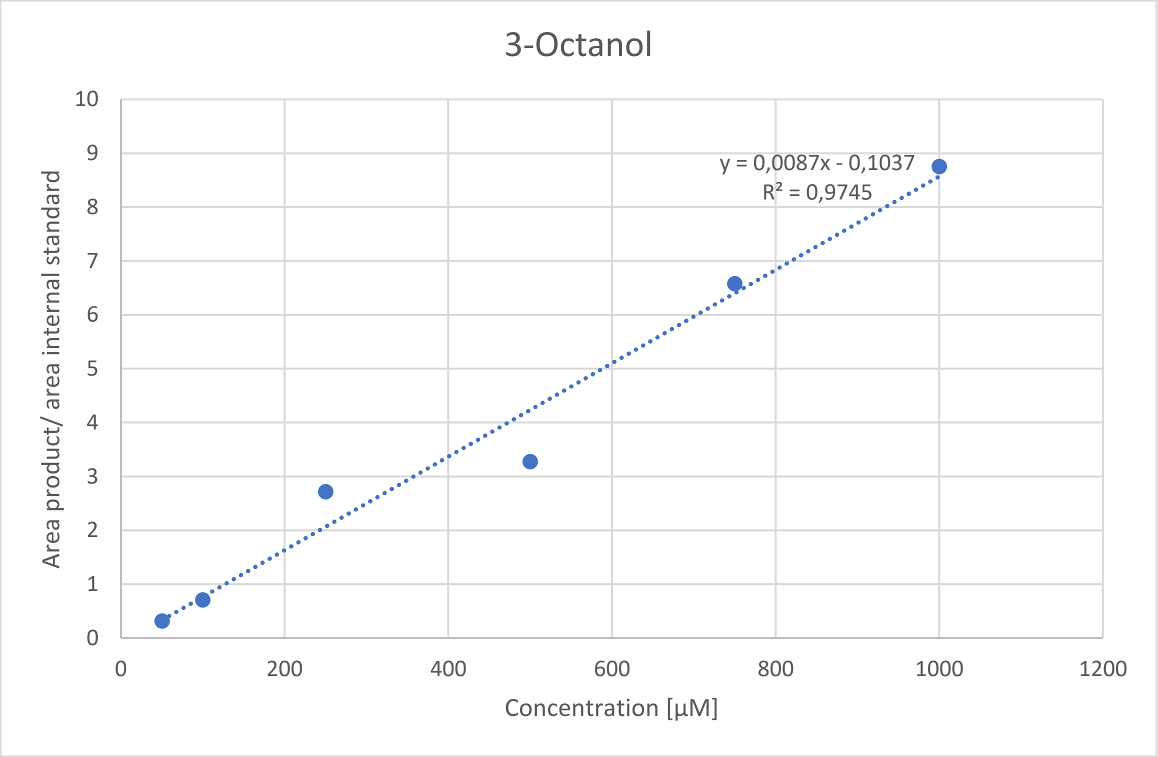


*Figure 8: Calibration curve of 3-Octanol.*

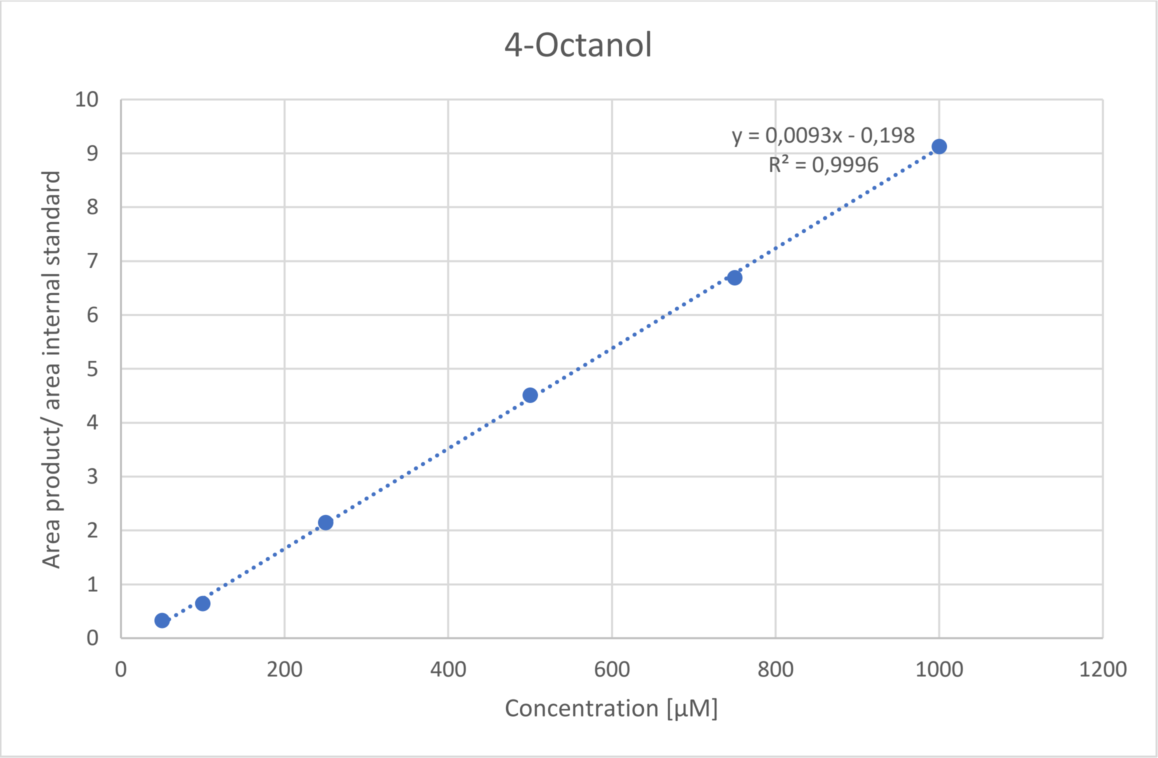


*Figure 9: Calibration curve of 4-Octanol.*

RESTRICTED

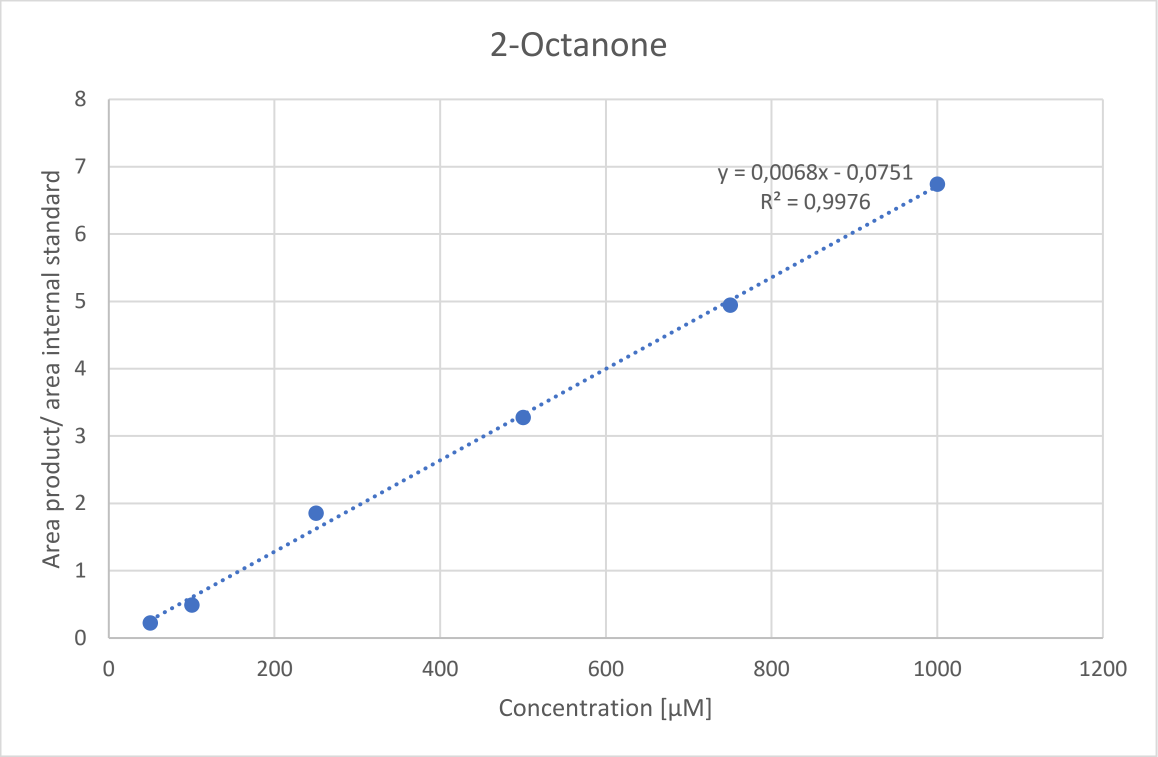


*Figure 10: Calibration curve of 2-Octanone.*

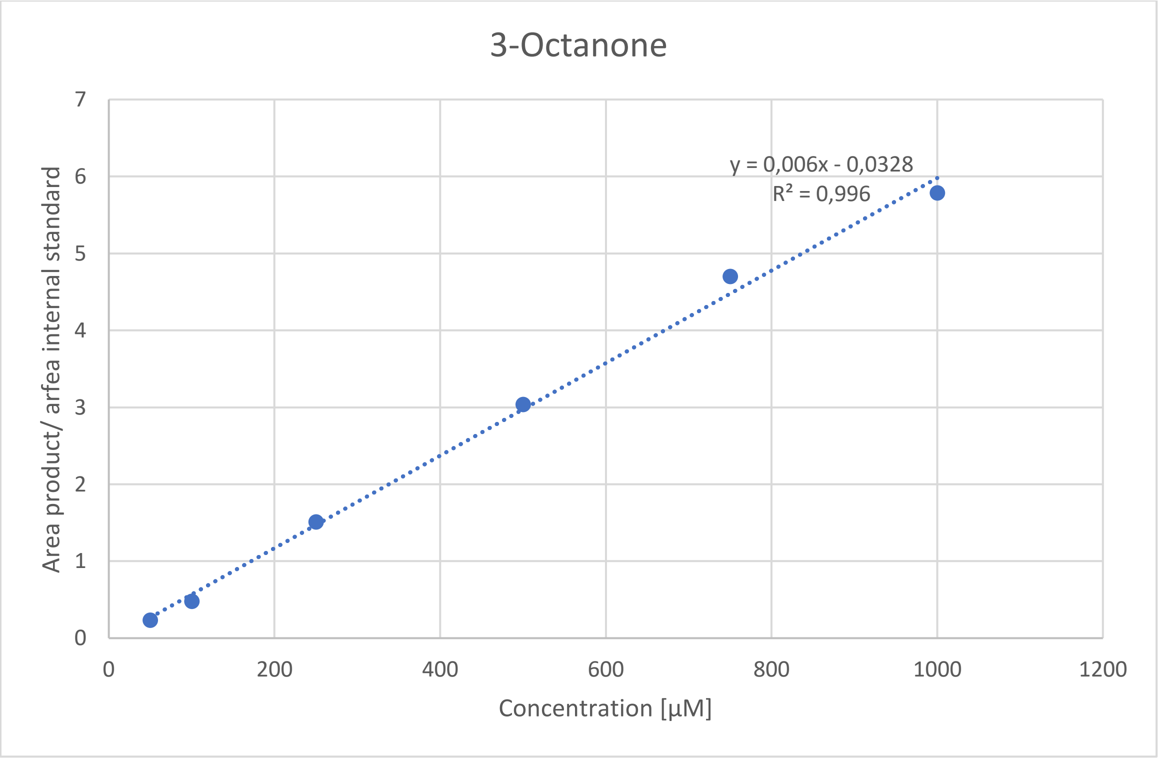


*Figure 11: Calibration of 3-Octanone.*

RESTRICTED

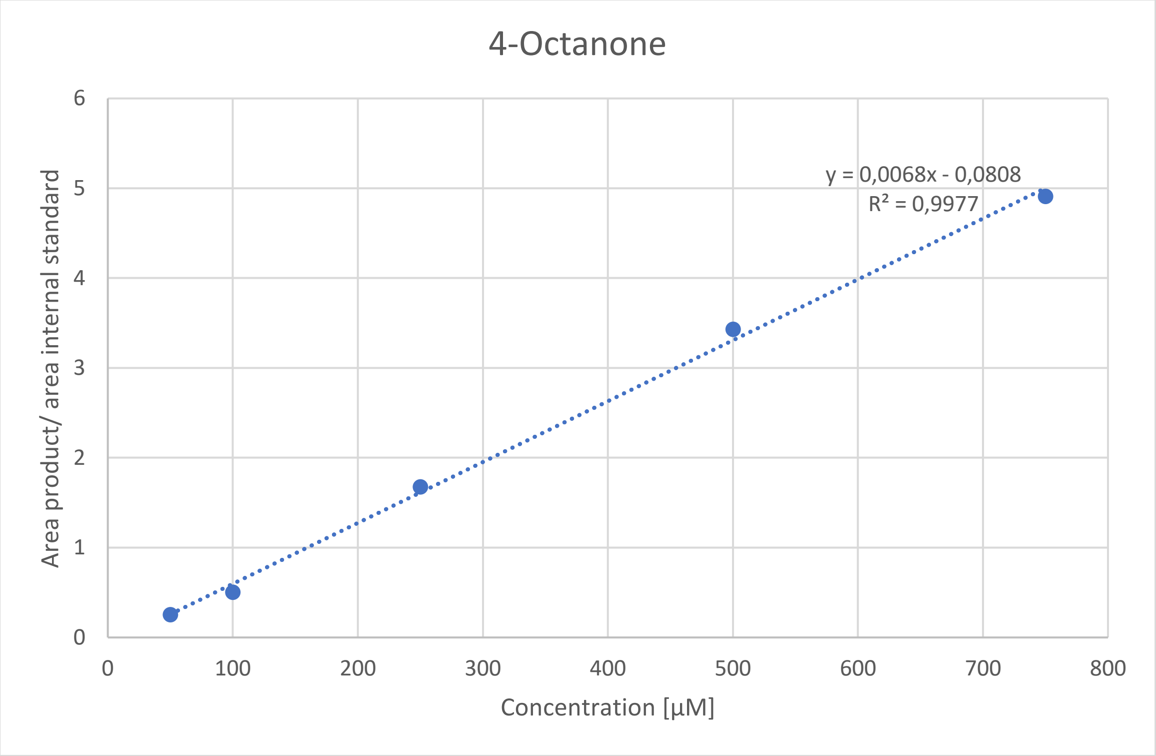


*Figure 12: Calibration curve of 4-Octanone.*

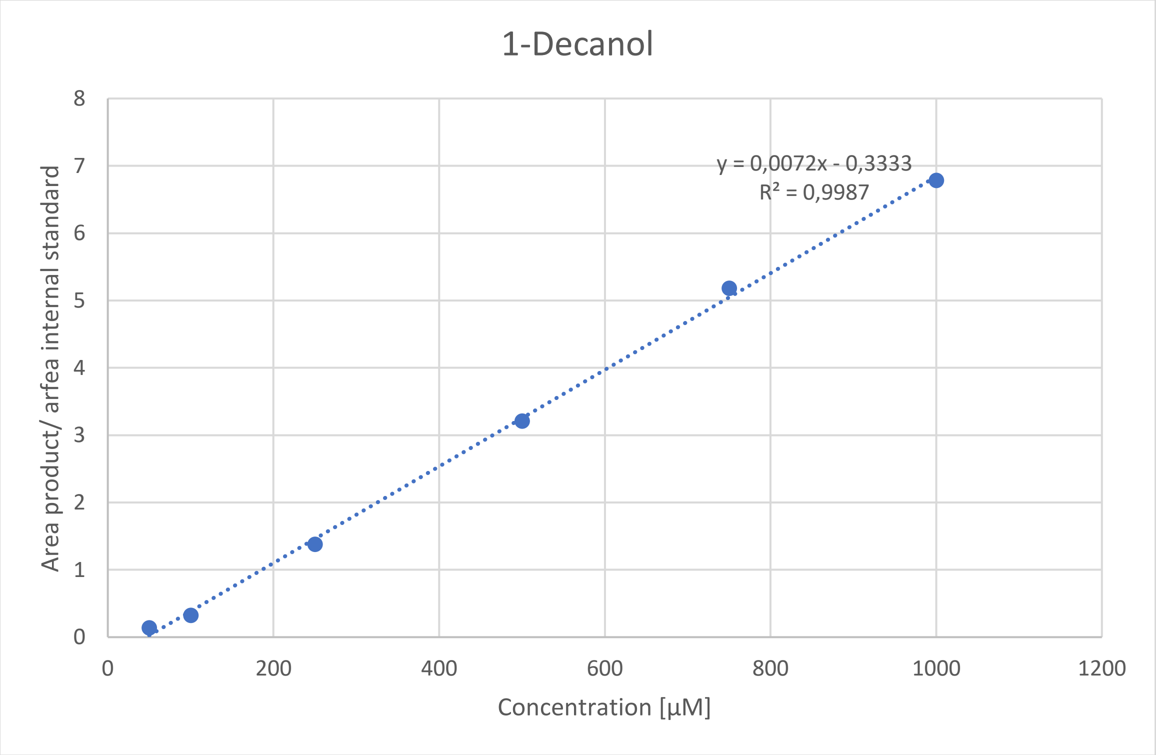


*Figure 13: Calibration curve of 1-Decanol.*

RESTRICTED

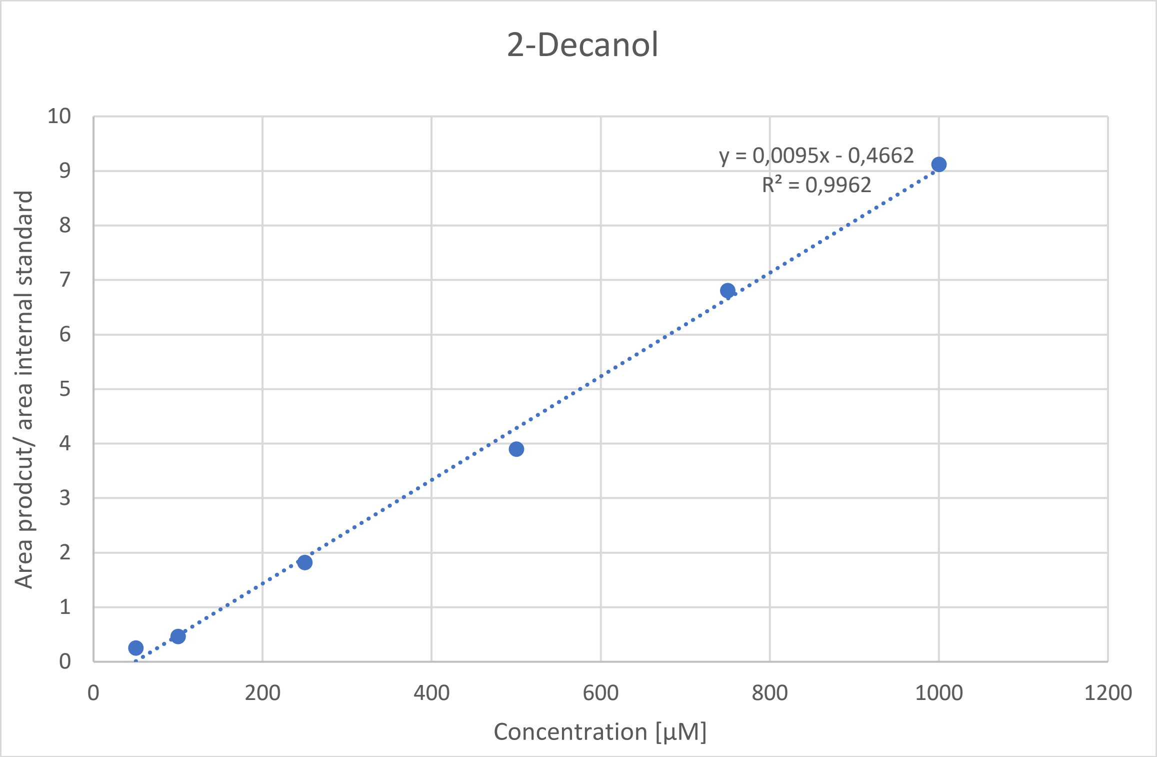


*Figure 14:Calibration curve of 2-Decanol.*

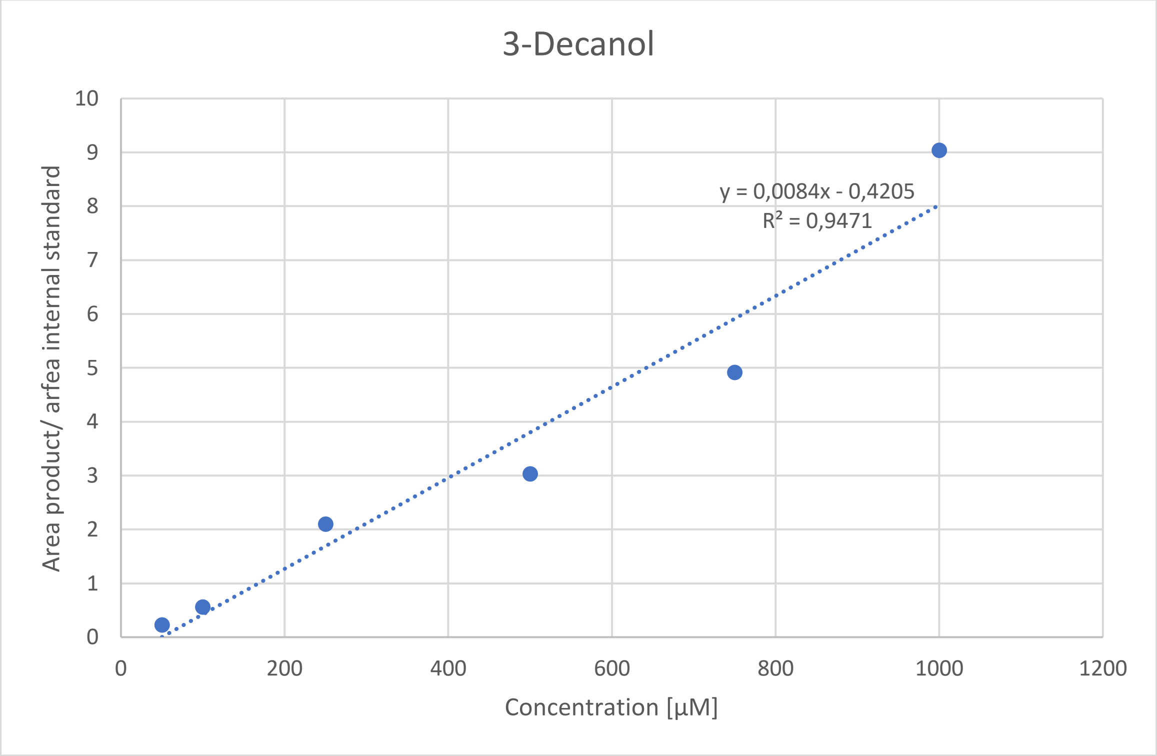


*Figure 15: Calibration curve of 3-Decanol.*

RESTRICTED

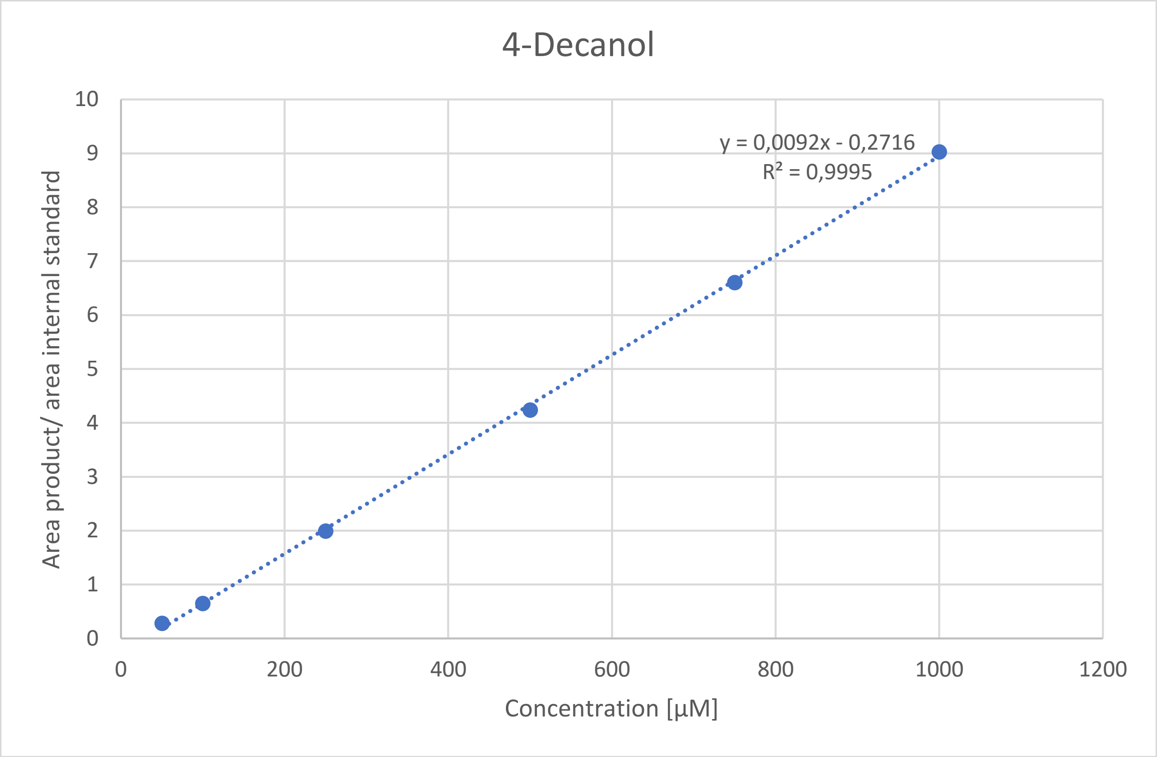


*Figure 16: Calibration curve of 4-Decanol.*

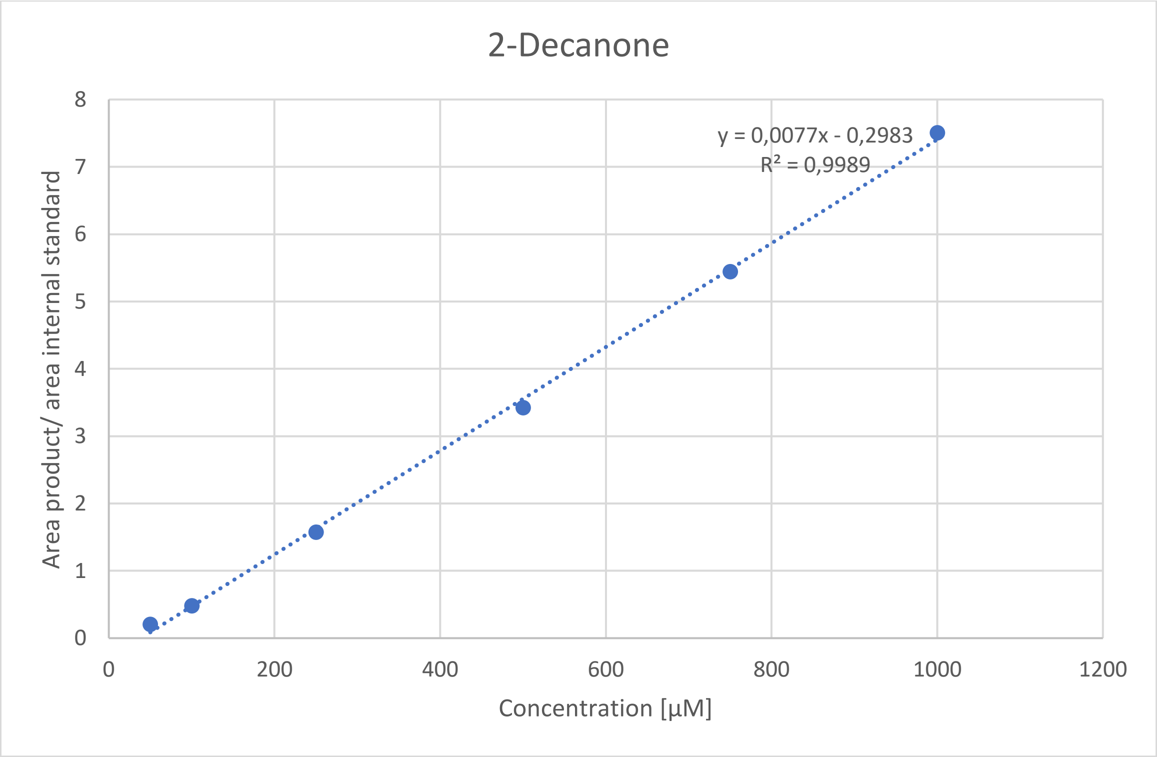


*Figure 17: Calibration curve of 2-Decanone.*

RESTRICTED

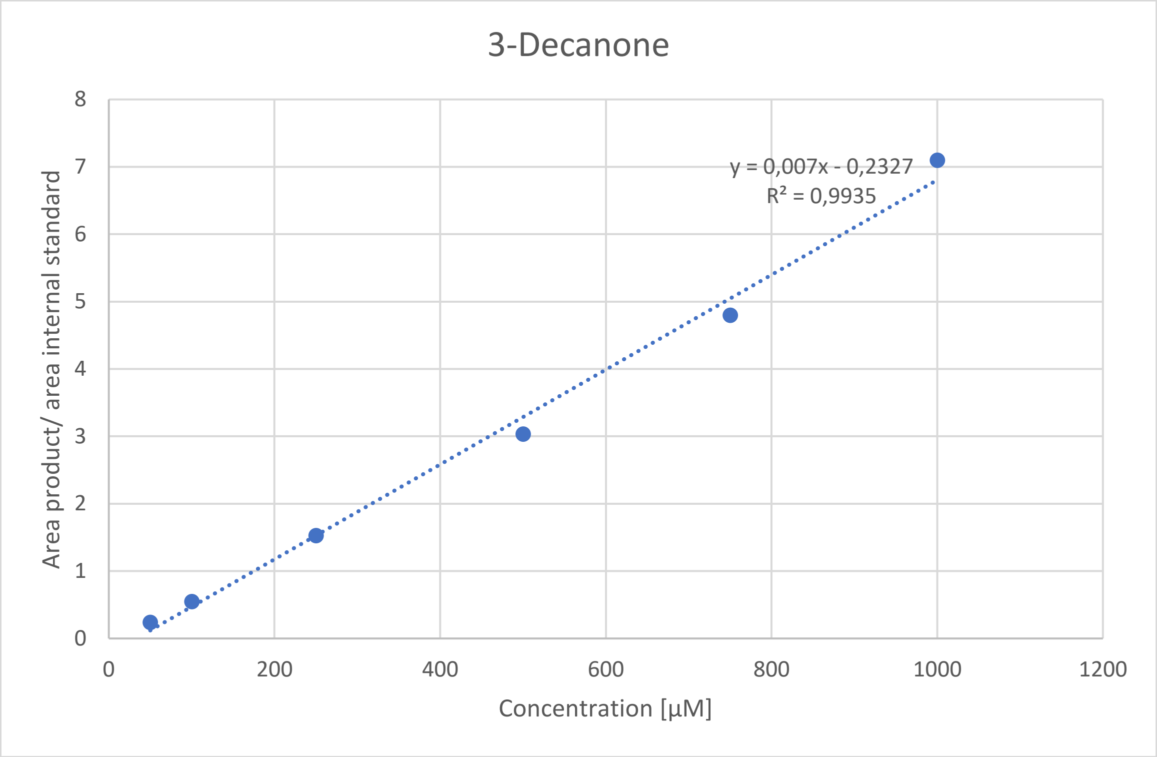


*Figure 18: Calibration curve of 3-Decanone.*

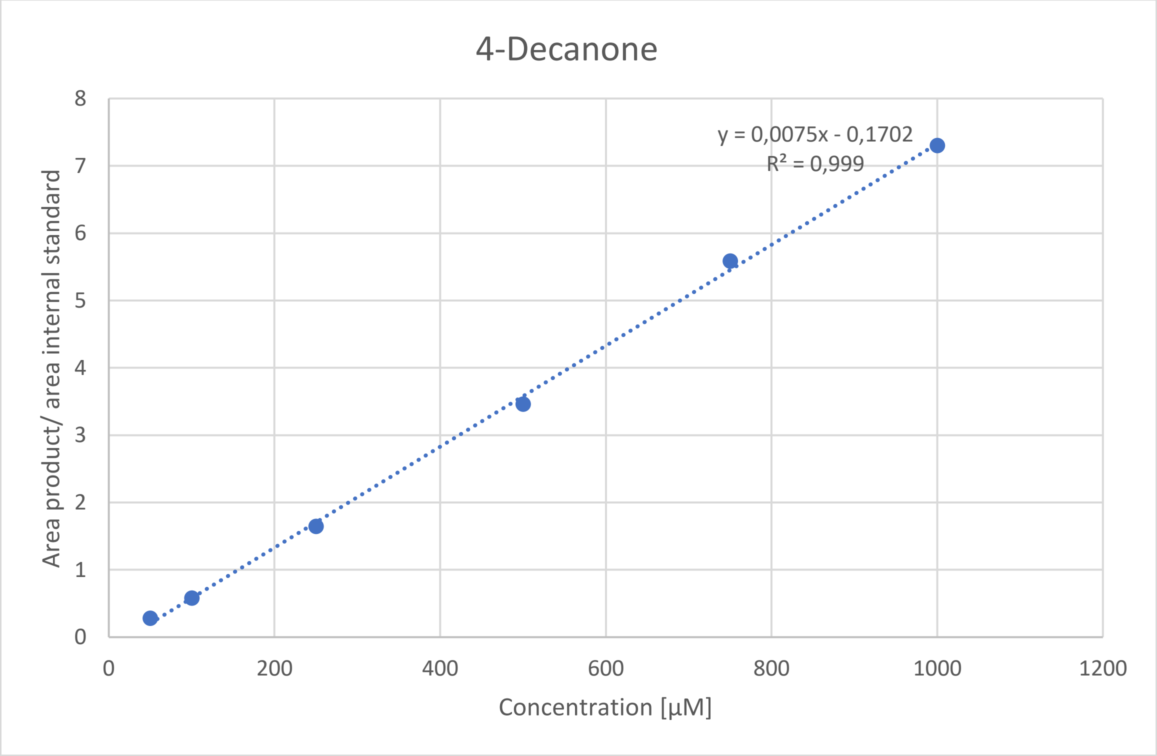


*Figure 19: Calibration curve of 4-Decanone.*

RESTRICTED

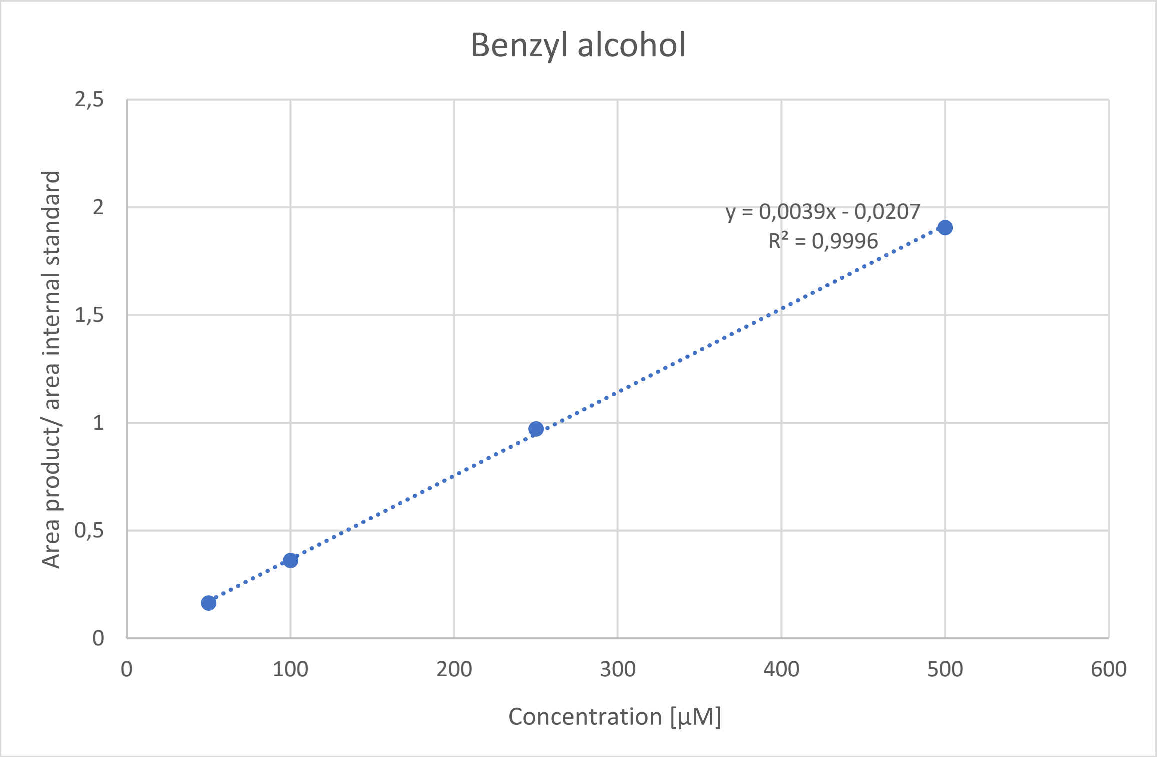


*Figure 20: Calibration curve of benzyl alcohol.*

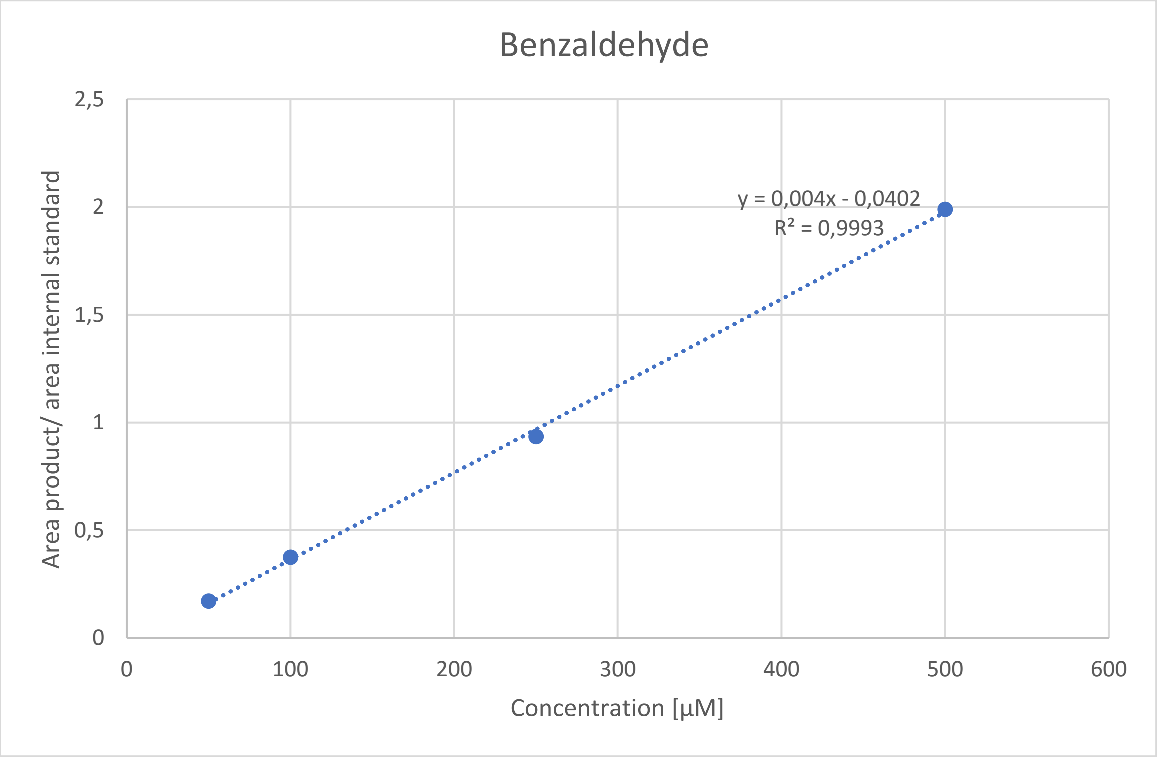


*Figure 21: Calibration curve of benzaldehyde.*

RESTRICTED

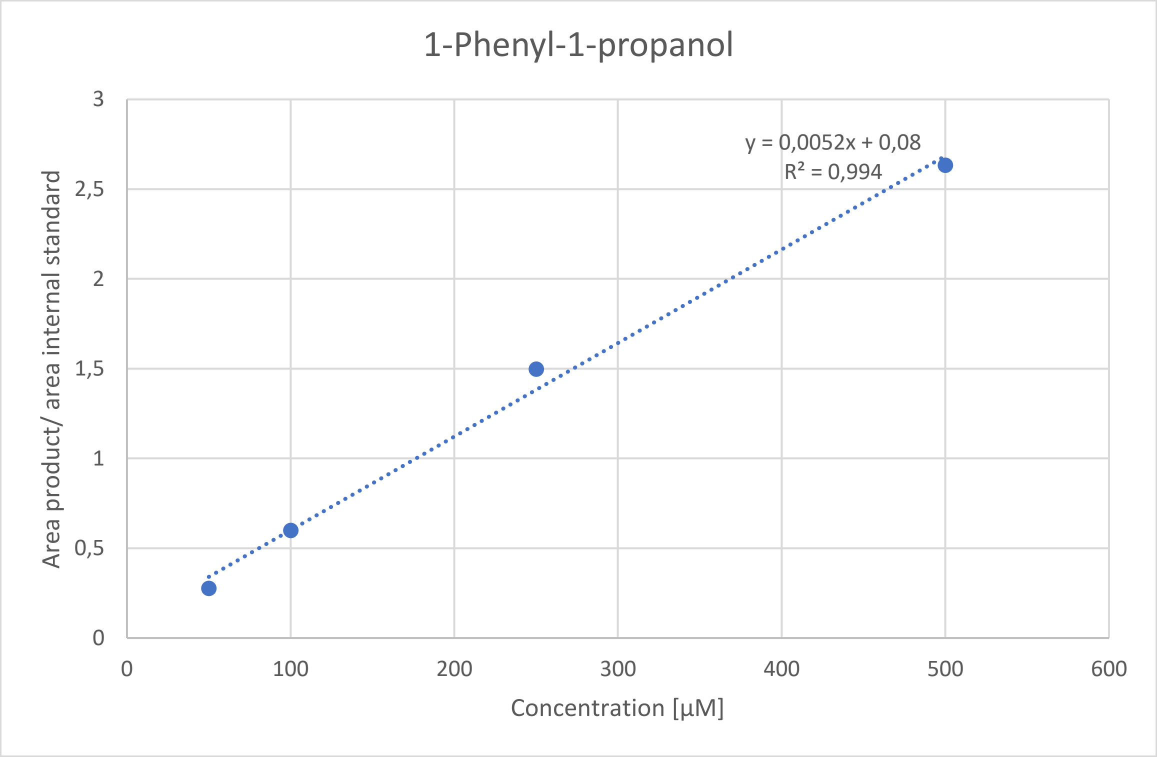


*Figure 22: Calibration curve of 1-phenyl-1-propanol.*

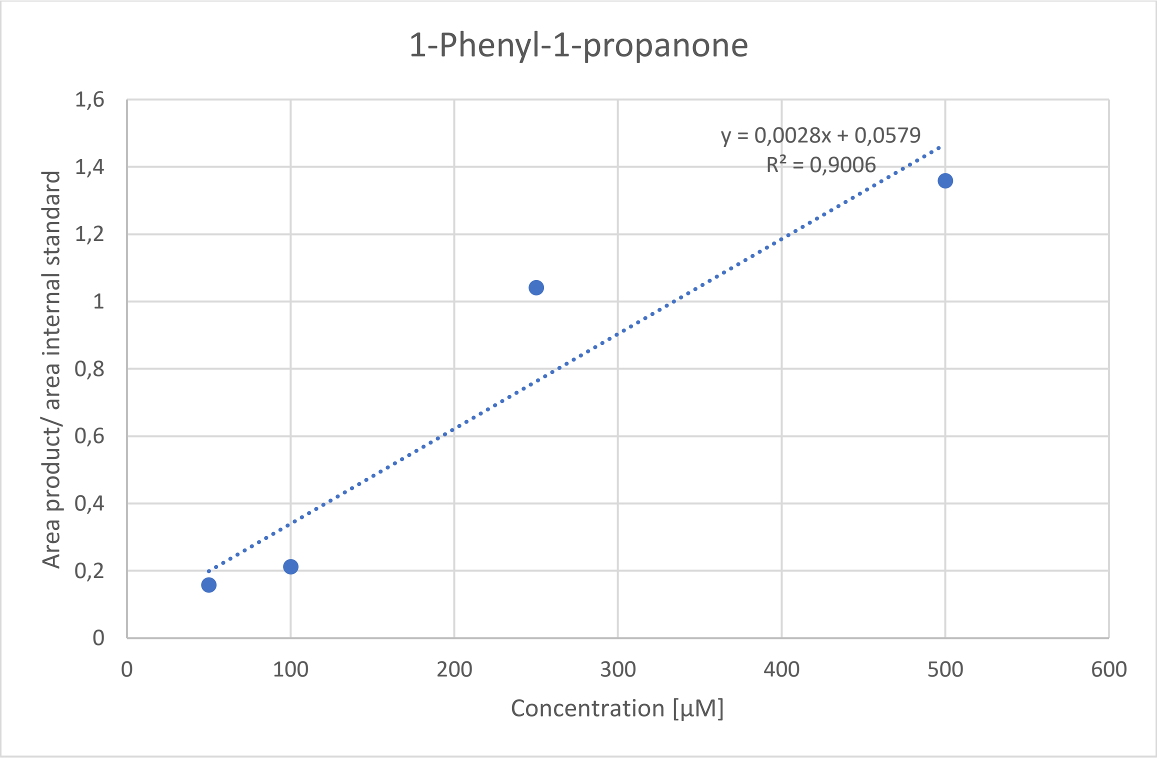


*Figure 23: Calibration curve of 1-phenyl-1-propanone.*

RESTRICTED

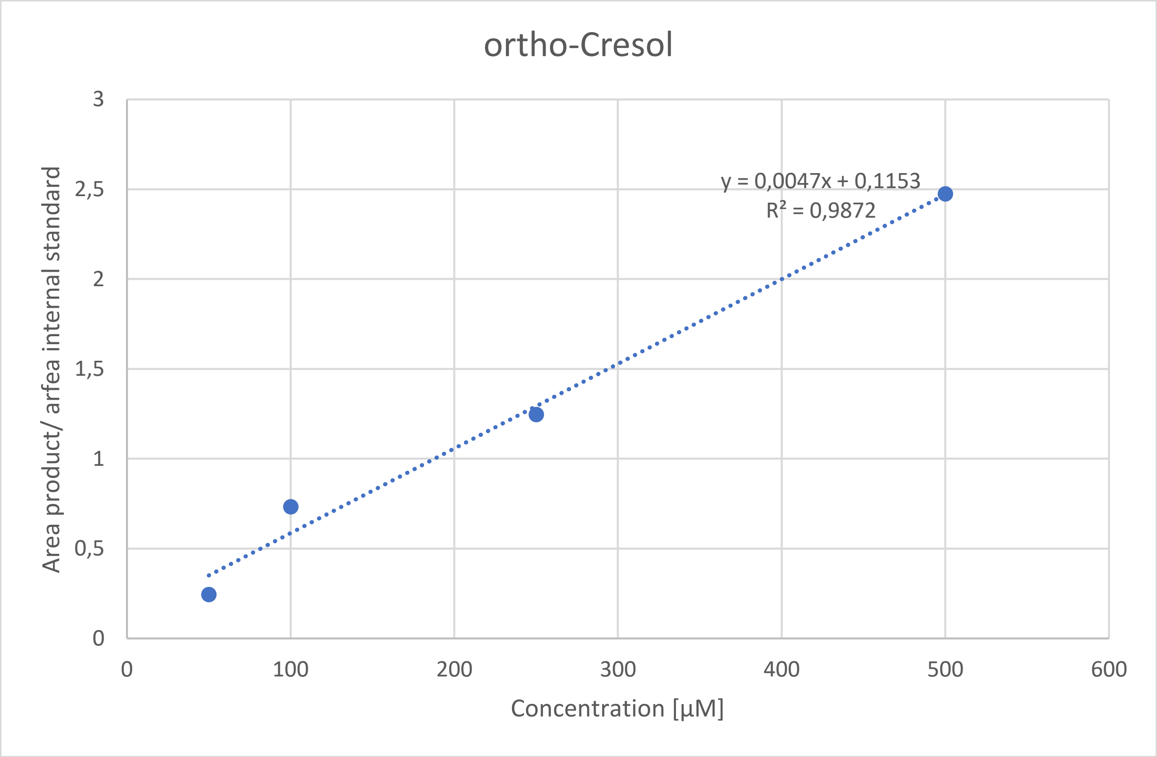


*Figure 24: Calibration curve of ortho-cresol.*

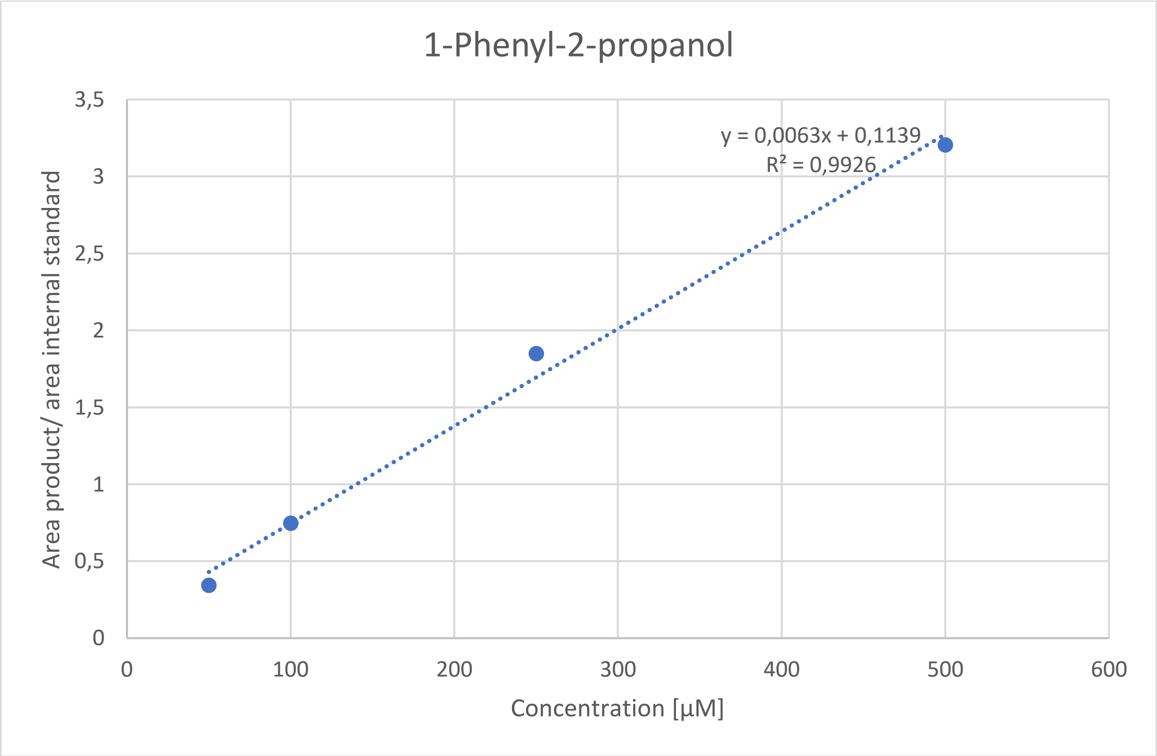


*Figure 25: Calibration curve of 1-phenyl-2-propanol.*

RESTRICTED

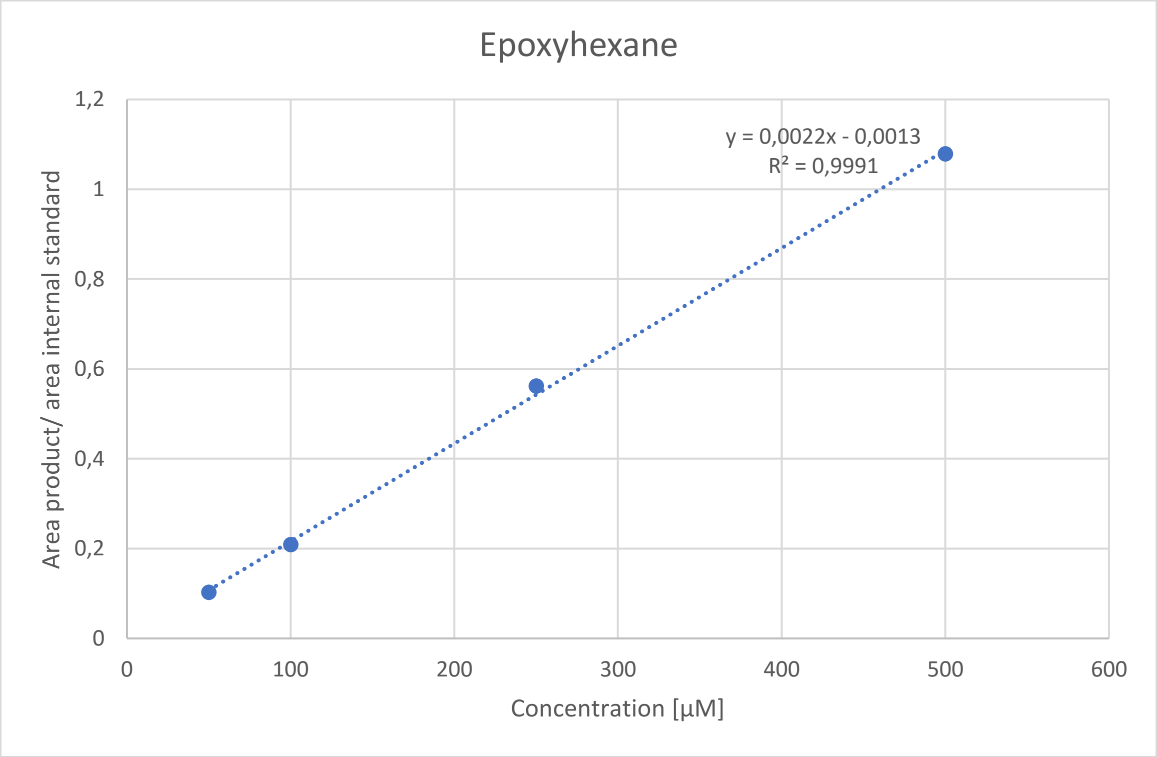


*Figure 26: Calibration curve of epoxyhexane.*

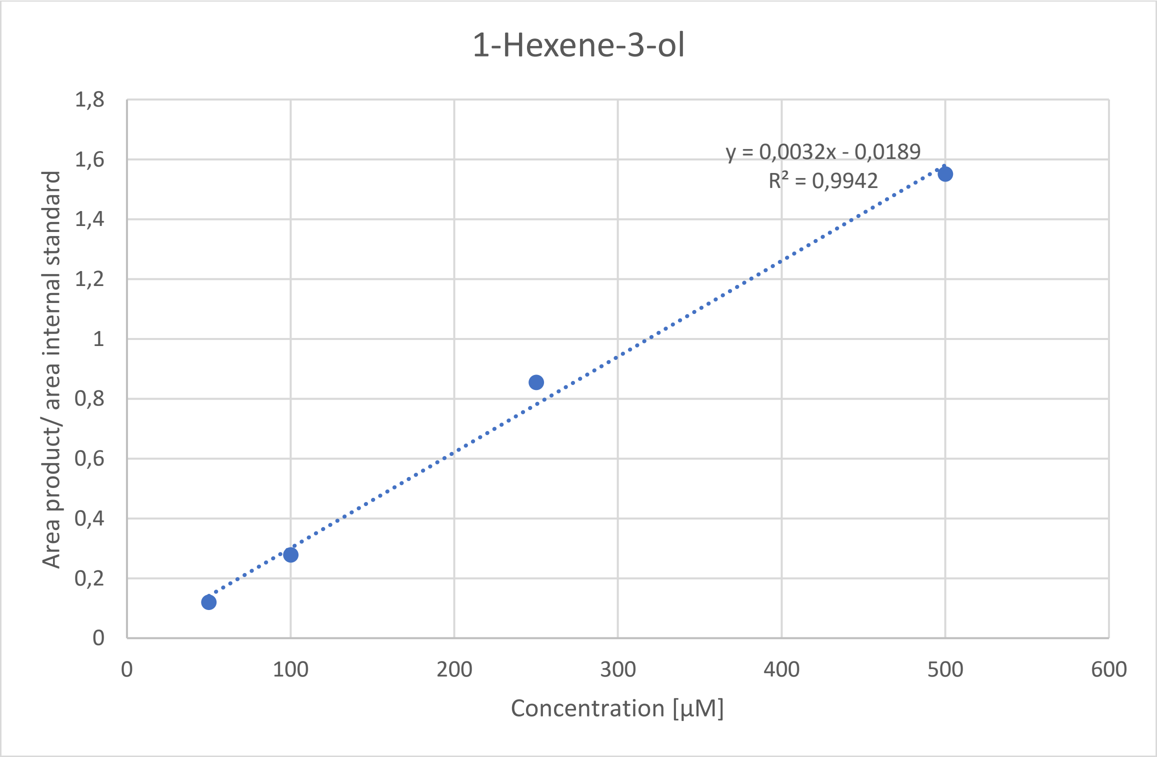


*Figure 27: Calibration curve of 1-hexene-3-ol.*

RESTRICTED

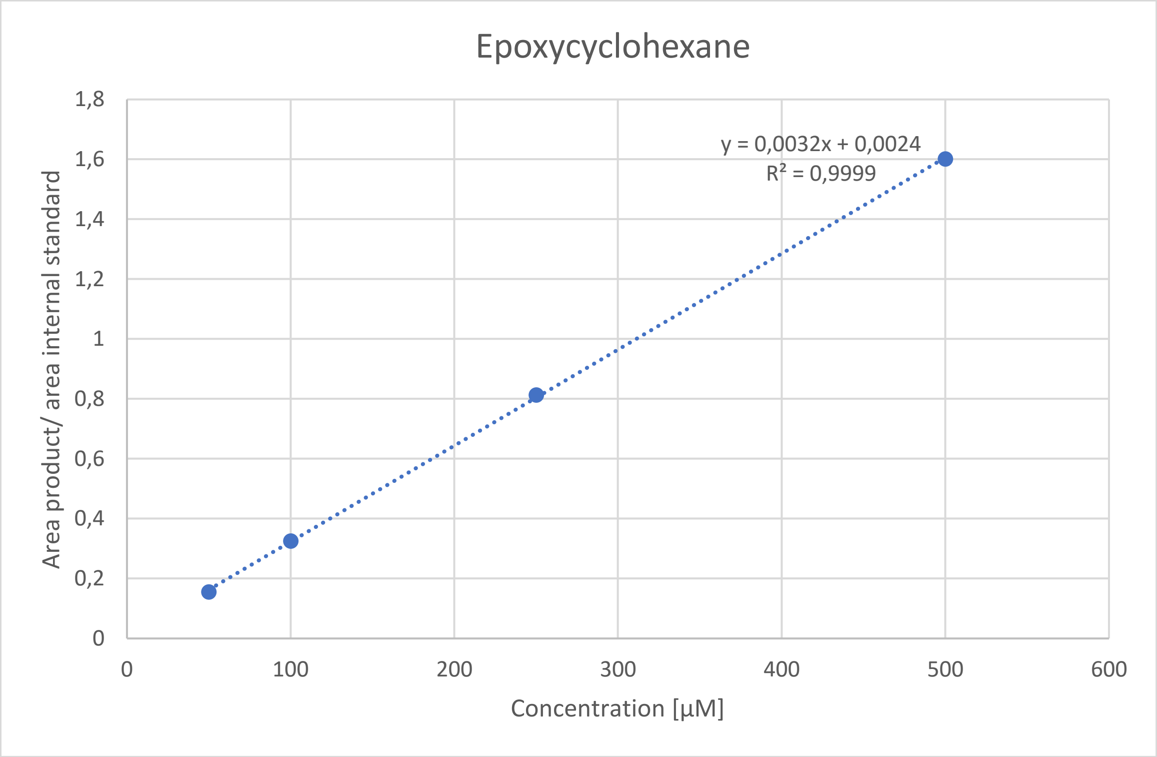


*Figure 28: Calibration curve of epoxycyclohexane.*

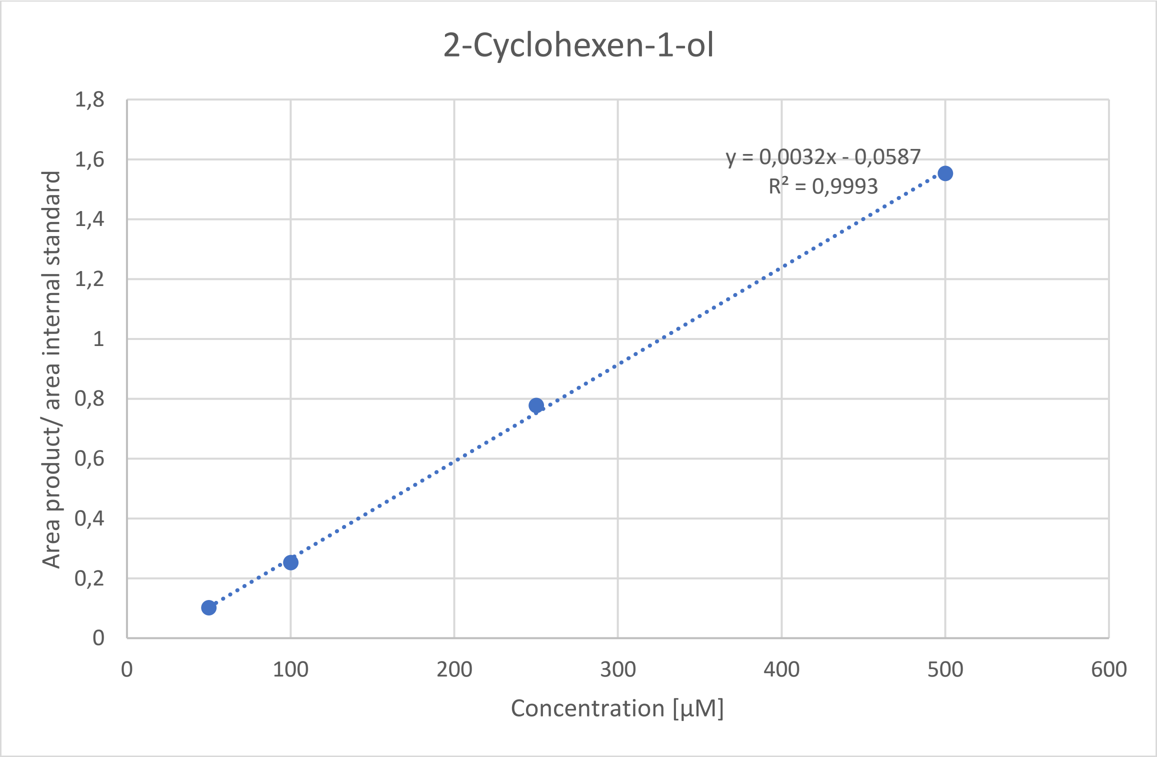


*Figure 29: Calibration curve of 2-cyclohexen-1-ol.*

RESTRICTED

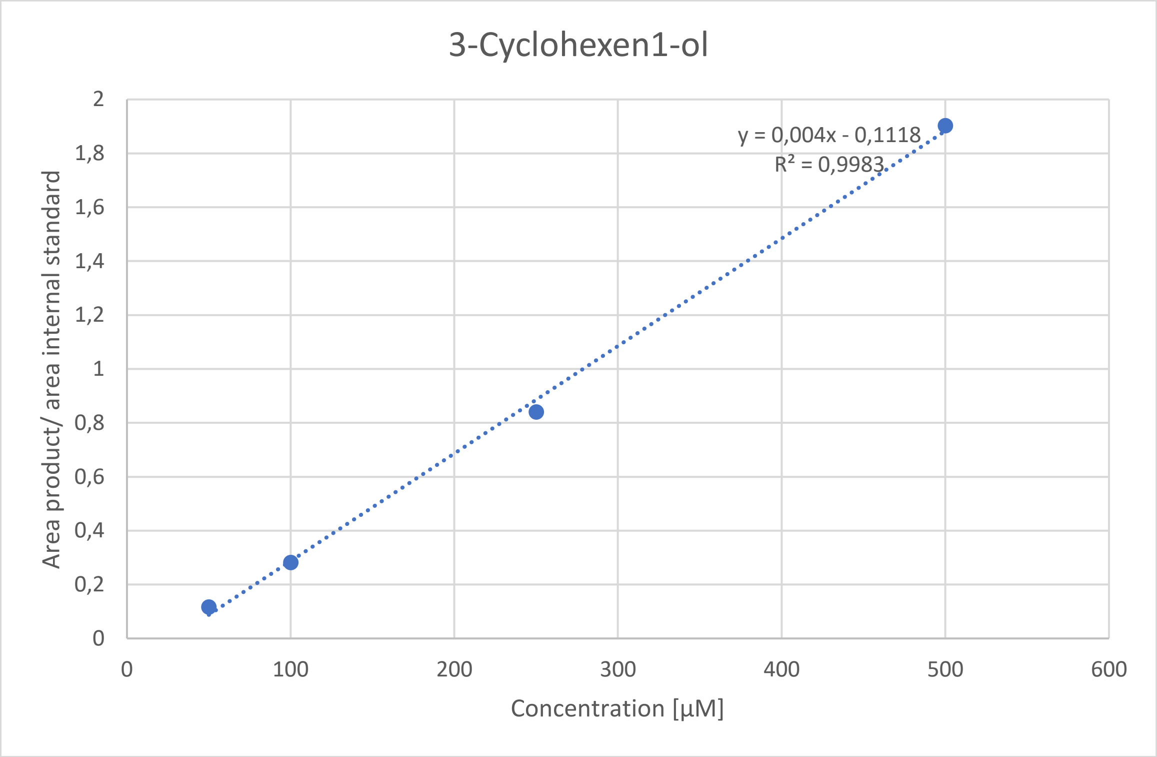


*Figure 30: Calibration curve of 3-cyclohexen-1-ol.*

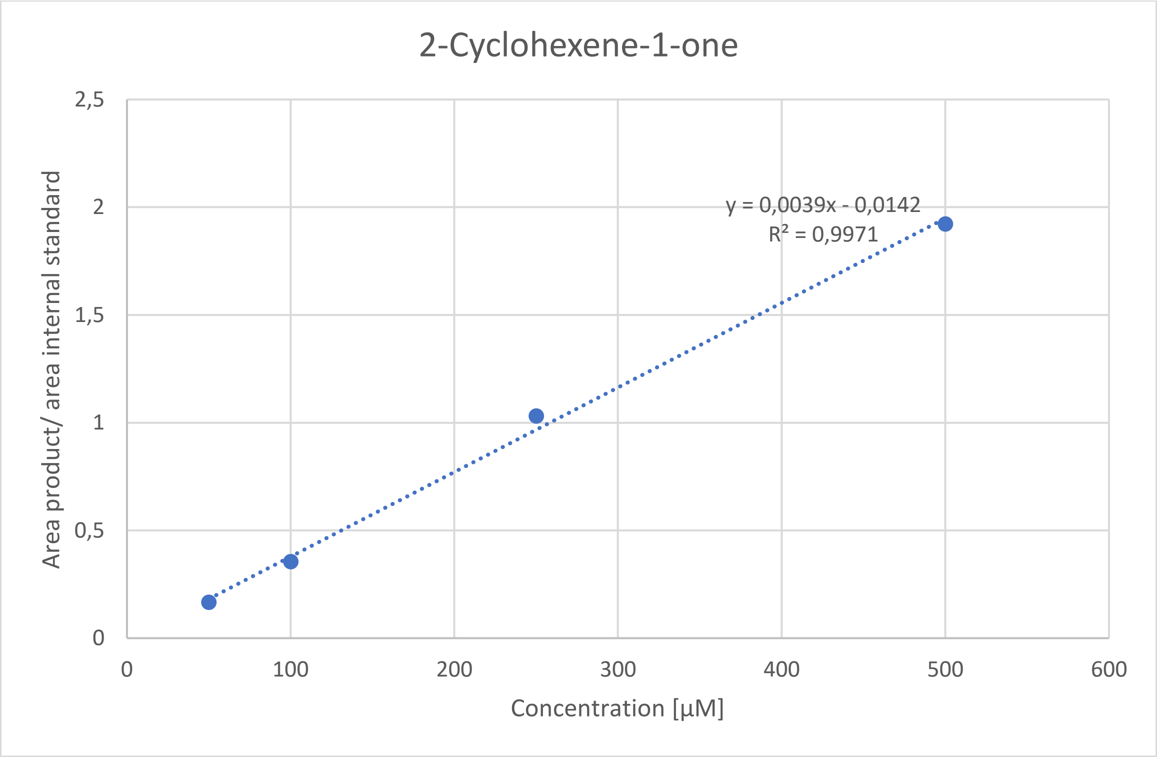


*Figure 31: Calibration curve of 2-cyclohexene-1-one.*

RESTRICTED

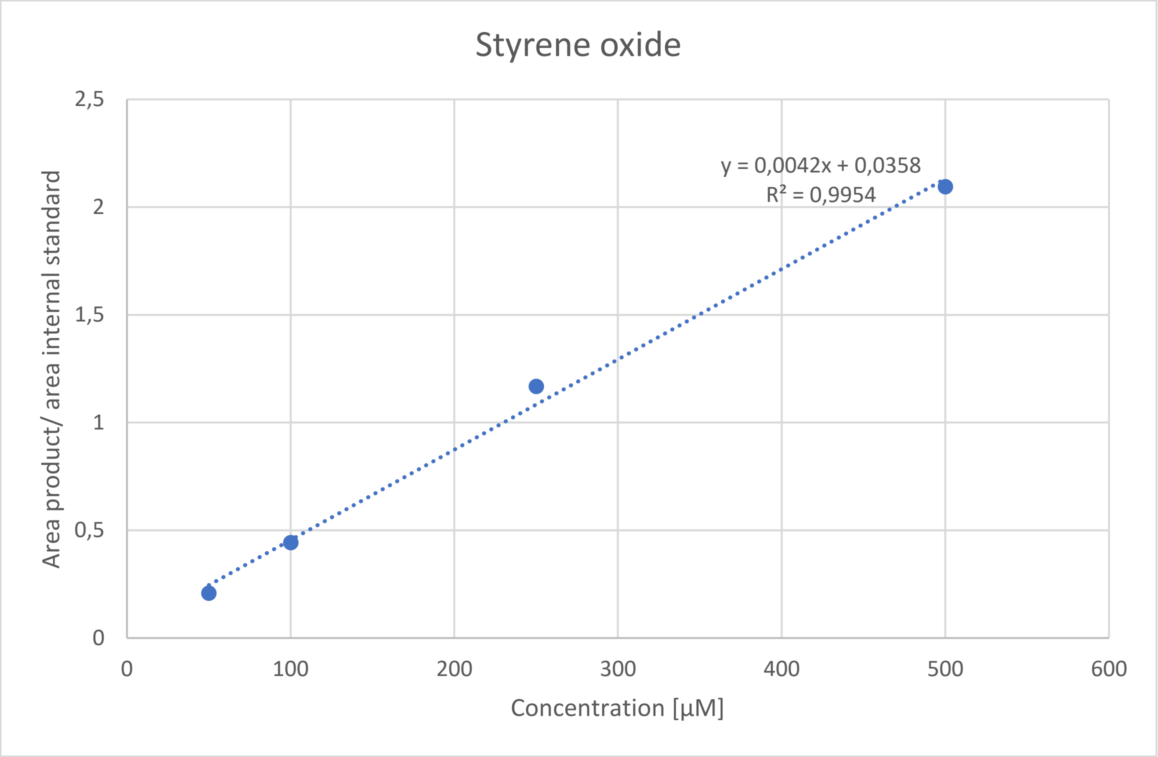


*Figure 32: Calibration curve of styrene oxide.*

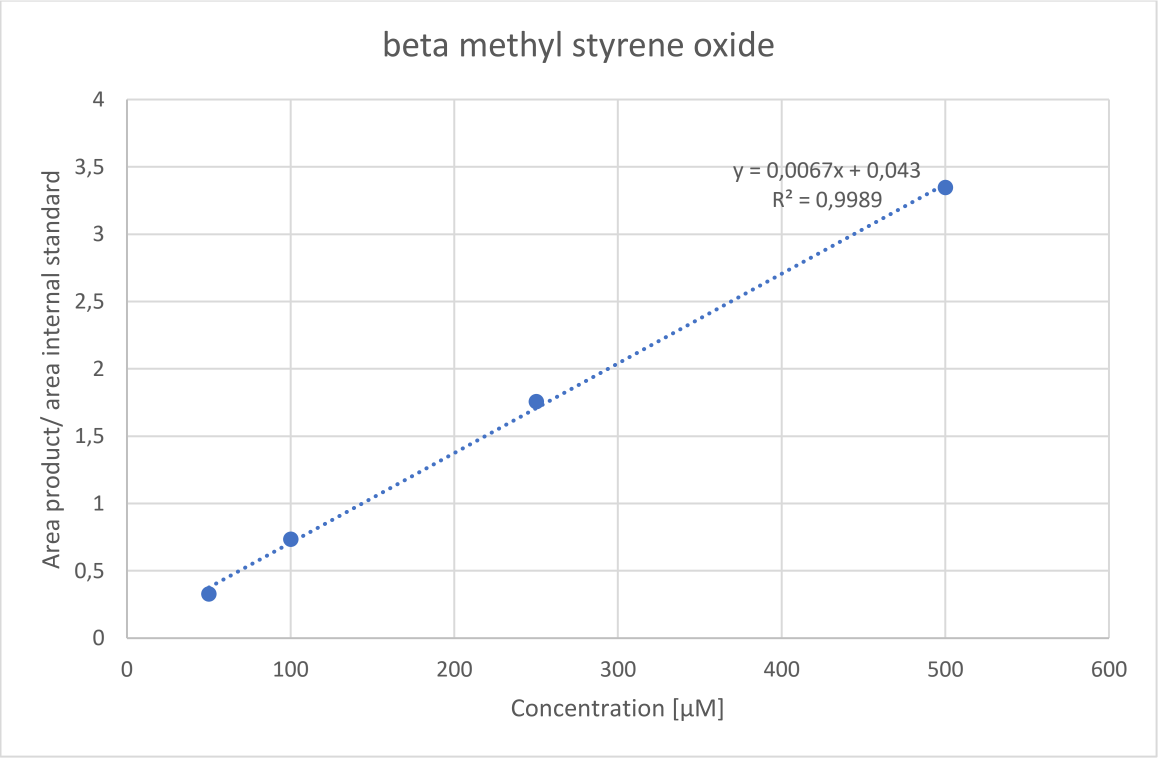


*Figure 33: Calibration curve of beta methyl styrene oxide.*

RESTRICTED

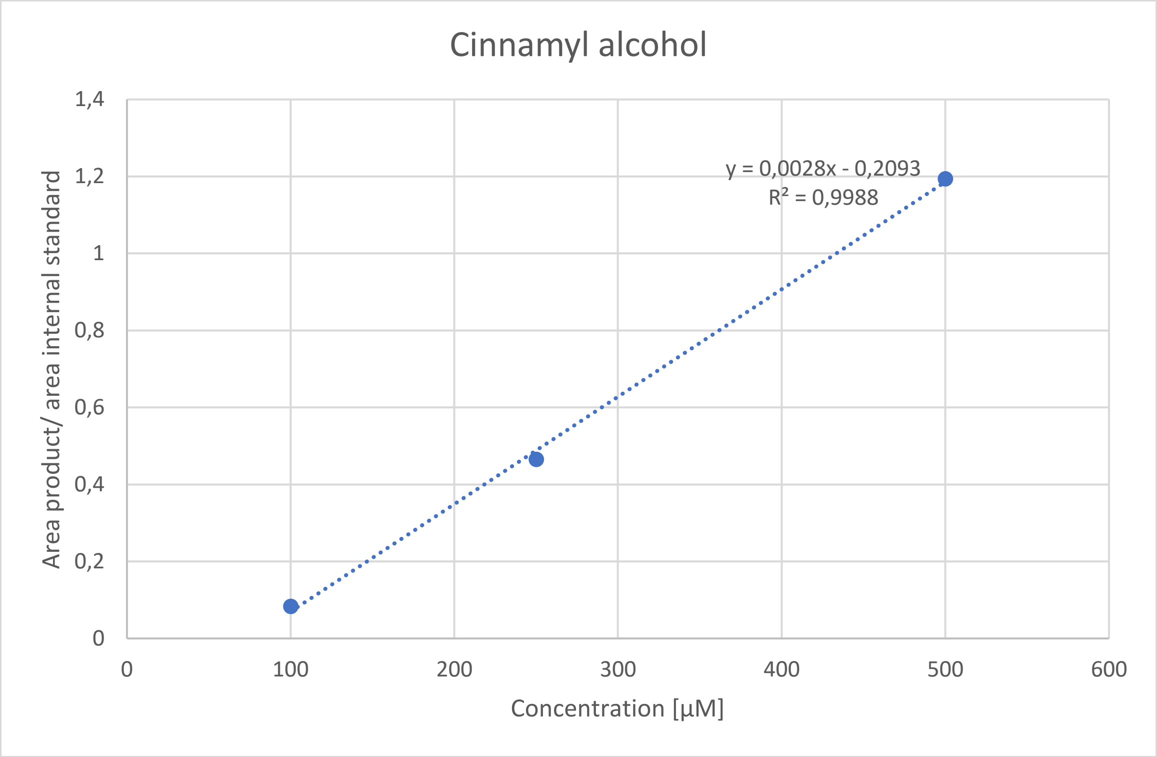


*Figure 34: Calibration curve of cinnamyl alcohol.*

**Linearity of different *m/z* values for different products.**

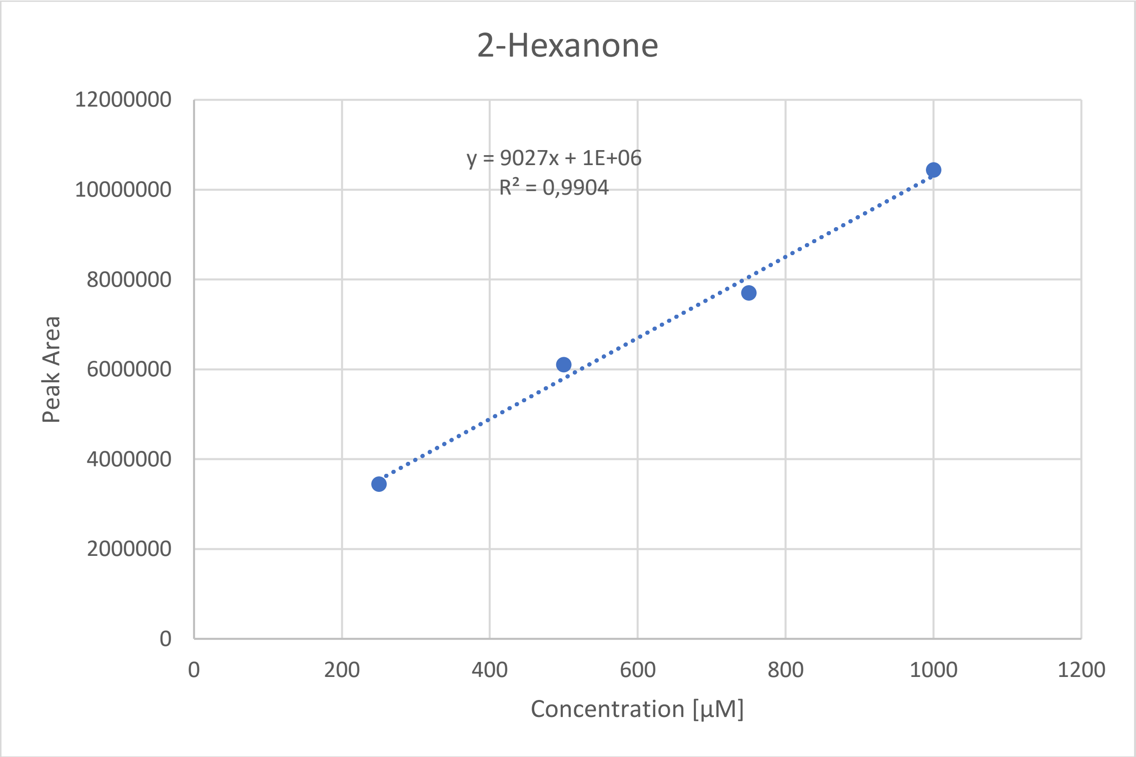


*Figure 35: Linearity of 2-Hexanone with the m/z of 43.*

RESTRICTED

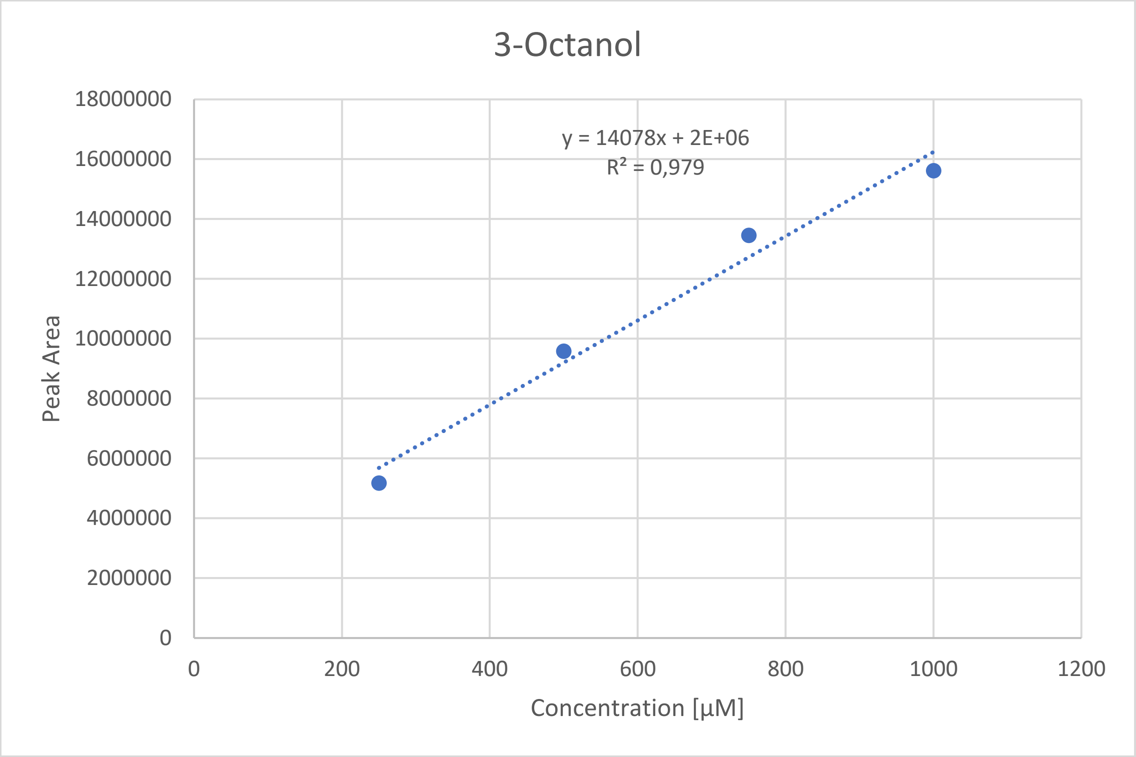


*Figure 36: Linearity of 3-Octanol with the m/z of 59.*

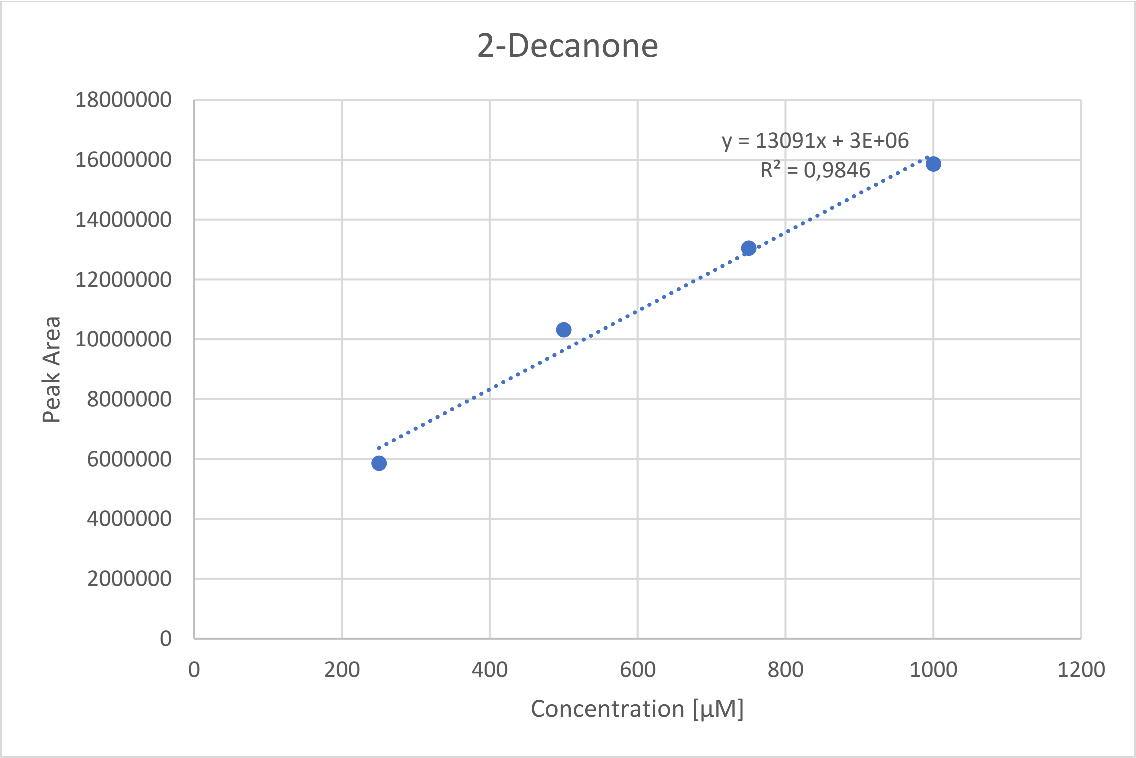


*Figure 37: Linearity of 2-Decanone with the m/z of 58.*

RESTRICTED

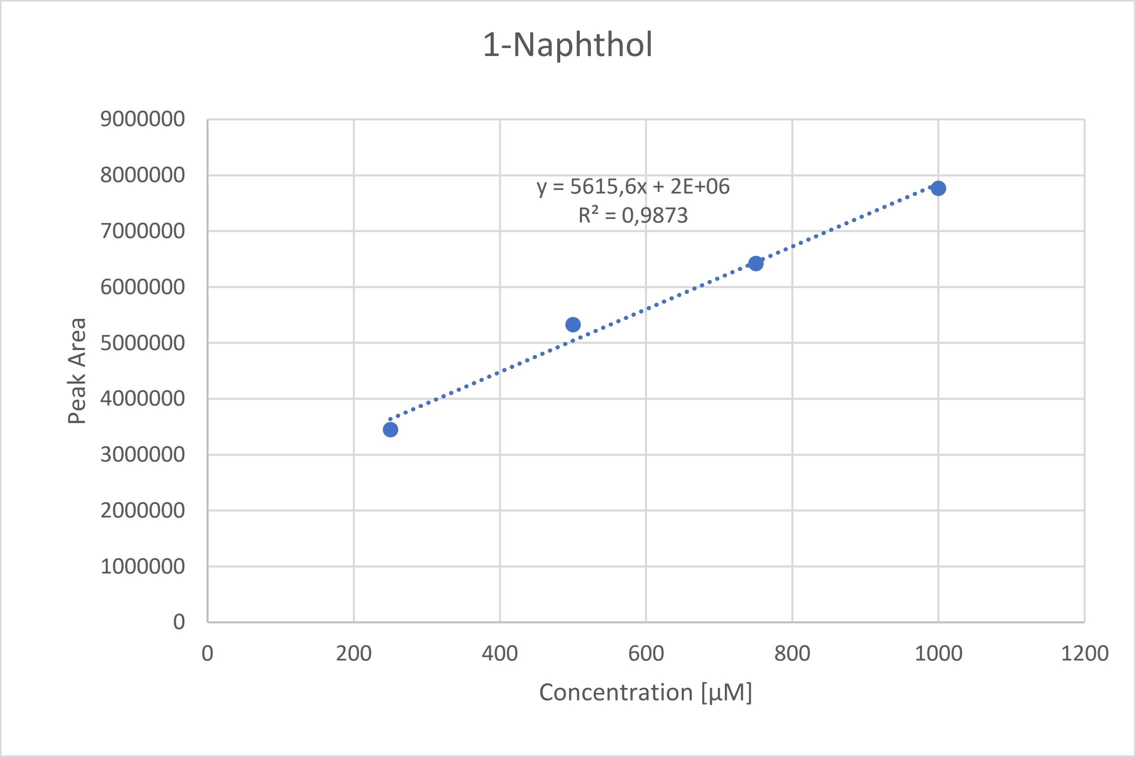


*Figure 38: Linearity of 1-Naphthol with the m/z of 144.*

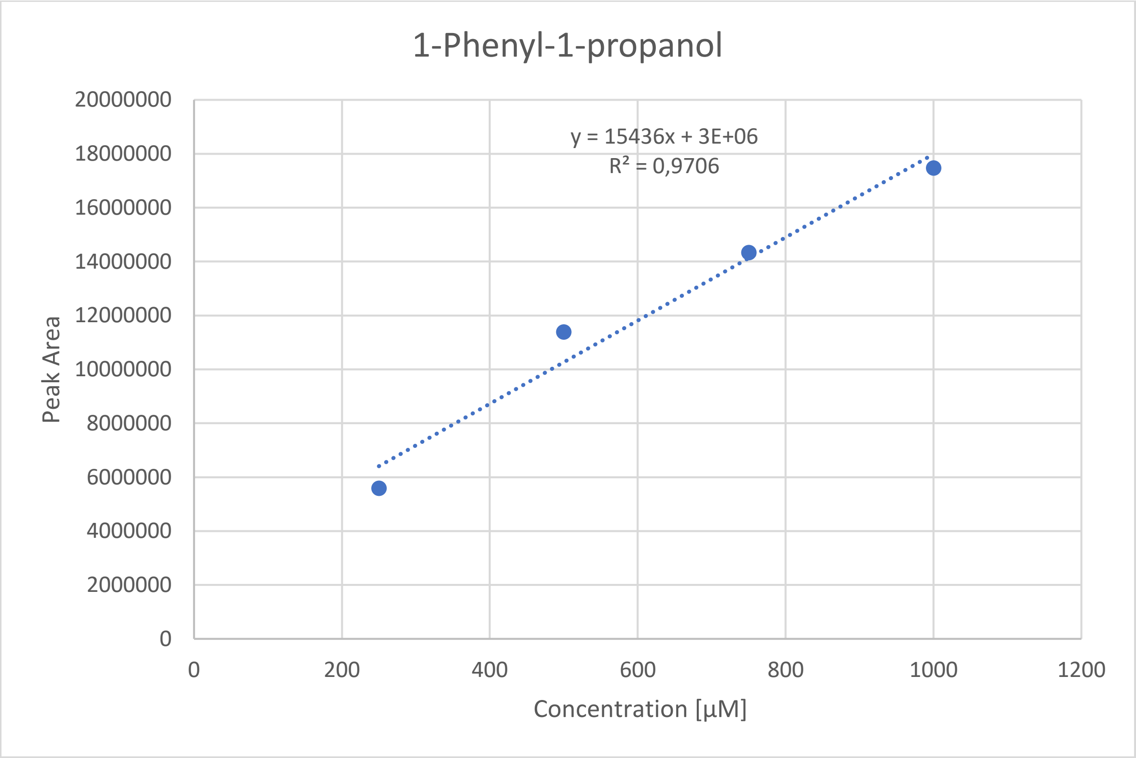


*Figure 39: Linearity of 1-Phenyl-1-propanol with the m/z of 107.*

RESTRICTED

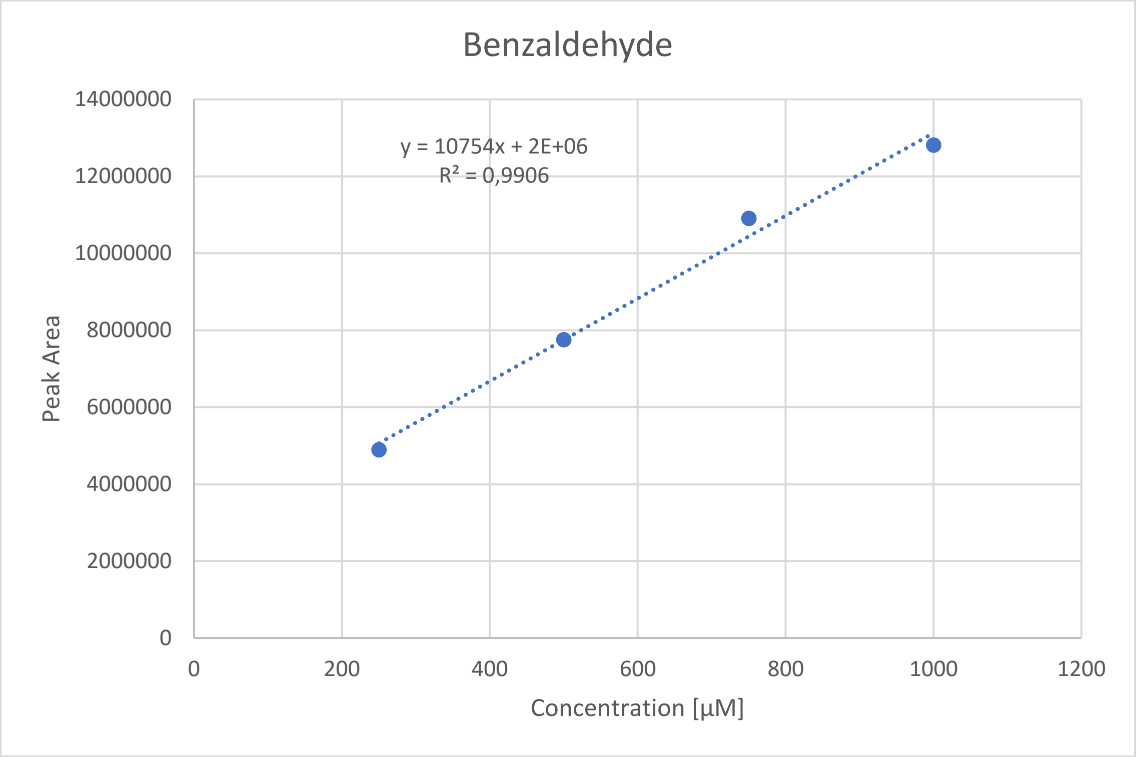


*Figure 40: Linearity of Benzaldehyde with the m/z of 77.*

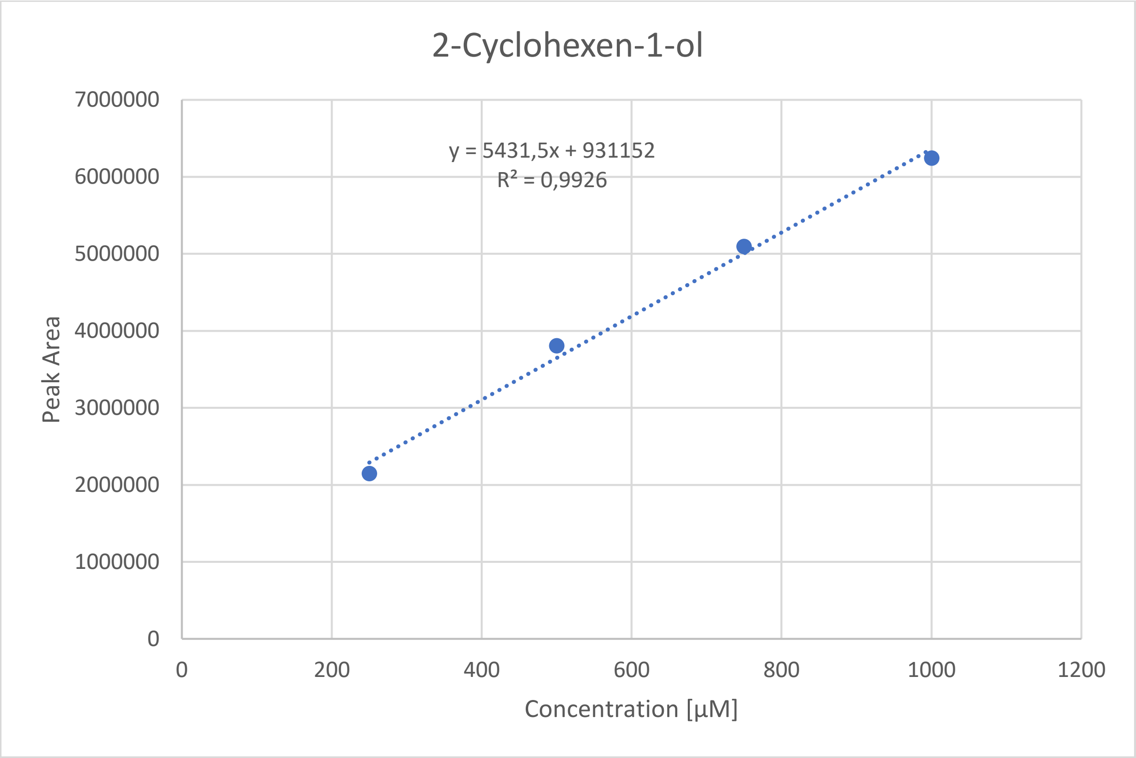


*Figure 41: Linearity of 2-Cyclohexen-1-ol with the m/z of 70.*

RESTRICTED

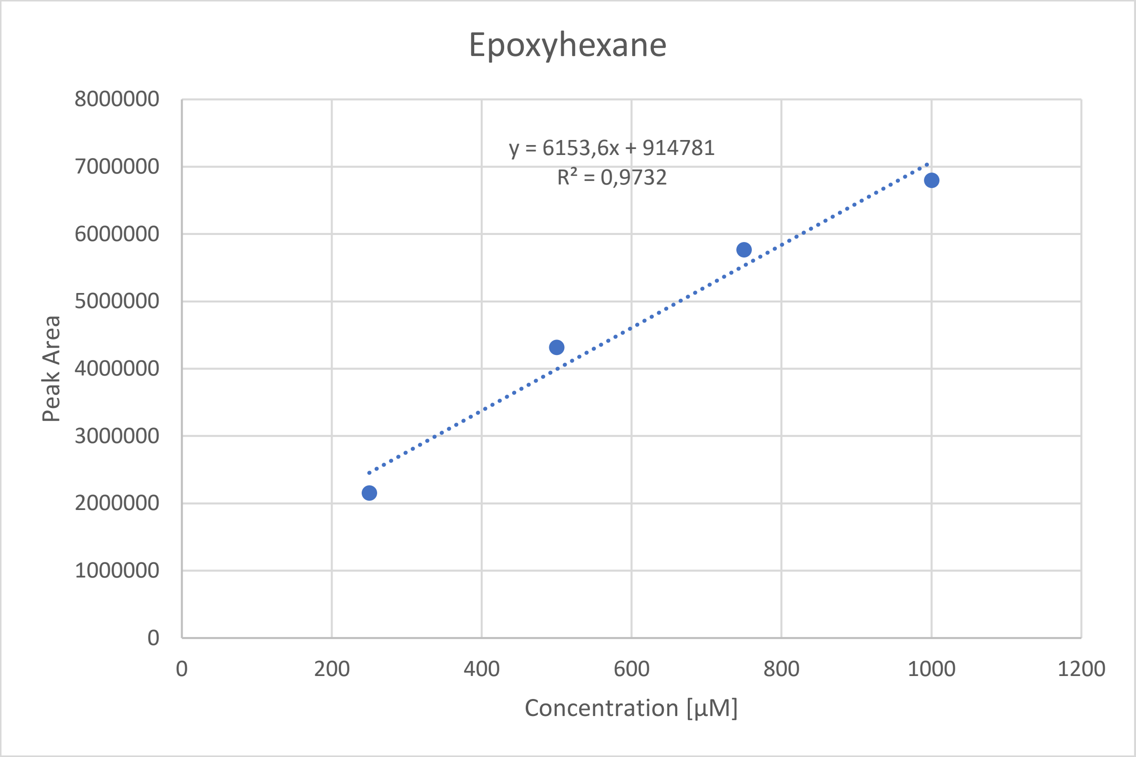


*Figure 42: Linearity of Epoxyhexane with the m/z of 71.*

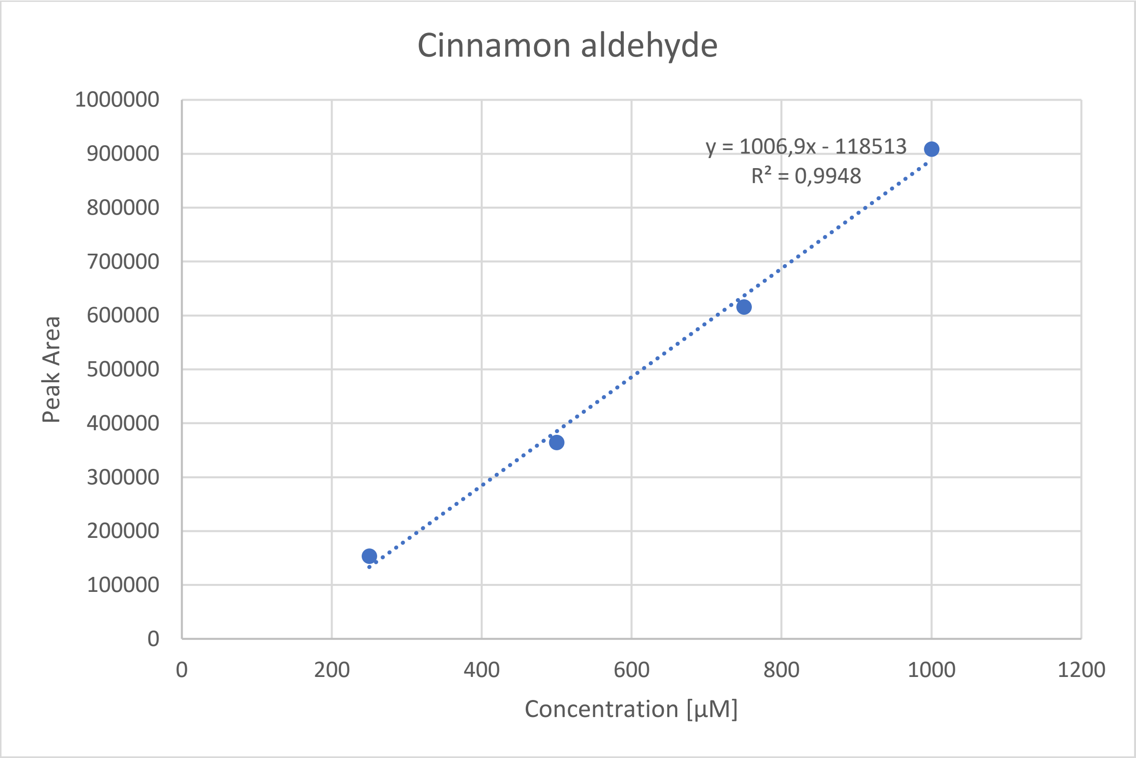


*Figure 43: Linearity of Cinnamon aldehyde with the m/z of 107.*

RESTRICTED

**Comparison of ionization of different products at the same concentration.**

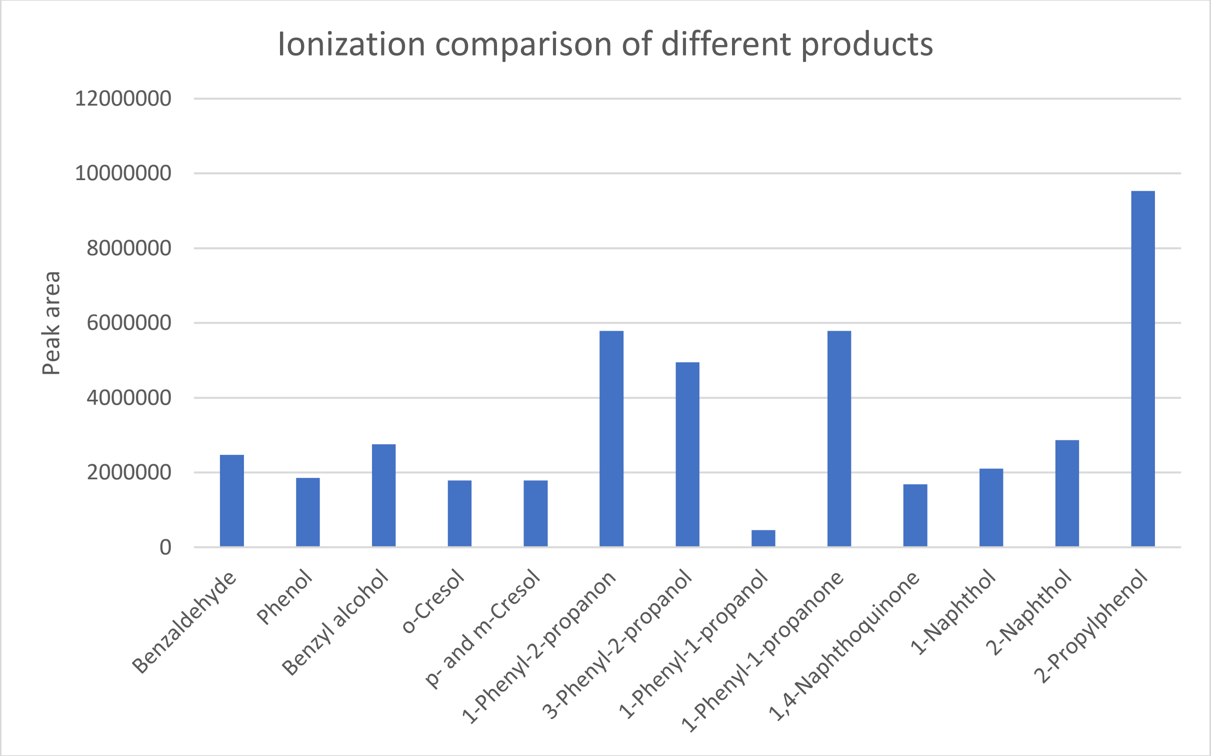


*Figure 44: Comparison of different products measured at the same concentration (500 µM) with GC-MS.*

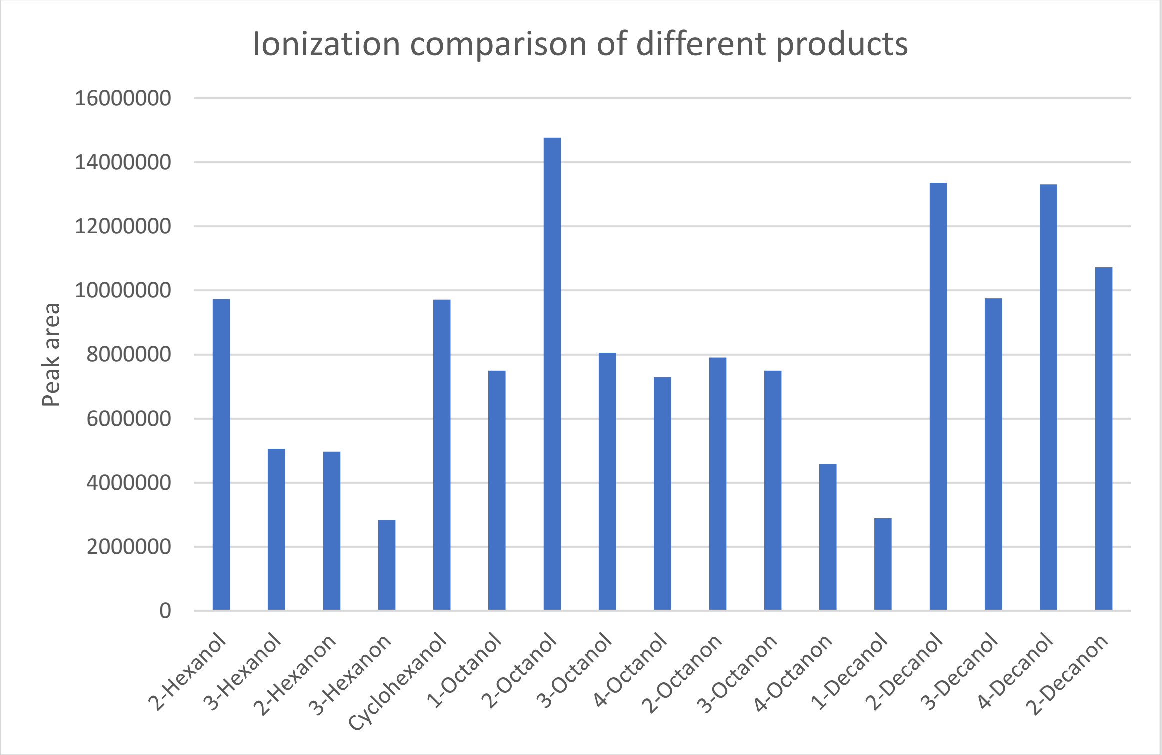


*Figure 45: Comparison of different products measured at the same concentration (500 µM) with GC-MS.*

RESTRICTED

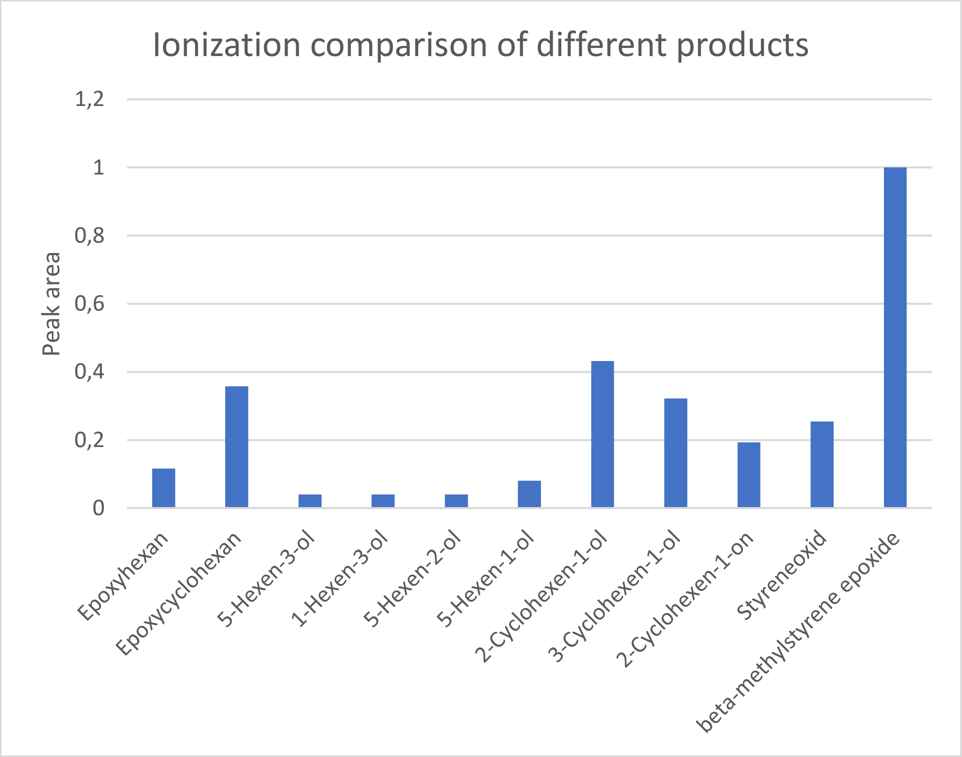


*Figure 46: Comparison of different products measured at the same concentration (500 µM) with GC-MS.*

**Comparison of long type vs short type UPOs for the hydroxylation of alkanes.**

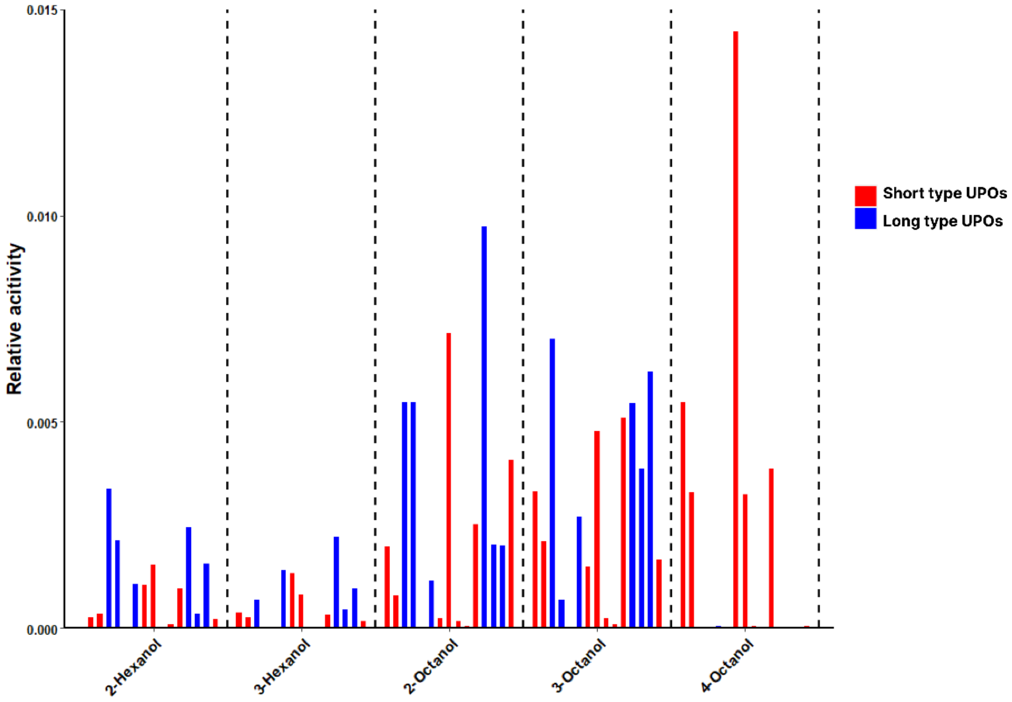


*Figure 47: Comparing long type and short type UPOs for the hydroxylation of alkanes; Part 1.*

RESTRICTED

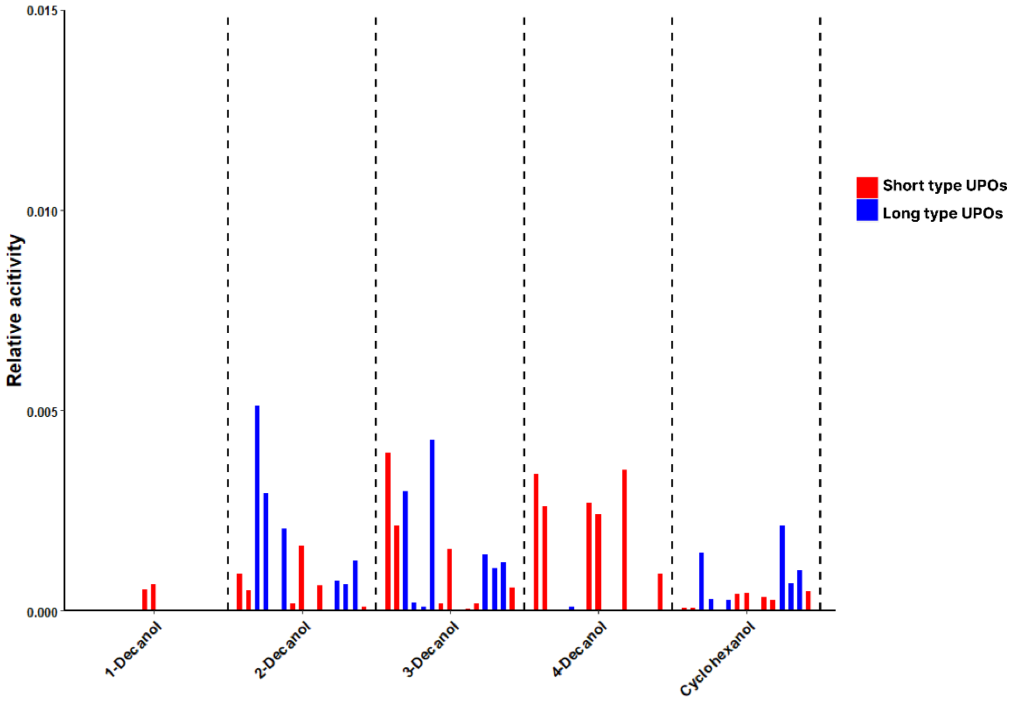


*Figure 48: Comparing long type and short type UPOs for the hydroxylation of alkanes; Part 2.*

### III. Supplementary Tables

##### T1. Overview of *m/z* used for different products in the screening.

*Table 1: m/z used to determine the products of the screening.*

| Product | *m/z* | |
|---|---|---|
| 2-hexanol | 45 | 69 |
| 3-hexanol | 59 | 55 |
| 2-hexanone | 43 | 58 |
| 3-hexanone | 43 | 57 |
| cyclohexanol | 57 | 82 |
| cyclohexanone | 56 | 98 |
| 2-octanol | 45 | 55 |
| 3-octanol | 59 | 83 |
| 4-octanol | 69 | 55 |
| 2-octanone | 58 | 43 |
| 3-octanone | 43 | 57 |
| 4-octanone | 71 | 57 |

RESTRICTED

| | | |
|---|---|---|
| 1-decanol | 55 | 56 |
| 2-decanol | 45 | 69 |
| 3-decanol | 69 | 59 |
| 4-decanol | 55 | 73 |
| 2-decanone | 58 | 43 |
| 3-decanone | 57 | 72 |
| 4-decanone | 43 | 71 |
| phenol | 94 | 66 |
| benzyl alcohol | 77 | 79 |
| benzaldehyde | 77 | 106 |
| *ortho*-cresol | 108 | 107 |
| *meta*-cresol | 108 | 107 |
| *para*-cresol | 107 | 108 |
| 1-phenyl-1-propanol | 107 | 79 |
| 1-phenyl-2-propanol | 92 | 91 |
| 1-phenyl-1-propanone | 105 | 77 |
| 1-phenyl-2-propanone | 43 | 91 |
| 2-propyplphenol | 107 | 136 |
| 1-naphthol | 144 | 115 |
| 2-naphthol | 144 | 115 |
| 1,4-naphthoquinone | 158 | 104 |
| 1,2-epoxyhexane | 71 | 42 |
| 1-hexen-3-ol | 57 | 29 |
| 1-hexen-5-ol | 67 | 54 |
| 1,2-epoxycylohexan | 83 | 41 |
| 2-cyclohexen-1-ol | 70 | 83 |
| 3-cyclohexen-1-ol | 54 | 70 |
| 2-cyclohexen-1-one | 68 | 42 |
| 2-phenyloxiran | 91 | 92 |
| β-methyl styrene epoxide | 90 | 91 |

**T2. Temperature programs for achiral measurements.**

*Table 2: Temperature programs for the screening of the substrate pools.*

| Products | Temperature program |
|---|---|
| Alkane substrate pool | 40 °C for 5 min |
| | 40 °C to 120 °C with 4 °C/min |

RESTRICTED

| | |
|---|---|
| Aromatics substrate pool | 60 °C to 120 °C with 4 °C/min<br>120 °C to 150 °C with 5 °C/min<br>150 °C to 160 °C with 4 °C/min |
| Epoxidation substrate pool | 30 °C for 5 min<br>30 °C to 120 °C with 10 °C/min<br>120 °C to 260 °C with 20 °C/min |

**T3. Temperature programs for chiral measurements.**

*Table 3: Temperature programs for the analysis of chiral products.*

| Products | Temperature program |
|---|---|
| 2-hexanol (derivatized) | 35 °C to 55 °C with 1 °C/min |
| 2-octanol (derivatized) | 35 °C to 45 °C with 1 °C/min<br>45 °C for 10 min<br>45 °C to 50 °C with 1 °C/min |
| 4-octanol (derivatized) | 35 °C to 45 °C with 1 °C/min<br>45 °C for 10 min<br>45 °C to 50 °C with 1 °C/min |
| 2-decanol (derivatized) | 55 °C for 100 min |
| 3-decanol (derivatized) | 55 °C for 100 min |
| 1-phenyl-1-propanol | 70 °C for 45 min<br>70 °C to 110 °C with 2.5 °C/min |
| 1-phenyl-2-propanol (derivatized)<br>*Cis* beta methyl styrene oxide | 100 °C for 9 min |

RESTRICTED

**T4. Regioselective Ratios from screening of alkane substrate pool. Results for hexane; alcohol and ketone were added up.**

*Table 4: Regioselective ratios from the screening of the alkane pool. The data was processed as described above.*

| Enzyme | 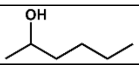 | 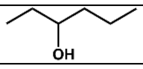 |
|---|---|---|
| *Tte*UPO | 0.549322529 | 0.450677471 |
| *Dca*UPO | 0.440378454 | 0.559621546 |
| *Mro*UPO | 1 | 0 |
| *Mfe*UPO | 0.651936981 | 0.348063019 |
| *Pab*UPO-I | 0.52478374 | 0.47521626 |
| *Pab*UPO-II | 0.433828256 | 0.566171744 |
| *Mth*UPO | 0.749353926 | 0.250646074 |
| *Aae*UPO* | 0.620736511 | 0.379263489 |
| *Mhi*UPO | 0 | 0 |
| *Anc*UPO-I | 0.400989468 | 0.599010532 |
| *Anc*UPO-II | 0.739938034 | 0.260061966 |
| *Chimera*UPO-I | 0.873311922 | 0.126688078 |
| *Chimera*UPO-II | 1 | 0 |
| *Chimera*UPO-III | 0 | 0 |
| *Chimera*UPO-IV | 0.431148379 | 0.568851621 |

**T5. Regioselective Ratios from screening of alkane substrate pool. Results for octane; alcohol and ketone were added up.**

*Table 5: Regioselective ratios from the screening of the alkane pool. The data was processed as described above.*

| Enzyme | 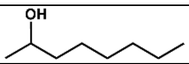 | 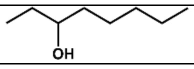 | 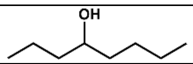 |
|---|---|---|---|
| *Tte*UPO | 0.76309769 | 0.232093294 | 0.004809016 |
| *Dca*UPO | 0.021350799 | 0.089106373 | 0.889542828 |
| *Mro*UPO | 0.285400012 | 0.714599988 | 0 |
| *Mfe*UPO | 0.536281994 | 0.279731999 | 0.183986007 |
| *Pab*UPO-I | 0.74079969 | 0.25920031 | 0 |
| *Pab*UPO-II | 0.450060055 | 0.549939945 | 0 |
| *Mth*UPO | 0.275042045 | 0.416639178 | 0.308318776 |
| *Aae*UPO* | 0.32501605 | 0.67498395 | 0 |
| *Mhi*UPO | 0.438973263 | 0.505889133 | 0.055137604 |

RESTRICTED

| | | | |
|---|---|---|---|
| *Anc*UPO-I | 0.226391838 | 0.294451933 | 0.479156229 |
| *Anc*UPO-II | 0.165634225 | 0.340402685 | 0.493963091 |
| *Chimera*UPO-I | 0.49082961 | 0.50917039 | 0 |
| *Chimera*UPO-II | 0.874441629 | 0.125558371 | 0 |
| *Chimera*UPO-III | 0 | 0 | 1 |
| *Chimera*UPO-IV | 0.297284415 | 0.702715585 | 0 |

**T6. Regioselective Ratios from screening of alkane substrate pool. Results for decane; alcohol and ketone were added up.**

*Table 6: Regioselective ratios from the screening of the alkane pool. The data was processed as described above.*

| Enzyme | 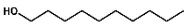 | 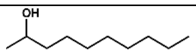 | 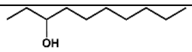 | 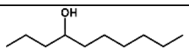 |
|---|---|---|---|---|
| *Tte*UPO | 0 | 0.033051044 | 0.37977448 | 0.587174475 |
| *Dca*UPO | 0.111117158 | 0.037051127 | 0.036292972 | 0.815538743 |
| *Mro*UPO | 0 | 0.912687857 | 0.087312143 | 0 |
| *Mfe*UPO | 0.070242177 | 0.278865591 | 0.280340547 | 0.370551685 |
| *Pab*UPO-I | 0 | 0.386782159 | 0.613217841 | 0 |
| *Pab*UPO-II | 0 | 0.411374176 | 0.588625824 | 0 |
| *Mth*UPO | 0 | 0 | 0.038693222 | 0.961306778 |
| *Aae*UPO* | 0 | 0.54902182 | 0.45097818 | 0 |
| *Mhi*UPO | 0 | 0.469985616 | 0 | 0.530014384 |
| *Anc*UPO-I | 0 | 0.11087179 | 0.487553294 | 0.401574916 |
| *Anc*UPO-II | 0 | 0.090074209 | 0.429085994 | 0.480839797 |
| *Chimera*UPO-I | 0 | 0.674212756 | 0.321031897 | 0.004755347 |
| *Chimera*UPO-II | 0 | 0.94549843 | 0.05450157 | 0 |
| *Chimera*UPO-III | 0 | 0.011624729 | 0.513311081 | 0.47506419 |
| *Chimera*UPO-IV | 0 | 0.35247392 | 0.646553106 | 0.000972975 |

**T7. Regioselective Ratios from screening of aromatics substrate pool. Results for toluene; alcohol and ketone were added up.**

RESTRICTED

*Table 7: Regioselective ratios from the screening of the aromatics pool. The data was processed as described above.*

| Enzyme | 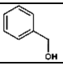 | 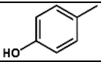 |
|---|---|---|
| *Tte*UPO | 0.964406372 | 0.035593628 |
| *Dca*UPO | 0.971764608 | 0.028235392 |
| *Mro*UPO | 0.984366915 | 0.015633085 |
| *Mfe*UPO | 0.847277722 | 0.152722278 |
| *Pab*UPO-I | 0.739092563 | 0.260907437 |
| *Pab*UPO-II | 0.34015734 | 0.65984266 |
| *Mth*UPO | 0.887467593 | 0.112532407 |
| *Aae*UPO* | 0.73975047 | 0.26024953 |
| *Mhi*UPO | 0.994993046 | 0.005006954 |
| *Anc*UPO-I | 0.988581603 | 0.011418397 |
| *Anc*UPO-II | 0.966085854 | 0.033914146 |
| *Chimera*UPO-I | 0.732176863 | 0.267823137 |
| *Chimera*UPO-II | 0.77998743 | 0.22001257 |
| *Chimera*UPO-III | 0.995203265 | 0.004796735 |
| *Chimera*UPO-IV | 0.963810185 | 0.036189815 |

**T8. Regioselective Ratios from screening of aromatics substrate pool. Results for phenylpropane; alcohol and ketone were added up.**

*Table 8: Regioselective ratios from the screening of the aromatics pool. The data was processed as described above.*

| Enzyme | 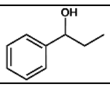 | 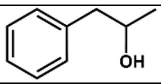 | 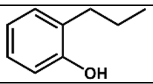 |
|---|---|---|---|
| *Tte*UPO | 0.989936976 | 0.003547883 | 0.006515141 |
| *Dca*UPO | 0.99466572 | 0.001123144 | 0.004211136 |
| *Mro*UPO | 0.57528998 | 0.415915032 | 0.008794989 |
| *Mfe*UPO | 0.99377966 | 0.004056867 | 0.002163473 |
| *Pab*UPO-I | 0.998777694 | 0.001222306 | 0 |
| *Pab*UPO-II | 0.925006848 | 0.074993152 | 0 |
| *Mth*UPO | 0.992770565 | 0.003907886 | 0.003321549 |
| *Aae*UPO* | 0.993677135 | 0.006322865 | 0 |
| *Mhi*UPO | 0.974324508 | 0 | 0.025675492 |
| *Anc*UPO-I | 0.985281084 | 0 | 0.014718916 |
| *Anc*UPO-II | 0.982709388 | 0.002706202 | 0.01458441 |

RESTRICTED

| | | | |
|---|---|---|---|
| *Chimera*UPO-I | 0.993830774 | 0.006122281 | 4.69454E-05 |
| *Chimera*UPO-II | 0.995725194 | 0 | 0.004274806 |
| *Chimera*UPO-III | 0.990080834 | 0 | 0.009919166 |
| *Chimera*UPO-IV | 0.999771901 | 0 | 0.000228099 |

**T9. Regioselective Ratios from screening of aromatics substrate pool. Results for naphthalene; alcohol and ketone were added up.**

*Table 9: Regioselective ratios from the screening of the aromatics pool. The data was processed as described above.*

| Enzyme | 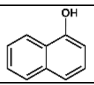 | 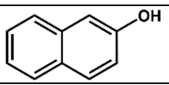 | 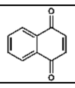 |
|---|---|---|---|
| *Tte*UPO | 0.410828113 | 0.063321385 | 0.525850502 |
| *Dca*UPO | 0.490822319 | 0.040887356 | 0.468290324 |
| *Mro*UPO | 0.415702574 | 0.188165272 | 0.396132153 |
| *Mfe*UPO | 0.552482354 | 0.037352987 | 0.410164659 |
| *Pab*UPO-I | 0.53447563 | 0.085633919 | 0.379890451 |
| *Pab*UPO-II | 0.575803069 | 0.041399773 | 0.382797158 |
| *Mth*UPO | 0.550559431 | 0.066175406 | 0.383265163 |
| *Aae*UPO* | 0.846715177 | 0.037229029 | 0.116055794 |
| *Mhi*UPO | 0 | 0 | 0 |
| *Anc*UPO-I | 0.397302697 | 0.080631125 | 0.522066178 |
| *Anc*UPO-II | 0.386838932 | 0.07944338 | 0.533717688 |
| *Chimera*UPO-I | 0.797564701 | 0.069928643 | 0.132506656 |
| *Chimera*UPO-II | 0.370992047 | 0.139202718 | 0.489805236 |
| *Chimera*UPO-III | 0.710692723 | 0.289307277 | 0 |
| *Chimera*UPO-IV | 0.434423474 | 0.334136483 | 0.231440044 |

**T10. Regioselective Ratios from screening of epoxide substrate pool. Results for 1-hexene; alcohol and ketone were added up.**

*Table 10: Regioselective ratios from the screening of the alkene pool. The data was processed as described above.*

| Enzyme | 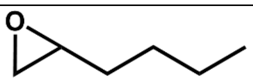 | 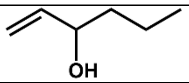 |
|---|---|---|

RESTRICTED

| | | |
|---|---|---|
| *Tte*UPO | 0.950879705 | 0.049120295 |
| *Dca*UPO | 0.9742611 | 0.0257389 |
| *Mro*UPO | 0.752858963 | 0.247141037 |
| *Mfe*UPO | 0.961474411 | 0.038525589 |
| *Pab*UPO-I | 0.831921724 | 0.168078276 |
| *Pab*UPO-II | 0.860595024 | 0.139404976 |
| *Mth*UPO | 0.977022968 | 0.022977032 |
| *Aae*UPO* | 0.72338314 | 0.27661686 |
| *Mhi*UPO | 0.707242228 | 0.292757772 |
| *Anc*UPO-I | 0.934343625 | 0.065656375 |
| *Anc*UPO-II | 0.94362664 | 0.05637336 |
| *Chimera*UPO-I | 0.845878202 | 0.154121798 |
| *Chimera*UPO-II | 0.929989775 | 0.070010225 |
| *Chimera*UPO-III | 0.785492233 | 0.214507767 |
| *Chimera*UPO-IV | 0.652859602 | 0.347140398 |

**T11. Regioselective Ratios from screening of epoxide substrate pool. Results for cyclohexene; alcohol and ketone were added up.**

*Table 11: Regioselective ratios from the screening of the alkene pool. The data was processed as described above.*

| Enzyme | 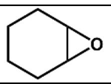 | 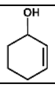 | 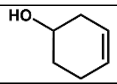 |
|---|---|---|---|
| *Tte*UPO | 0.708059922 | 0.29130776 | 0.000632318 |
| *Dca*UPO | 0.459288635 | 0.540711365 | 0 |
| *Mro*UPO | 0.149425453 | 0.844777726 | 0.005796821 |
| *Mfe*UPO | 0.787848763 | 0.212151237 | 0 |
| *Pab*UPO-I | 0.533745901 | 0.466254099 | 0 |
| *Pab*UPO-II | 0.447624865 | 0.549272909 | 0.003102226 |
| *Mth*UPO | 0.747654548 | 0.252345452 | 0 |
| *Aae*UPO* | 0.33586312 | 0.66413688 | 0 |
| *Mhi*UPO | 0.85649519 | 0.140315432 | 0.003189378 |
| *Anc*UPO-I | 0.45643055 | 0.54356945 | 0 |
| *Anc*UPO-II | 0.534604077 | 0.464160993 | 0.00123493 |
| *Chimera*UPO-I | 0.38950132 | 0.60938214 | 0.00111654 |
| *Chimera*UPO-II | 0.62326881 | 0.373911141 | 0.00282005 |

RESTRICTED

| | | | |
|---|---|---|---|
| *Chimera*UPO-III | 0.113416278 | 0.880315889 | 0.006267833 |
| *Chimera*UPO-IV | 0.062538234 | 0.935148094 | 0.002313672 |

### IV. Chromatograms

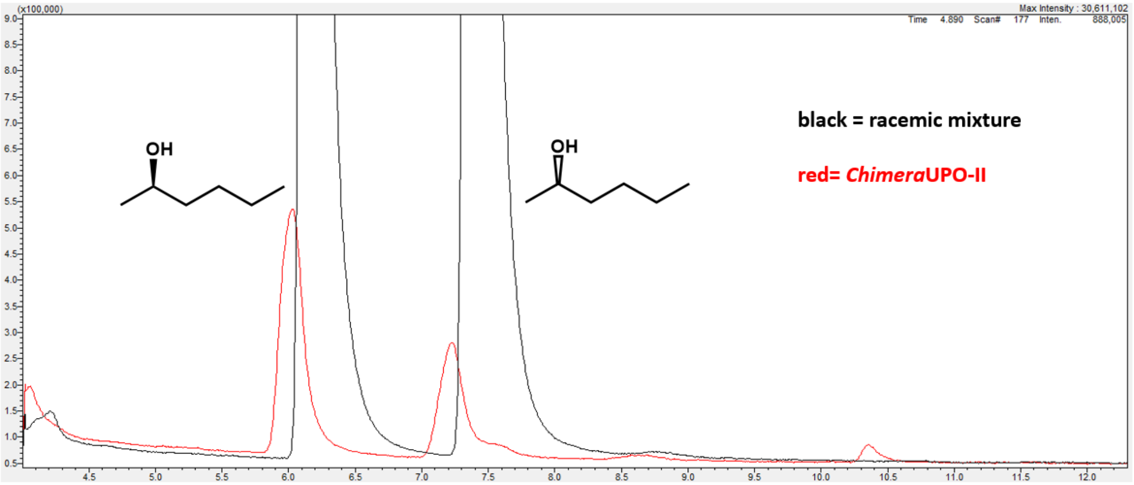


*Figure 49: Chiral measurement of 2-hexanol produced by ChimeraUPO-II in comparison to the racemic mixture.*

RESTRICTED

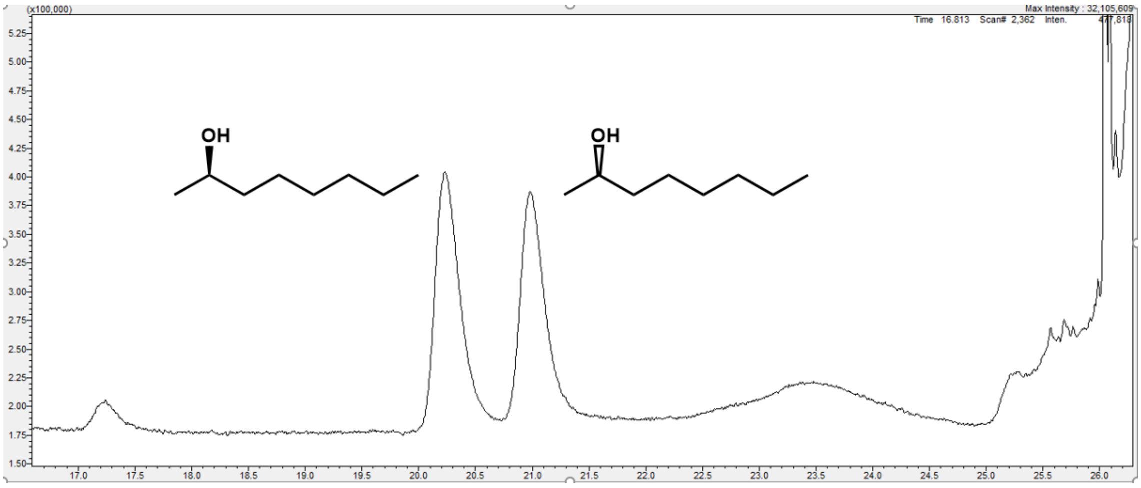


*Figure 50: Chiral measurement of 2-octanol produced by ChimeraUPO-II.*

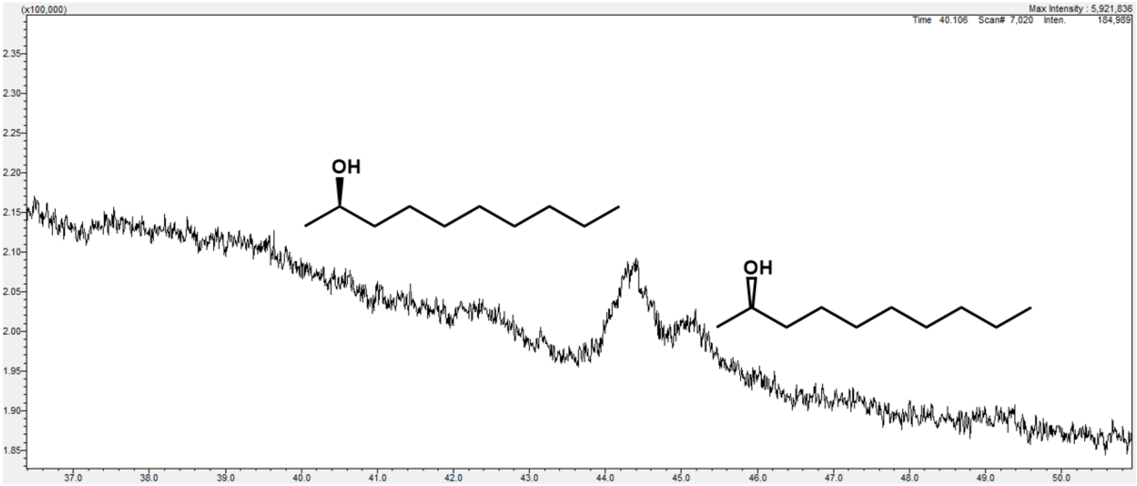


*Figure 51: Chiral measurement of 2-decanol produced by ChimeraUPO-II.*

RESTRICTED

### V. NMR spectra

$^1$H-NMR (400 MHz, $CDCl_3$) δ (ppm) = 7.35 – 7.20 (m, 5 H), 4.52 (m, 1 H), 1.85 – 1.64 (m, 2 H), 0.88 (m, 3 H)

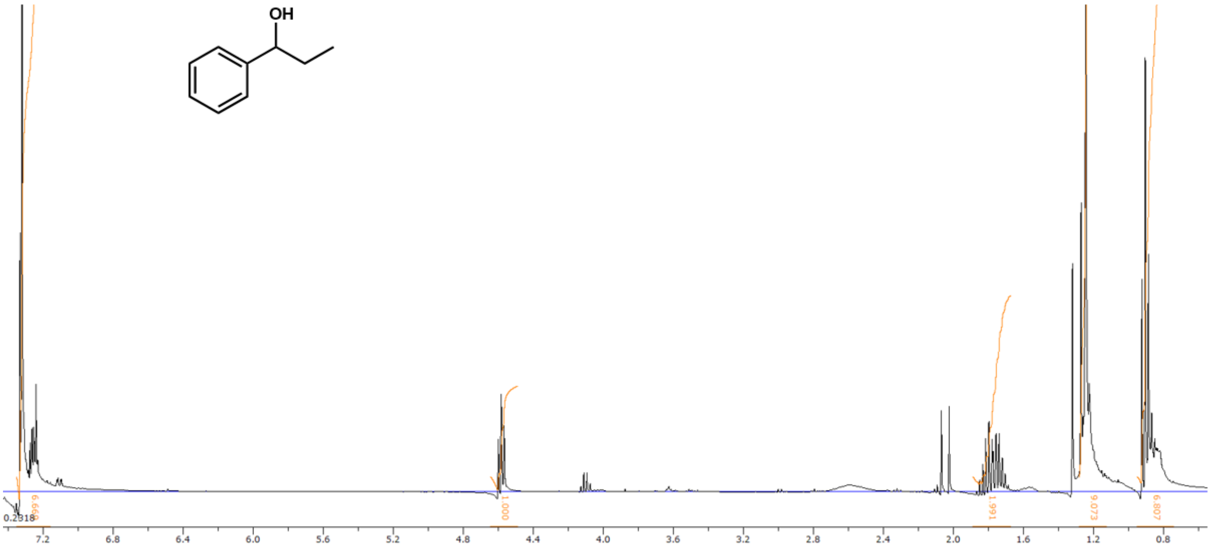


*Figure 52: $^1$H-NMR of 1-phenyl-1-propanol. Impurities are still visible.*

$^1$H-NMR (400 MHz, $CDCl_3$) δ (ppm) = 5.81 (m, 1 H), 5.71 (m, 1 H), 4.17 (s, 1 H), 2.1 – 1.5 (m, 4 H), 1.25 (m, 2 H)

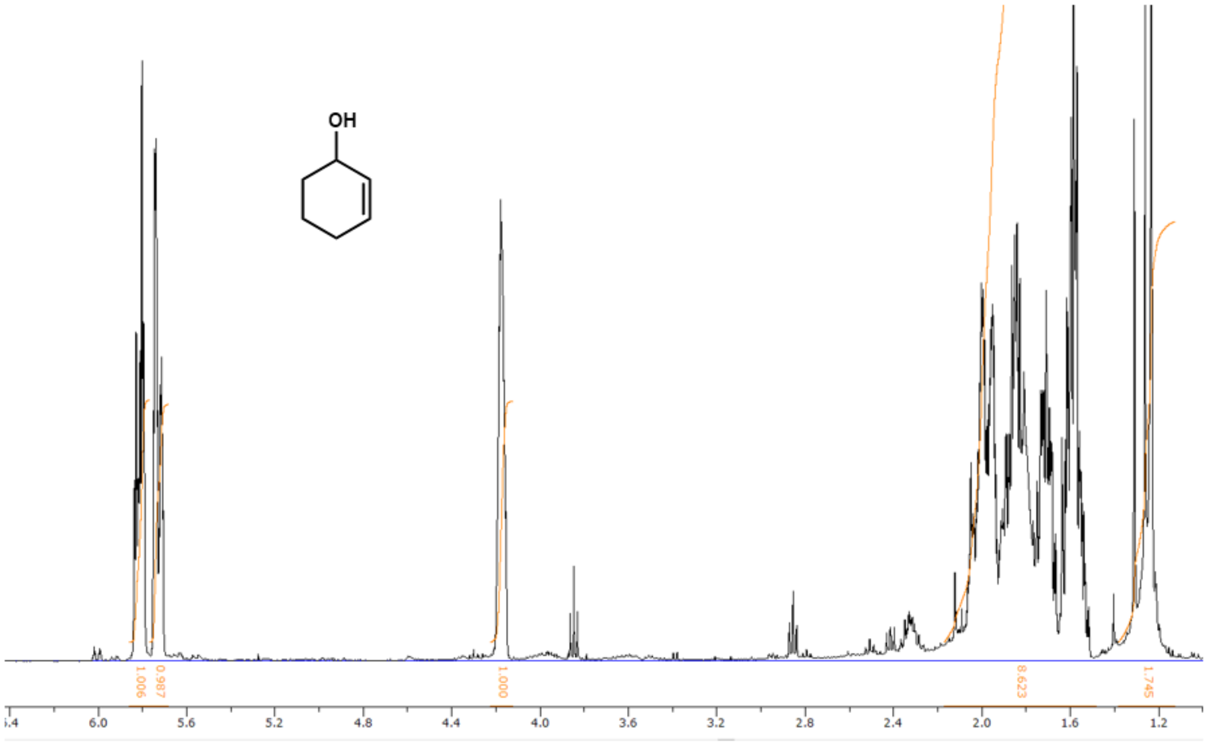


*Figure 53: $^1$H-NMR of 2-cyclohexen-1-ol. Impurities are still visible.*

RESTRICTED

RESTRICTED
